## Supplemental information for "Hierarchical Gaussian process models explore the dark meltome of thermal proteome profiling experiments"

#### Notation

Notations are the same than in the main text and we restate Table 1 of the main text here for convenience:

**Table 1. Notation**

| Notation | Definition |
|---|---|
| $p \in [\![1, P]\!]$ | Set of proteins |
| $c \in [\![1, C_p]\!]$ | Set of conditions for protein $p$ |
| $r \in [\![1, R_{pc}]\!]$ | Set of replicates for condition $c$ of protein $p$ |
| $\pi_j$ with $j \in [\![1, \Pi_p]\!]$ | Set of observed peptides for protein $p$ |
| $N$ | Number of temperatures in the TPP-TR experiment |
| $N_{pcr} \leq N$ | Number of temperatures at which replicate $r$ of condition $c$ of protein $p$ has been observed |
| $N_{pc} = \sum_r N_{pcr}$ | Total dimension of the observations for condition $c$ of protein $p$ |
| $N_p = \sum_c N_{pc}$ | Total dimension of the observations for protein $p$ |
| $T = [t_1, ..., t_N]^T$ | Set of temperatures measured during the TPP-TR experiment |
| $T_{pcr} = [t_1^{pcr}, ..., t_{N_{pcr}}^{pcr}]^T \subseteq T$ | Set of temperatures at which replicate $r$ of condition $c$ of protein $p$ has been observed |
| $T_{pc} = [(T_{pc1})^T, ..., (T_{pcR_{pc}})^T]^T \in \mathcal{R}^{N_{pc}}$ | Concatenation of all observed temperatures for all replicates in conditions $c$ of protein $p$ |
| $T_p = [(T_{p1})^T, ..., (T_{pC_p})^T]^T \in \mathcal{R}^{N_p}$ | Concatenation of all observed temperatures for all conditions of protein $p$ |
| $\Gamma_{pcr} = [\gamma_1^{pcr}, ..., \gamma_{N_{pcr}}^{pcr}]^T$ | Measured raw abundance for replicate $r$ of condition $c$ of protein $p$ |
| $\rho_{pcr}$ | Scaling factor for replicate $r$ of condition $c$ of protein $p$ |
| $y_i^{pcr} = \frac{\gamma_i^{pcr}}{\rho_{pcr}}$ | Scaled raw abundance at $t_i$ for replicate $r$ of condition $c$ of protein $p$ |
| $Y_{pcr} = [y_1^{pcr}, ..., y_{N_{pcr}}^{pcr}]^T$ | Scaled raw abundances for replicate $r$ of condition $c$ of protein $p$ |
| $Y_{pc} = [(Y_{pc1})^T, ..., (Y_{pcR_{pc}})^T]^T \in \mathcal{R}^{N_{pc}}$ | Concatenation of all raw abundances of all replicates of condition $c$ of protein $p$ |
| $Y_p = [(Y_{p1})^T, ..., (Y_{pC_p})^T]^T \in \mathcal{R}^{N_p}$ | Concatenation of all raw abundances of all conditions of protein $p$ |
| $k_\kappa(t, t'\|\lambda) = \sigma_\kappa^2 \cdot k(t, t'\|\lambda)$<br>$k(t, t'\|\lambda) = \exp(-\frac{\|\|t-t'\|\|^2}{2\lambda^2})$ | Radial basis function kernel |
| $\theta_p$ | Set of all parameters and hyper-parameters of the hierarchical Gaussian process model for protein $p$ |
| $\theta_p^{MLE-II}$ | Type II maximum likelihood estimate of the set of parameters and hyper-parameters of the hierarchical Gaussian process model for protein $p$ |

### S1 Appendix: Bayesian model inference and selection

This section aims to clarify the similarities and differences for the models inference and selection between the Bayesian semi-parametric model [16] and the presented model. The content of this section is an adaptation of chapter 5 from Williams and Rasmussen [22].

To illustrate the explanation, we focus on a simple GP regression model, which aims to explain the observations $(T, Y) \in \mathcal{R}^{N\times 2}$:

$$\forall \; i \in [\![1, N]\!] \;\; \begin{cases} y_i = f(t_i) + \epsilon_i \\ f(\cdot) \sim GP(0, k(\cdot,\cdot)) \\ \epsilon_i \overset{iid}{\sim} \mathcal{N}(0, \beta^2) \end{cases} \tag{1}$$

Without loss of generality, the mean of the GP prior has been set to 0, and $k$ is defined as being the squared-exponential kernel, also called Radial Basis Function, or RBF kernel:

$$k(t_i, t_j) = \sigma^2 \cdot \exp(-\frac{||t_i - t_j||^2}{2\lambda^2}) \; . \tag{2}$$

This model has 3 *parameters*, denoted by $\theta = \{\sigma, \lambda, \beta\}$, with $\sigma$ and $\lambda$ often referred to as *hyper-parameters* of the GP prior.

A GP regression is a Bayesian approach because a prior is defined over $f$, which describes a priori believes on $f$. As such, Bayesian inference can be used to estimate the regression's parameters and their posterior distributions, thus giving uncertainty quantification to the parameters estimates.

Moreover, in a Bayesian setting, one can see the task of model selection as equivalent to the *frequentist* task of hypothesis testing. Indeed, considering two models $\mathcal{M}_0$ and $\mathcal{M}_1$, respectively corresponding to two hypothesis $\mathcal{H}_0$ and $\mathcal{H}_1$, Bayesian inference provides a tool to select between these two models.

We describe in the following the different steps of Bayesian inference and model selection, along with the way these are performed in Fang et al. [16] vs in GPMelt.

#### Marginal likelihood

Using Bayes rules, the posterior distribution over the parameters is given by:

$$\begin{aligned} p(\theta|Y,T,\mathcal{M}) &= \frac{p(Y|T,\theta,\mathcal{M})p(\theta|\mathcal{M})}{p(Y|T,\mathcal{M})} \\ \text{with } \; p(Y|T,\mathcal{M}) &= \int p(Y|T,\theta,\mathcal{M})p(\theta|\mathcal{M})d\theta \\ \text{and } \; p(Y|T,\theta,\mathcal{M}) &= \int p(Y|f(T))p(f(T)|T)df. \end{aligned} \tag{3}$$

$p(\theta|\mathcal{M})$ is the prior over the hyper-parameters, and $p(Y|T,\theta,\mathcal{M})$ is the marginal likelihood of the observations given the GP model $\mathcal{M}$. The log marginal likelihood can be shown to take the following form:

$$\begin{aligned} \log\, p(Y|T,\theta,\mathcal{M}) &= -\frac{1}{2}Y^T K_y^{-1} Y - \frac{1}{2}log|K_y| - \frac{N}{2}log2\pi \\ \text{with } \; K_y &= k(T,T) + \beta^2 I_N \; . \end{aligned} \tag{4}$$

#### Bayes factor

Once the hyper-parameters posterior distribution $p(\theta|Y,T,\mathcal{M})$ is known (*model inference*), the *model selection* relies on the computation of the quantity $p(\mathcal{M}_i|Y,T)$ for $i \in \{0,1\}$, i.e. the posterior probability of each model, given the observations. Using Bayes rules, this quantity is given by:

$$p(\mathcal{M}_i|Y,T) = \frac{p(Y|T,\mathcal{M}_i)p(\mathcal{M}_i)}{\sum_i p(Y|T,\mathcal{M}_i)p(\mathcal{M}_i)} \ . \tag{5}$$

The posterior probabilities of models $\mathcal{M}_0$ and $\mathcal{M}_1$ are further compared via :

$$\begin{aligned} \frac{p(\mathcal{M}_1|Y,T)}{p(\mathcal{M}_0|Y,T)} &= \frac{p(\mathcal{M}_1)}{p(\mathcal{M}_0)} \cdot \frac{p(Y|T,\mathcal{M}_1)}{p(Y|T,\mathcal{M}_0)} \\ &= \frac{p(\mathcal{M}_1)}{p(\mathcal{M}_0)} \cdot B \ . \end{aligned} \tag{6}$$

with $B$ the Bayes factor [26], and $p(\mathcal{M}_i)$ for $i \in \{0,1\}$ the prior probability of model $i$. However, an important limitation of this inference is linked to the computation of the quantity $p(Y|T,\mathcal{M}_i)$ given in Eq (3), which is often not analytically tractable, and requires approximation methods or estimation via Markov chain Monte Carlo (MCMC) methods [22].
The Bayesian semi-parametric model relies on the Bayes factor to perform model selection, requiring the estimation of the quantities $p(Y|T,\mathcal{M}_i)$ for $i \in 1,2$. To this aim, Fang et al. [16] used the Metropolis-Laplace estimator with Hamiltonian Monte-Carlo sampling to estimate the hyper-parameters $\theta$.

#### Type II Maximum Likelihood Estimation

While the Bayesian semi-parametric model [16] is a fully Bayesian approach, the GPMelt framework lies at the boundary between Bayesian and frequentist statistics. As mentioned above, a GP regression is a Bayesian approach due to the use of GP priors. However, we choose here to proceed to a frequentist inference of the model's hyper-parameters via type II Maximum Likelihood Estimation (type II MLE), and further develop a hypothesis testing framework presented in the main text.
Type II Maximum Likelihood Estimation is a widely used frequentist method to estimate hyper-parameters in GP regression. In type II MLE, the derivatives of the marginal likelihood w.r.t the hyper-parameters are maximised [22]:

$$\begin{aligned} \frac{\partial}{\partial \theta_j} \log\, p(Y|T,\theta,\mathcal{M}_i) &= \frac{1}{2} tr((\alpha\alpha^T - K^{-1})\frac{\partial K}{\partial \theta_j}) \\ &\text{with } \ \alpha = K^{-1}y \\ &\text{and } \ K = k(X,X) \ . \end{aligned}$$

While more efficient than full Bayesian inference in our case, this method also presents limitations. In particular, the marginal likelihood can suffer from multiple local maxima. When the amount of available data is low, choosing between local maxima can be a difficult task. These local maxima can be seen as different interpretations of the data [22], typically 1) a small lengthscale and a little amount of noise versus 2) a large lengthscale and a large noise. In interpretation 1), the model tends to overfit the data due to the small lengthscale offering a large flexibility to the GP. In this case, the model can explain almost all the data and the noise is reduced to a minimum. In situation 2), the large lengthscale constraints the GP to be very rigid and to not fit the data very well, leading to a large noise estimation to explain the data.

**Note to the GPMelt users**: Given that protein-level TPP-TR data might present a small amount of observations per replicate (at most $N$ temperatures, with $N \approx 10$), these data could typically suffer from this multiple local maxima problem. Using a multi-task learning approach as proposed in GPMelt helps to deal with this limitation, via information sharing between replicates. Similarly, very complex models (e.g. peptide-level analysis presented in the main text with up to 21 conditions) can also lead to multiple local optima. We thus recommend the users to always assess the quality of the model on a few IDs first, and to adapt adequately the constraint on the lengthscale(s) and/or noise parameters if the fits seem to fall in one of the two extremes presented above.

### S2 Appendix: Linking the hierarchical model to the multitask framework

The hierarchical Gaussian process (HGP) model implementation using the multi-task Gaussian process regression framework is detailed in this section.
While GPMelt is based on an adaptation of the hierarchical model presented in Hensman et al. [24], we suggest to interpret this model in light of the multi-task GP regression [25] framework, for which an efficient implementation exists in the python package GPyTorch [39]. Indeed, breaking down the complexity of the covariance matrix $\Sigma_p$ in terms of index kernels greatly simplifies the understanding and interpretation of the model. Hence, extension of the model to deeper hierarchies or different experimental setups is considerably facilitated.
Multi-task learning is a field of Machine Learning that aims to share information among related tasks to improve individual task learning and predictions compared to individual and independent tasks learning [25]. Similarly, the three-level hierarchical model described in the main text and given hereafter by Eq (8), improves the modelling of each replicate in each condition by sharing information between all replicates of all conditions for each protein. A *task* of the multi-task learning framework thus corresponds to a *replicate* in our hierarchical model framework. We restate the three-level hierarchical model for clarity:

$$\begin{matrix} \forall \ i \in [\![1, N]\!] \\ \forall \ r \in [\![1, R]\!] \\ \forall \ c \in [\![1, C]\!] \end{matrix} \begin{cases} h \sim GP(0, k_h(t, \cdot|\lambda_1)) & \textit{protein} \\ g_c \sim GP(h, k_g(t, \cdot|\lambda_1)) & \textit{conditions} \\ f_{cr} \sim GP(g_c, k_{f_{cr}}(t, \cdot|\lambda_2)) & \textit{replicates} \\ y_{cri} = f_{cr}(t_i) + \epsilon_{cri} & \textit{observations} \\ \epsilon_{cri} \overset{iid}{\sim} \mathcal{N}(0, \beta^2) \end{cases} \tag{8}$$

Thanks to the linearity of the hierarchical model [24], the likelihood of model (8) for protein $p$ can be rewritten as:

$$Y_p|T_p, \theta_p \sim \mathcal{N}(\mathbf{0}, \Sigma_p + \beta_p^2 I_{N_p}) \tag{9}$$

where $Y_p \in \mathcal{R}^{N_p}$ is the concatenation of all observations of all replicates in all conditions, $T_p \in \mathcal{R}^{N_p}$ the concatenation of all observed temperatures for all replicates in all conditions, $\theta_p$ the set of all parameters and hyper-parameters to be estimated, $\mathbf{0} \in \mathcal{R}^{N_p}$ the null vector, $I_{N_p} \in \mathcal{R}^{N_p \times N_p}$ the identity matrix and $\Sigma_p \in \mathcal{R}^{N_p \times N_p}$.

In this section, we show how we propose to rewrite the covariance matrix $\Sigma_p$ from Eq (9) as a special matrix product of an index kernel $K^y$ and a correlation matrix $K^{t,\lambda}$. We start by introducing the Hadamard and Kronecker products as two special matrix products.

#### Hadamard product

The Hadamard product is an element-wise product between matrices of the same dimensions [40]. We illustrate this product considering the block matrices $A$ and $B$, for which $A_{ij}$ and $B_{ij}$ are of the same dimensions for all $i, j \in [\![1, N]\!]$:

$$\begin{bmatrix} A_{11} & \cdots & A_{1N} \\ \vdots & & \vdots \\ A_{N1} & \cdots & A_{NN} \end{bmatrix} \circ \begin{bmatrix} B_{11} & \cdots & B_{1N} \\ \vdots & & \vdots \\ B_{N1} & \cdots & B_{NN} \end{bmatrix} = \begin{bmatrix} A_{11}B_{11} & \cdots & A_{1N}B_{1N} \\ \vdots & & \vdots \\ A_{N1}B_{N1} & \cdots & A_{NN}B_{NN} \end{bmatrix} \tag{10}$$

#### Kronecker product

The Kronecker product [41] between a matrix $A \in \mathcal{R}^{a \times b}$ and a matrix $B \in \mathcal{R}^{c \times d}$ is a $ca \times db$ block matrix given by:

$$A \otimes B = \begin{bmatrix} a_{11}B & \cdots & a_{1b}B \\ \vdots & & \vdots \\ a_{a1}B & \cdots & a_{ab}B \end{bmatrix} \tag{11}$$

#### General formula

We start by deriving the general formula for the covariance matrix $\Sigma_p$ of Eq (9), and present special cases in the subsequent sections.
The matrix $\Sigma_p$ is a symmetric block matrix, whose blocks $\Sigma_p[cr, c'r'] \in \mathcal{R}^{N_{pcr} \times N_{pc'r'}}$ are given by:

$$\begin{cases} K_h(T_{pcr}, T_{pcr}|\lambda_1) + K_g(T_{pcr}, T_{pcr}|\lambda_1) + & K_{f_{cr}}(T_{pcr}, T_{pcr}|\lambda_2) \\ & if \;\; r = r' \;\; and \;\; c = c' \\ K_h(T_{pcr}, T_{pcr'}|\lambda_1) + K_g(T_{pcr}, T_{pcr'}|\lambda_1) & if \;\; r \neq r' \;\; and \;\; c = c' \\ K_h(T_{pcr}, T_{pc'r'}|\lambda_1) & otherwise \end{cases} \tag{12}$$

with $K_\kappa(T, T'|\lambda)$ the covariance matrix obtained by evaluating the kernel $k_\kappa(\cdot, \cdot|\lambda)$ at temperature vectors $T$ and $T'$, with $\kappa \in \{h, g, f_{cr}\}$.
To align with the multi-task Gaussian process regression framework introduced in Bonilla et al. [25], we aim to rewrite $\Sigma_p$ as a special matrix product of an index kernel $K^y$ and a correlation matrix $K^{t,\lambda}$. The matrix $K^{t,\lambda} \in \mathcal{R}^{N_{cr} \times N_{c'r'}}$ is obtained by evaluating the kernel $k(t, t'|\lambda) = exp(-\frac{||t-t'||^2}{2\lambda^2})$ at some temperature vectors $T$ and $T'$. $K^{t,\lambda}$ defines a correlation matrix (instead of a covariance matrix), given that the kernel $k$ has unit variance [25].

To achieve this goal, we introduce in the following subsections distinct simplifying assumptions corresponding to different specifications of model (8).

#### Constraints on the RBF kernels lengthscales

As a first simplifying assumption, we consider the case where

$$\lambda_1 = \lambda_2 \equiv \lambda$$

In this case, blocks $\Sigma_p[cr, c'r']$ of Eq (12) can be rewritten as:

$$\begin{cases} \left(\sigma_h^2 + \sigma_g^2 + \sigma_{f_{cr}}^2\right) \cdot K^{t,\lambda}(T_{pcr}, T_{pcr}) & if \;\; r = r' \;\; and \;\; c = c' \\ \left(\sigma_h^2 + \sigma_g^2\right) \cdot K^{t,\lambda}(T_{pcr}, T_{pcr'}) & if \;\; r \neq r' \;\; and \;\; c = c' \\ \sigma_h^2 \cdot K^{t,\lambda}(T_{pcr}, T_{pc'r'}) & otherwise \end{cases} \tag{13}$$

The subsequent choice of the index kernel $K^y$ and the special matrix product depends on the presence of missing observations and is considered in the next paragraphs.

**Case 1: No missing observations**

In this simplifying case,

$$T_{pcr} \equiv T_{pc'r'} \equiv T \quad \forall \;\; c, c', r, r'$$

The block structure of $\Sigma_p$ described in Eq (13) simplifies to

$$\Sigma_p[cr, c'r'] = \begin{cases} \left(\sigma_h^2 + \sigma_g^2 + \sigma_{f_{cr}}^2\right) \cdot K^{t,\lambda}(T, T) & if \;\; r = r', c = c' \\ \left(\sigma_h^2 + \sigma_g^2\right) \cdot K^{t,\lambda}(T, T) & if \;\; r \neq r', c = c' \\ \sigma_h^2 \cdot K^{t,\lambda}(T, T) & otherwise \end{cases} \tag{14}$$

We define $K^y \in \mathcal{R}^{\sum_c R_c \times \sum_c R_c}$ by:

$$K^y_{cr,c'r'} = \begin{cases} \left(\sigma_h^2 + \sigma_g^2 + \sigma_{f_{cr}}^2\right) & if \;\; r = r' \;\; and \;\; c = c' \\ \left(\sigma_h^2 + \sigma_g^2\right) & if \;\; r \neq r' \;\; and \;\; c = c' \\ \sigma_h^2 & otherwise \end{cases} \tag{15}$$

with $K^y_{cr,c'r'}$ the entry of the matrix $K^y$ measuring the similarity between replicate $r$ of condition $c$ and replicate $r'$ of condition $c'$.
Thus, the matrix $\Sigma_p$ can be rewritten as a Kronecker product, denoted by $\otimes$, between $K^y$ and $K^{t,\lambda}$ as follows:

$$\Sigma_p = K^y \otimes K^{t,\lambda}(T, T)$$

This product is illustrated in Fig 2B of the main text.

**Case 2: General case**

In the general case of potential missing values, we define the block matrix $K^y$ by :

$$K^y[cr, c'r'] = \mathbb{1}_{cr,c'r'} \cdot \begin{cases} \left(\sigma_h^2 + \sigma_g^2 + \sigma_{f_{cr}}^2\right) & if \;\; r = r' \;\; and \;\; c = c' \\ \left(\sigma_h^2 + \sigma_g^2\right) & if \;\; r \neq r' \;\; and \;\; c = c' \\ \sigma_h^2 & otherwise \end{cases} \tag{16}$$

with $\mathbb{1}_{cr,c'r'}$ the matrix of size $\mathcal{R}^{N_{cr} \times N_{c'r'}}$ filled with ones. $N_{cr}$ represents the number of temperatures at which replicate $r$ of condition $c$ has been observed.
Similarly, we redefine the correlation matrix $K^{t,\lambda}$ as a block matrix given by:

$$K^{t,\lambda}[cr, c'r'] = K^{t,\lambda}(T_{pcr}, T_{pc'r'}) \in \mathcal{R}^{N_{cr} \times N_{c'r'}} \tag{17}$$

The matrix $\Sigma_p$ (Eq (13)) is given by the Hadamard product between the index kernel $K^y$ and the correlation matrix $K^{t,\lambda}$:

$$\begin{aligned} (K^y \circ K^{t,\lambda})[cr, c'r'] &= K^y[cr, c'r'] \cdot K^{t,\lambda}(T_{pcr}, T_{pc'r'}) \\ &= \begin{cases} \left(\sigma_h^2 + \sigma_g^2 + \sigma_{f_{cr}}^2\right) \cdot K^{t,\lambda}(T_{pcr}, T_{pcr}) & if \;\; r = r' \;\; and \;\; c = c' \\ \left(\sigma_h^2 + \sigma_g^2\right) \cdot K^{t,\lambda}(T_{pcr}, T_{pcr'}) & if \;\; r \neq r' \;\; and \;\; c = c' \\ \sigma_h^2 \cdot K^{t,\lambda}(T_{pcr}, T_{pc'r'}) & otherwise \end{cases} \end{aligned} \tag{18}$$

**Note:** Because the Kronecker product is easier to visualize, we illustrate in the main text (Fig 2B) the three-level HGP model applied on protein-level TPP-TR using the simplifying assumption of no missing observations. However, the model implementation involves the Hadamard multi-task GP regression implemented with the python package GPyTorch [39]. Indeed, this formulation is more general and can deal with or without missing observations in the data.

#### Different RBF kernels lengthscales

In the more complex setting of

$$\lambda_1 \neq \lambda_2$$

we can rewrite $\Sigma_p$ as the sum of special matrix products. Indeed, blocks $\Sigma_p[cr, c'r']$ of Eq (12) can be rewritten as:

$$\begin{cases} \left(\sigma_h^2 + \sigma_g^2\right) \cdot K^{t,\lambda_1}(T_{pcr}, T_{pcr}) + \sigma_{f_{cr}}^2 \cdot K^{t,\lambda_2}(T_{pcr}, T_{pcr}) & if \;\; r = r' \;\; and \;\; c = c' \\ \left(\sigma_h^2 + \sigma_g^2\right) \cdot K^{t,\lambda_1}(T_{pcr}, T_{pcr'}) & if \;\; r \neq r' \;\; and \;\; c = c' \\ \sigma_h^2 \cdot K^{t,\lambda_1}(T_{pcr}, T_{pc'r'}) & otherwise \end{cases} \tag{19}$$

In this case, we propose to write $\Sigma_p$ using a sum of Hadamard products:

$$\Sigma_p = (K^{y,\lambda_1} \circ K^{t,\lambda_1}) + (K^{y,\lambda_2} \circ K^{t,\lambda_2}) \tag{20}$$

with $K^{y,\lambda_1}, K^{y,\lambda_2}$ defined by:

$$\begin{aligned} K^{y,\lambda_1}[cr, c'r'] &= \mathbb{1}_{cr,c'r'} \cdot \begin{cases} \left(\sigma_h^2 + \sigma_g^2\right) & if \;\; r \neq r' \;\; and \;\; c = c' \\ \sigma_h^2 & otherwise \end{cases} \\ K^{y,\lambda_2} &= \Sigma_{f_{cr}} \end{aligned} \tag{21}$$

with $\mathbb{1}_{cr,c'r'}$ the matrix of size $\mathcal{R}^{N_{cr} \times N_{c'r'}}$ filled with ones, and $\Sigma_{f_{cr}}$ the block diagonal matrix such that $\forall\, r, c$, $\Sigma_{f_{cr}}[r,r] = \sigma_{f_{cr}}^2 \cdot I_{N_{cr}} \in \mathcal{R}^{N_{cr} \times N_{cr}}$. The block matrices $K^{t,\lambda_1}$ and $K^{t,\lambda_2}$ are defined similarly as in Eq (17).
Likewise, the covariance matrix $\Sigma_p$ of any hierarchical models with more than three levels and two lengthscales can be described as a sum of Hadamard products between appropriate matrices.

### S3 Appendix: GPMelt statistic $\Lambda$ and the case of multiple conditions

##### Intuition for our special choice of statistic

As mentioned in the main text, we propose to use a non-standard statistic, denoted $\Lambda$, for our hypothesis testing framework. More precisely, this statistic is defined as the ratio of the log marginal likelihood of our joint model $\mathcal{M}_0$ and full model $\mathcal{M}_1$:

$$\Lambda_p = -2 \cdot \log \frac{p(Y_p | T_p, \theta_p^{MLE-II}, \mathcal{M}_0)}{p(Y_p | T_p, \theta_p^{MLE-II}, \mathcal{M}_1)} \;, \tag{22}$$

where $\theta_p^{MLE-II}$ are the hyper-parameters estimated via type II MLE for the full model $\mathcal{M}_1$.
This statistic is different from a standard Likelihood Ratio test (LRT) statistic because, for a standard LRT statistic, the full and joint models would have to be fitted

independently by (type II) MLE to estimate the set of parameters and hyper-parameters $\theta^{MLE}_{p,\mathcal{M}_0}$ and $\theta^{MLE}_{p,\mathcal{M}_1}$. Hence the LRT typically takes the form:

$$LRT_p = -2 \cdot \log \frac{p(Y_p|T_p, \theta^{MLE}_{p,\mathcal{M}_0})}{p(Y_p|T_p, \theta^{MLE}_{p,\mathcal{M}_1})} \ . \tag{23}$$

In GPMelt, we propose to only fit the full model $\mathcal{M}_1$ and to directly plug in the estimated parameters and hyper-parameters $\theta^{MLE-II}_{p,\mathcal{M}_1} \equiv \theta^{MLE-II}_{p}$ in the joint model $\mathcal{M}_0$. Hereafter, we explain this choice by linking $\Lambda$ to another statistic introduced by Liu and Barahona [27] and shown to be an appropriate similarity measure for time series modeled by GPs.

**Notation and definitions as introduced by Liu and Barahona [27]** We present hereafter how the framework introduced by Liu and Barahona [27] could be linked to the objective of TPP-TR datasets modelling.
We consider a protein $p$, for which three conditions ($c_1, c_2$ and $c_3$) are measured. We denote by $Y_{c_1}, Y_{c_2}$ and $Y_{c_3}$ the corresponding observations, dropping the index $p$ of the protein for notation simplicity. Without lost of generality, we aim to test if conditions $c_1$ and $c_2$ have significantly different melting behaviours.
Independently of our hierarchical framework, the following full model $\mathcal{M}_1$ could be introduced to describe the melting curves of replicates in the different conditions:

$$\begin{array}{l} \forall \ i \in [\![1, N]\!] \\ \forall \ r \in [\![1, R_c]\!] \\ \forall \ c \in \{c_1, c_2, c_3\} \end{array} \quad \begin{cases} y_{cri} = g_c(t_i) + \epsilon_{cri} \\ g_c(\cdot) \sim GP(0, k(\cdot, \cdot|\lambda)) \\ \epsilon_{cri} \overset{iid}{\sim} \mathcal{N}(0, \beta^2) \ . \end{cases} \tag{24}$$

With the aim of comparing the melting behaviours between conditions $c_1$ and $c_2$, an artificial condition $c_0$ is introduced with the joint model $\mathcal{M}_0$ to describe a joint behaviour between $c_1$ and $c_2$:

$$\begin{array}{l} \forall \ i \in [\![1, N]\!] \\ \forall \ r \in [\![1, R_c]\!] \\ \forall \ c \in \{c_0, c_1, c_2, c_3\} \end{array} \quad \begin{cases} y_{cri} = \begin{cases} g_{c_0}(t_i) + \epsilon_{cri} \ \ if \ c \in \{c_1, c_2\} \\ g_{c_3}(t_i) + \epsilon_{cri} \ \ if \ c = c_3 \end{cases} \\ g_c(\cdot) \sim GP(0, k(\cdot, \cdot|\lambda)) \\ \epsilon_{cri} \overset{iid}{\sim} \mathcal{N}(0, \beta^2) \ . \end{cases} \tag{25}$$

Following the notations and derivations introduced by Liu and Barahona [27], replicate observations from conditions $c_1$ and $c_2$ under the full model $\mathcal{M}_1$ (24) are modeled by two different functions $g_{c_1}$ and $g_{c_2}$ sampled from a same GP given by $GP(0, k(\cdot, \cdot|\lambda))$. Thus, the joint likelihood of $Y_{c_1}$ and $Y_{c_2}$ is given by:

$$p_{\text{diff}}(Y_{c_1}, Y_{c_2}|T_{c_1}, T_{c_2}, \theta_p) = p(Y_{c_1}|T_{c_1}, \theta_p)p(Y_{c_2}|T_{c_2}, \theta_p) \ . \tag{26}$$

On the other hand, under the joint model $\mathcal{M}_0$ (25), replicate observations from conditions $c_1$ and $c_2$ are modeled by a unique function $g_{c_0}$ sampled from $GP(0, k(\cdot, \cdot|\lambda))$. The likelihood is thus given by:

$$p_{\text{same}}(Y_{c_1}, Y_{c_2}|T_{c_1}, T_{c_2}, \theta_p) = p\left( \begin{bmatrix} Y_{c_1} \\ Y_{c_2} \end{bmatrix} \middle| \begin{bmatrix} T_{c_1} \\ T_{c_2} \end{bmatrix}, \theta_p \right) \ . \tag{27}$$

Using this, Liu and Barahona [27] define a new *GP similarity measure* which is itself a likelihood ratio:

$$s(Y_{c_1}, Y_{c_2}) = \log \frac{p_{\text{same}}(Y_{c_1}, Y_{c_2}|T_{c_1}, T_{c_2}, \theta_p)}{p_{\text{diff}}(Y_{c_1}, Y_{c_2}|T_{c_1}, T_{c_2}, \theta_p)} \ . \tag{28}$$

The authors further show that this measure is an appropriate similarity measure for time series modeled by GPs.

**Extension to multiple conditions** The similarity measure $s$ introduced by Liu and Barahona [27] would be especially handful in our multiple conditions setting. We introduce the new variables $N_Z = N_{c_1} + N_{c_2}, T_Z = [T_{c_1}, T_{c_2}] \in \mathcal{R}^{N_Z}$ and $Z = [Y_{c_1}, Y_{c_2}] \in \mathcal{R}^{N_Z}$ and consider the observations of the third condition $c_3$. Under the full model $\mathcal{M}_1$ (24), replicate observations from condition $c_3$ are modeled by a function $g_{c_3}$ different from $g_{c_1}$ and $g_{c_2}$ but sampled from the same GP. Hence, we have:

$$\begin{aligned} p_{\mathcal{M}_1}(Y_{c_1}, Y_{c_2}, Y_{c_3}|T_{c_1}, T_{c_2}, T_{c_3}, \theta_p) &\equiv p(Y_{c_1}, Y_{c_2}, Y_{c_3}|T_{c_1}, T_{c_2}, T_{c_3}, \theta_p, \mathcal{M}_1) \\ &= p_{\text{diff}}(Y_{c_1}, Y_{c_2}, Y_{c_3}|T_{c_1}, T_{c_2}, T_{c_3}, \theta_p) \\ &= p(Y_{c_1}|T_{c_1}, \theta_p)p(Y_{c_2}|T_{c_2}, \theta_p)p(Y_{c_3}|T_{c_3}, \theta_p) \ . \end{aligned} \tag{29}$$

Under the joint model $\mathcal{M}_0$ (25), replicate observations from condition $c_3$ are still modeled by the function $g_{c_3}$. This function is different from $g_{c_0}$, and also sampled from the same GP given by $GP(0, k(\cdot, \cdot|\lambda))$. Therefore,

$$\begin{aligned} p_{\mathcal{M}_0}(Y_{c_1}, Y_{c_2}, Y_{c_3}|T_{c_1}, T_{c_2}, T_{c_3}, \theta_p) &\equiv p(Y_{c_1}, Y_{c_2}, Y_{c_3}|T_{c_1}, T_{c_2}, T_{c_3}, \theta_p, \mathcal{M}_0) \\ &= p_{\text{diff}}(Z, Y_{c_3}|T_Z, T_{c_3}, \theta_p) \\ &= p(Z|T_Z, \theta_p)p(Y_{c_3}|T_{c_3}, \theta_p) \\ &= p_{\text{same}}(Y_{c_1}, Y_{c_2}|T_{c_1}, T_{c_2}, \theta_p)p(Y_{c_3}|T_{c_3}, \theta_p) \ , \end{aligned} \tag{30}$$

where we used that under the null, $p(Z|T_Z, \theta_p) = p_{\text{same}}(Y_{c_1}, Y_{c_2}|T_{c_1}, T_{c_2}, \theta_p)$ as given in Eq (27).
Consequently, the similarity measure $s$ (Eq (28)) between $Y_{c_1}$ and $Y_{c_2}$ reduces to:

$$\begin{aligned} s(Y_{c_1}, Y_{c_2}) &= \log \frac{p_{\mathcal{M}_0}(Y_{c_1}, Y_{c_2}, Y_{c_3}|T_{c_1}, T_{c_2}, T_{c_3}, \theta_p)}{p_{\mathcal{M}_1}(Y_{c_1}, Y_{c_2}, Y_{c_3}|T_{c_1}, T_{c_2}, T_{c_3}, \theta_p)} \\ &= \log \frac{p_{\text{same}}(Y_{c_1}, Y_{c_2}|T_{c_1}, T_{c_2}, \theta_p)p(Y_{c_3}|T_{c_3}, \theta_p)}{p(Y_{c_1}|T_{c_1}, \theta_p)p(Y_{c_2}|T_{c_2}, \theta_p)p(Y_{c_3}|T_{c_3}, \theta_p)} \\ &= \log \frac{p(Y_{c_1}, Y_{c_2}|T_{c_1}, T_{c_2}, \theta_p)}{p(Y_{c_1}|T_{c_1}, \theta_p)p(Y_{c_2}|T_{c_2}, \theta_p)} \end{aligned} \tag{31}$$

Importantly, this expression of $s$ only depends on the observations from the conditions being compared ($Y_{c_1}$ and $Y_{c_2}$), and are independent from the other conditions (e.g. here of $Y_{c_3}$). This is especially interesting in presence of a large number of conditions.

**Conceptual link to our GPMelt framework** The proposed modelling of replicates via HGP models is a bit more complex than the models inspired from Liu and Barahona [27] that we described in Eq (24) and Eq (25). Especially, for both the three and four-level HGP models (Eqs (8), (52)), each replicate is modeled by a different function $f_{cr}$ (or $\eta_{c\pi_j r}$). We consider the three-level HGP model with fixed output-scale for the last level of the hierarchy (i.e. $\sigma_{f_{cr}} \equiv \sigma_f \quad \forall r, c$) to give the reader an intuition of the link between the similarity measure $s$ proposed by Liu and Barahona [27] (Eq (28)) and the statistic $\Lambda$ used in GPMelt (Eq (22)).
By using the same set of parameters $\theta_p^{MLE-II}$ estimated by fitting $\mathcal{M}_1$ via type II MLE for both the joint and full models, one can notice that:

- Under the full model (Eq (8)), $f_{c_j,r}$ can be seen as *replicate observations* from different functions $g_{c_j}$, with $g_{c_j} \sim GP(h, k_g(t, .|\lambda_1)) \quad \forall j, r$.
- Under the joint model comparing condition $c_1$ and $c_2$ (Eq (34)), $f_{c_j,r}$ can be seen as *replicate observations* from functions $g_{c_0}$ for $j \in \{1, 2\}$, while $f_{c_3,r}$ are *replicate observations* from functions $g_{c_3}$. Both $g_{c_0}$ and $g_{c_3}$ are sampled from the same GP given by $GP(h, k_g(t, .|\lambda_1))$.

However, the hierarchical structure of our model leads to a more complex expression for the marginal likelihood ratio between joint and full models than $s$ (Eq (28)). We derive in the following the simplified expression of the statistic $\Lambda$ in our GPMelt framework.

**Simplified expression of the statistic $\Lambda$ for GPMelt**

Similarly as in the previous paragraphs, we consider a protein $p$, for which three conditions ($c_1, c_2$ and $c_3$) are measured. We denote by $Y_1, Y_2$ and $Y_3$ the corresponding observations, dropping the index $p$ of the protein and index $c$ of the condition for notation simplicity. Without lost of generality, we aim to test if conditions $c_1$ and $c_2$ have significantly different melting behaviours. We define new vectors, $T_Z = [T_1, T_2] \in \mathcal{R}^{N_Z}$ and $Z = [Y_1, Y_2] \in \mathcal{R}^{N_Z}$, and propose to write the likelihood of independent observations $Y_1, Y_2, Y_3$ under a model $\mathcal{M}$ in function of conditional distributions:

$$\begin{aligned} p_{\mathcal{M}}(Y_1, Y_2, Y_3|T_1, T_2, T_3, \theta) &\equiv p(Y_1, Y_2, Y_3|T_1, T_2, T_3, \theta, \mathcal{M}) \\ &= p_{\mathcal{M}}(Z, Y_3|T_Z, T_3, \theta) \\ &= p_{\mathcal{M}}(Z|Y_3, T_Z, T_3, \theta)p_{\mathcal{M}}(Y_3|T_3, \theta) \ . \end{aligned} \tag{32}$$

In this equation, the first term is the conditional probability of $Z$ given $Y_3$, while the second term is the marginal probability of $Y_3$. This expression is very general, and we now suggest to consider the three-level HGP model (Eq (8)) with likelihood given by Eq (9):

$$\begin{aligned} & Y|T, \theta, \mathcal{M}_1 \sim \mathcal{N}(\mathbf{0}, \Sigma^{\mathcal{M}_1}) \\ \text{with} \quad & \Sigma^{\mathcal{M}_1} = \widetilde{\Sigma}^{\mathcal{M}_1} + \beta^2 I_N \ . \end{aligned} \tag{33}$$

The joint model $\mathcal{M}_0$ associated to comparing conditions $c_1$ and $c_2$ is restated hereafter:

$$\begin{array}{c} \forall \ i \in [\![1, N]\!] \\ \forall \ r \in [\![1, R]\!] \end{array} \begin{cases} h \sim GP(0, k_h(t, \cdot|\lambda_1)) & \textit{protein} \\ g_{c_0} \sim GP(h, k_g(t, \cdot|\lambda_1)) & \textit{cond.} c_1 \ \& \ c_2 \\ g_{c_3} \sim GP(h, k_g(t, \cdot|\lambda_1)) & \textit{cond.} c_3 \\ f_{cr} \sim \begin{cases} GP(g_{c_0}, k_{f_{cr}}(t, \cdot|\lambda_2)) \ if \ c \in \{c_1, c_2\} \\ GP(g_{c_3}, k_{f_{cr}}(t, \cdot|\lambda_2)) \ if \ c = c_3 \end{cases} & \textit{replicates} \\ y_{cri} = f_{cr}(t_i) + \epsilon_{cri} & \textit{observations} \\ \epsilon_{cri} \overset{iid}{\sim} \mathcal{N}(0, \beta^2) \end{cases} \tag{34}$$

Similarly to Eq (9), the distribution of the observations follows

$$\begin{aligned} & Y|T, \theta, \mathcal{M}_0 \sim \mathcal{N}(\mathbf{0}, \Sigma^{\mathcal{M}_0}) \\ \text{with} \quad & \Sigma^{\mathcal{M}_0} = \widetilde{\Sigma}^{\mathcal{M}_0} + \beta^2 I_N \ . \end{aligned} \tag{35}$$

$\Sigma^{\mathcal{M}_0}$ and $\Sigma^{\mathcal{M}_1}$ can be expressed as block matrices:

$$\begin{aligned} \Sigma^{\mathcal{M}_1} &= \begin{bmatrix} \alpha_1 & \delta_{12} & \delta_{13} \\ \delta_{21} & \alpha_2 & \delta_{23} \\ \delta_{31} & \delta_{32} & \alpha_3 \end{bmatrix} + \beta^2 I_N \\ \Sigma^{\mathcal{M}_0} &= \begin{bmatrix} \alpha_1 & \nu_{12} & \delta_{13} \\ \nu_{21} & \alpha_2 & \delta_{23} \\ \delta_{31} & \delta_{32} & \alpha_3 \end{bmatrix} + \beta^2 I_N \ , \end{aligned} \tag{36}$$

with $\forall\, i,j \in \{1,2,3\}$:

$$
\begin{gathered}
\alpha_i \in \mathcal{R}^{T_{c_i} \times T_{c_i}} \\
\delta_{i,j}, \nu_{i,j} \in \mathcal{R}^{T_{c_i} \times T_{c_j}} \\
\delta_{j,i} = \delta_{i,j}^T \quad \text{and} \quad \nu_{j,i} = \nu_{i,j}^T \ .
\end{gathered}
\tag{37}
$$

It can be observed that, $\forall\, i \in \{1,2,3\}, \alpha_i$ are covariance matrices between replicates of condition $c_i$. However, $\delta_{i,j}, \nu_{i,j}$ are potentially non-square matrices and correspond to the correlation between observations of conditions $i$ and $j$. We refer the reader to S2 Appendix for details about $\alpha_i, \delta_{i,j}$ and $\nu_{i,j}$.
Subsequently, we propose to rewrite $\Sigma^{\mathcal{M}_1}$ and $\Sigma^{\mathcal{M}_0}$ under the form of different block matrices:

$$
\begin{aligned}
\Sigma^{\mathcal{M}_1} &= \begin{bmatrix} \widetilde{A}^{\mathcal{M}_1} + \beta^2 I_{N_Z} & C^T \\ C & \widetilde{B} + \beta^2 I_{N_3} \end{bmatrix} &= \begin{bmatrix} A^{\mathcal{M}_1} & C^T \\ C & B \end{bmatrix} \\
\Sigma^{\mathcal{M}_0} &= \begin{bmatrix} \widetilde{A}^{\mathcal{M}_0} + \beta^2 I_{N_Z} & C^T \\ C & \widetilde{B} + \beta^2 I_{N_3} \end{bmatrix} &= \begin{bmatrix} A^{\mathcal{M}_0} & C^T \\ C & B \end{bmatrix}
\end{aligned}
\tag{38}
$$

Thus, the following equalities hold:

$$
\begin{aligned}
\widetilde{A}^{\mathcal{M}_1} &= \begin{bmatrix} \alpha_1 & \delta_{21}^T \\ \delta_{21} & \alpha_2 \end{bmatrix} \\
\widetilde{A}^{\mathcal{M}_0} &= \begin{bmatrix} \alpha_1 & \nu_{21}^T \\ \nu_{21} & \alpha_2 \end{bmatrix} \\
C &= \begin{bmatrix} \delta_{31} & \delta_{32} \end{bmatrix} \\
\widetilde{B} &= \begin{bmatrix} \alpha_3 \end{bmatrix}
\end{aligned}
\tag{39}
$$

By definition of GPs, $Z$ and $Y_3$ are joint Gaussian distributed under the joint and full models:

$$
\begin{aligned}
\begin{bmatrix} Z \\ Y_3 \end{bmatrix} &\overset{\mathcal{M}_1}{\sim} \mathcal{N}\left( \begin{bmatrix} 0 \\ 0 \end{bmatrix}, \begin{bmatrix} A^{\mathcal{M}_1} & C^T \\ C & B \end{bmatrix} \right) \\
\begin{bmatrix} Z \\ Y_3 \end{bmatrix} &\overset{\mathcal{M}_0}{\sim} \mathcal{N}\left( \begin{bmatrix} 0 \\ 0 \end{bmatrix}, \begin{bmatrix} A^{\mathcal{M}_0} & C^T \\ C & B \end{bmatrix} \right) \ .
\end{aligned}
\tag{40}
$$

Following Gaussian Identities from Appendix A.2 [22], the marginal distribution of $Y_3$ is given by:

$$
\begin{aligned}
Y_3 &\overset{\mathcal{M}_1}{\sim} \mathcal{N}(0, B) \\
Y_3 &\overset{\mathcal{M}_0}{\sim} \mathcal{N}(0, B) \ .
\end{aligned}
\tag{41}
$$

Moreover, the conditional distributions of $Z$ given $Y_3$ are:

$$
\begin{aligned}
Z &\overset{\mathcal{M}_1}{\sim} \mathcal{N}(CB^{-1}Y_3, A^{\mathcal{M}_1} - CB^{-1}C^T) \\
Z &\overset{\mathcal{M}_0}{\sim} \mathcal{N}(CB^{-1}Y_3, A^{\mathcal{M}_0} - CB^{-1}C^T) \ .
\end{aligned}
\tag{42}
$$

Coming back to Eq (32), we have:

$$
\begin{aligned}
p_{\mathcal{M}_1}(Y_1, Y_2, Y_3 | T_1, T_2, T_3, \theta) &= p_{\mathcal{M}_1}(Z | Y_3, T_Z, T_3, \theta) p_{\mathcal{M}_1}(Y_3 | T_3, \theta) \\
p_{\mathcal{M}_0}(Y_1, Y_2, Y_3 | T_1, T_2, T_3, \theta) &= p_{\mathcal{M}_0}(Z | Y_3, T_Z, T_3, \theta) p_{\mathcal{M}_0}(Y_3 | T_3, \theta) \ ,
\end{aligned}
\tag{43}
$$

leading to the following expression for the statistic $\Lambda$, evaluated in $\theta^{MLE-II}$, the type II maximum likelihood estimator of $\theta$ obtained by fitting the full model $\mathcal{M}_1$:

$$\begin{aligned}\Lambda &= -2 \cdot \log \frac{p_{\mathcal{M}_0}(Y_1, Y_2, Y_3 | T_1, T_2, T_3, \theta^{MLE-II})}{p_{\mathcal{M}_1}(Y_1, Y_2, Y_3 | T_1, T_2, T_3, \theta^{MLE-II})} \\ &= -2 \cdot \log \frac{p_{\mathcal{M}_0}(Z | Y_3, T_Z, T_3, \theta^{MLE-II})}{p_{\mathcal{M}_1}(Z | Y_3, T_Z, T_3, \theta^{MLE-II})} \ .\end{aligned} \tag{44}$$

Thanks to this simplification, the computation of the $\Lambda$ statistic consists in computing the ratio between the likelihood of two conditional multivariate normal of dimension $N_Z$. This is especially advantageous in presence of a large number of conditions, where $N_Z \ll N$.
The log likelihood of model $m$, for $m \in \{\mathcal{M}_0, \mathcal{M}_1\}$, is given by:

$$\begin{aligned}\log p_m(Z | Y_3, T_Z, T_3, \theta^{MLE-II}) = -\frac{1}{2} \,[\,&(Z - CB^{-1}Y_3)^T (A^m - CB^{-1}C^T)^{-1} (Z - CB^{-1}Y_3) \\ &+ \log det(A^m - CB^{-1}C^T) \\ &+ N_Z \log(2\pi) \ .\end{aligned} \tag{45}$$

Denoting $W = Z - CB^{-1}Y_3$, the $\Lambda$ statistic takes the form of:

$$\begin{aligned}\Lambda &= -2 \cdot \log \frac{p_{\mathcal{M}_0}(Z | Y_3, T_Z, T_3, \theta^{MLE-II})}{p_{\mathcal{M}_1}(Z | Y_3, T_Z, T_3, \theta^{MLE-II})} \\ &= W^T [(A^{\mathcal{M}_0} - CB^{-1}C^T)^{-1} - (A^{\mathcal{M}_1} - CB^{-1}C^T)^{-1}] W \\ &+ \log \frac{det(A^{\mathcal{M}_0} - CB^{-1}C^T)}{det(A^{\mathcal{M}_1} - CB^{-1}C^T)}\end{aligned} \tag{46}$$

**Simplification in presence of constraints**

As an *extreme* simplified case, we consider a protein with synchronous observations, and the same number of replicates $R$ in all conditions. Moreover, we assume that the output-scale of the lowest level of the hierarchy has been fixed. In the case of the three-level HGP model (Eq (8)), this means that $\forall c, r \quad \sigma_{f_{cr}} \equiv \sigma_f$.
Under these assumptions, the conditions $c$ are exchangeable along the axis of the matrices $\Sigma^{\mathcal{M}_1}$ and $\Sigma^{\mathcal{M}_0}$, and for any two-by-two comparisons, the matrices $A^{\mathcal{M}_1}, A^{\mathcal{M}_0}, C, B$ are constant. This means that $W_c = Z - CB^{-1}Y_c$ is the only variable to be recomputed across comparisons (with $Z$ the vector of observations corresponding to the two conditions being compared).
If the control condition is fixed, then it is sufficient that treatment conditions present the same number of replicates, with synchronous observations and constant output-scale $\sigma_f$. Fig 3A of the main text illustrates a similar case, but does not assumes fixed output-scales.

### S4 Appendix: Area between the curves

In the results section of the main text, the use of the Area Between the Curves (ABC) is suggested as a measure of the discrepancies between the melting curves. The exact formulation of this ABC is presented in this Appendix.
As a first approximation, the ABC can be computed directly using the observations at hand, without requiring any statistical modeling of the data. Considering a protein $p$, we denote the observations of replicate $r$ of condition $c$ by $(T, Y_{rc}) = (\{t_1, ..., t_N\}, \{y_1^{rc}, ..., y_N^{rc}\})$. We use here as simplification assumption that all

replicates have been measured at the same set of temperatures $T$. If this is not the case, $T$ can be taken as the intersection of temperatures at which all replicates of all conditions have been observed. The median observation for condition $c$ is thus given by $(T, \overline{Y^c})$, with $\overline{Y^c}$ the median observation across replicates at each available temperature. The ABC computed from this median principle, denoted by $ABC_{median}$, is given by:

$$ABC_{median} = \sum_{t=t_1}^{t_N} \overline{Y_t^{c_1}} - \overline{Y_t^{c_2}} \ . \tag{47}$$

As a second measure, we suggest to use the output of the HGP model to refine the value of the area between the curves. Especially, we propose to take advantage of the *predictive distribution* computed from the GP regression. Using a similar framework than S1 Appendix, we consider a simple GP regression model, which aims to explain the observations $(T, Y) \in \mathcal{R}^{N \times 2}$:

$$\forall \ i \in [\![1, N]\!] \ \begin{cases} y_i = f(t_i) + \epsilon_i \\ f(\cdot) \sim GP(0, k(\cdot, \cdot)) \\ \epsilon_i \overset{iid}{\sim} \mathcal{N}(0, \beta^2) \end{cases} \tag{48}$$

Without loss of generality, the mean of the GP prior has been set to 0, and $k$ is defined as being an RBF kernel (Eq (2)).
For any set of *test points* $T^*$, the predictive distribution of observations $f^*$ is given by:

$$\begin{aligned} & f^*|T, Y, T^* \sim \mathcal{N}(\overline{f^*}, cov(f^*)) \\ & \quad \text{with} \quad \overline{f^*} \equiv \mathbb{E}[f^*|T, Y, T^*] = K(T^*, T) K_y^{-1} Y \ , \\ & \quad \quad cov(f^*) = K(T^*, T^*) - K(T^*, T) K_y^{-1} K(T, T^*) \\ & \quad \text{and} \quad K_y = K(T, T) + \beta^2 I_N \ . \end{aligned} \tag{49}$$

with $K(T, T)$ the matrix given by $K(T, T)_{i,j} = k(T_i, T_j)$.
$\overline{f^*}$ is called the *posterior mean* of the predictive distribution. We suggest that the posterior mean $\overline{f^{*,c}}$ for condition $c$ can be used as a better approximation of the condition-wise melting behaviour than the median of the observations $\overline{Y^c}$ for this condition. Indeed, by better integrating the observations of other replicates and replicates of other conditions in the model, this posterior mean is more robust to outliers observations that could impact $\overline{Y^c}$, especially if only two replicates per condition are available. Moreover, this posterior mean can be computed for any choice of $T^*$, ensuring a comparison of the melting behaviours on a denser grid of temperatures, rather than a potential poor intersect of observed temperatures. This measure of the Area Between the Curves, denoted $ABC_{GPMelt}$ is given by:

$$ABC_{GPMelt} = \sum_{t=t_1^*}^{t_N^*} \overline{f_t^{*,c_1}} - \overline{f_t^{*,c_2}} \ . \tag{50}$$

The proposed ABC leads to *signed* values, indicating the dominating effect of the change. For example, a positive value denotes that the curve of condition $c_1$ is mainly above the curve of condition $c_2$ over the set of temperatures. However, for cases where there is no dominating effect, but where curves are instead crossing (potentially multiple times), this value of the ABC can tend towards very small values (positive and negative effects compensating each other). Therefore, we propose to compute the absolute ABC instead, measuring the absolute area between the curves. The formula are given hereafter:

$$\begin{aligned} |ABC|_{median} &= \sum_{t=t_1}^{t_N} |\overline{Y_t^{c_1}} - \overline{Y_t^{c_2}}|, \\ |ABC|_{GPMelt} &= \sum_{t=t_1^*}^{t_N^*} |\overline{f_t^{*,c_1}} - \overline{f_t^{*,c_2}}| \ . \end{aligned} \tag{51}$$

### S1 Supporting Information: Introducing constraints on HGP models parameters.

##### Constraints on RBF kernels lengthscales

We introduced the three-level HGP model in main text and restated it the S2 Appendix, model (8). In this model, two different lengthscales $\lambda_1$ and $\lambda_2$ are introduced. Similarly, the four-level HGP model, including three different lengthscales, is introduced in the main text and restated hereafter for simplicity:

$$\begin{array}{l} \forall\ i \in [\![1, N]\!] \\ \forall\ r \in [\![1, R]\!] \\ \forall\ j \in [\![1, \Pi]\!] \\ \forall\ c \in [\![1, C]\!] \end{array} \begin{cases} h \sim GP(0, k_h(t, \cdot|\lambda_1)) & \textit{protein} \\ g_c \sim GP(h, k_g(t, \cdot|\lambda_1)) & \textit{conditions} \\ f_{c\pi_j} \sim GP(g_c, k_{f_{c\pi_j}}(t, \cdot|\lambda_2)) & \textit{peptides} \\ \eta_{c\pi_j r} \sim GP(f_{c\pi_j}, k_{\eta_{c\pi_j r}}(t, \cdot|\lambda_3)) & \textit{replicates} \\ y_{c\pi_j ri} = \eta_{c\pi_j r}(t_i) + \epsilon_{c\pi_j ri} & \textit{observations} \\ \epsilon_{c\pi_j ri} \overset{iid}{\sim} \mathcal{N}(0, \beta^2) & \end{cases} \tag{52}$$

While our hypothesis framework requires the lengthscale of the two first levels (protein and conditions in both cases) to be the same due to the joint model definition, we allow for flexibility in the definition of the lengthscales for the lower levels.

Typically, introducing an independent lengthscale for the lowest level (i.e the *replicates* level) can be used to capture fast variations in the melting curves. This is generally the case for peptide-level melting curves, presenting larger replicate to replicate variability and a larger amount of unconventional melting behaviours. Thus, if $X$ is the number of levels of the HGP model, with the first level corresponding to the protein and the $X^{th}$ level to the replicates, we would recommend to set:

$$\lambda_1 = ... = \lambda_{X-1} \neq \lambda_X \ .$$

In this case, $\lambda_1 = ... = \lambda_{X-1}$ will tend to be large, and capture slow variations, corresponding to the more general trends observed in the data. On the other hand, $\lambda_X$, expected to be smaller, will capture faster variations, better modelling replicate-wise variability.

**Note to the GPMelt**users: we recommend to first consider the results of the simplest model with $\lambda_1 = ... = \lambda_X$, and to only release the constraint on $\lambda_X$ if fast variations in the replicates melting curves are too poorly captured.

##### Constraints on RBF kernels output-scales

For the general statement of the model, we only introduced constraints on the second level (corresponding to the *conditions*) of the three-level (Eq (8)) and the four-level (Eq (52)) HGP models. Indeed, we require the output-scale $\sigma_g^2$ to be the same across conditions, thus allowing to compute the marginal likelihood of the data under the joint

and full models. However, for the three- and four-level HGP models, replicate-specific output-scales, denoted resp. $\sigma^2_{f_{cr}}$ and $\sigma^2_{\eta_{c\pi_j r}}$, are introduced. Moreover, peptide-specific output-scales $\sigma^2_{f_{c\pi_j}}$ are also proposed for the four-level HGP model.
We suggest that replicate-specific output-scales are helpful to better capture greater level of variability between replicates of a same condition. Especially, for the three-level HGP model applied on protein-level TPP-TR datasets, these replicate-specific output-scales can help capturing outlier replicates, containing one to multiple outlier observations. This is further illustrated in the main text, panels A and B of Fig 6. For the four-level HGP model applied on peptide-level TPP-TR datasets, replicate-specific output-scales improves the modelling of replicate to replicate variability, thus better capturing the underlying common trends.

##### Example of constrained model

For the re-analysis of the peptide-level TPP-TR dataset, denoted as phosphoTPP dataset [11], we used GPMelt with a four-level HGP model, considering the following constraints:

$$\lambda_1 = \lambda_2 = \lambda_3 \quad and \quad \sigma_{f_{c\pi_j}} \equiv \sigma_f \;\; \forall\, j, \forall\, c\ .$$

The covariance matrix $\Sigma$ (Eq (9)) resulting from this constrained model is illustrated in S13D Fig.
**Note to the GPMelt users:** Reducing the number of parameters reduces the model complexity and thus the cost of the model estimation. Nevertheless, the release of constraints, providing more flexibility to the model, could be interesting (or even required) depending on the quality of the dataset. In general, a lower quality dataset should preferentially be analysed using a more constrained model, pulling the model away from noise over-fitting.

#### S2 Supporting Information: Hypothesis testing framework and null distribution approximation

##### The choice of the statistic $\Lambda$

We discuss in the main text the use of the statistic $\Lambda$ to evaluates the evidences against the null hypothesis, with the null hypothesis being that the melting curves of the two conditions of interest can be modeled using a shared behaviour. Considering protein $p$, this statistic is defined as the ratio of the log marginal likelihood of the joint model $\mathcal{M}_0$ and the full model $\mathcal{M}_1$, both evaluated considering the hyper-parameters $\theta_p^{MLE-II}$ estimated via type II MLE for the full model $\mathcal{M}_1$. This statistic reads:

$$\Lambda_p = -2 \cdot \log \left( \frac{p(Y_p | T_p, \theta_p^{MLE-II}, \mathcal{M}_0)}{p(Y_p | T_p, \theta_p^{MLE-II}, \mathcal{M}_1)} \right) . \tag{53}$$

Multiple motivations have already been explained in the previous appendices and in the main, and we restate hereafter three main reasons of this choice:

- The *log marginal likelihood* of the data is a key quantity in GP regression (see S1 Appendix) and is readily accessible.
- Using the same set of parameters and hyper-parameters ($\theta_p^{MLE-II}$) for both the joint and full models allows us to derive a link between $\Lambda$ and a statistic shown to

be an appropriate similarity measure for time series modeled by GPs [27] (see S3 Appendix for the derivation).

- We also discussed an additional advantage of this choice, namely the reduction of the global computational cost of GPMelt. This choice of statistic only requires one fitting process per protein, even if multiple conditions are observed for this protein. Indeed, it is enough to fit the full model $\mathcal{M}_1$ to be able to compute the statistic $\Lambda$ for any "two conditions - by - two conditions" comparisons. Moreover, as shown in S3 Appendix with Eq (44), the expression of $\Lambda$ can be simplified as a ratio between the likelihood of two conditional multivariate normal of dimensions $N_Z$, with $N_Z = N_{Control} + N_{Treatment}$. This is especially advantageous in presence of a large number of conditions, where $N_Z \ll N_p$, with $N_p$ the total number of temperatures observed across all replicates and conditions for this protein $p$.

We now discuss an additional reason behind this choice, linked to parameters un-identifiability. We restate hereafter the joint model for a three-level HGP model with two conditions:

$$\begin{array}{l} \forall\ i \in [\![1, N]\!] \\ \forall\ r \in [\![1, R]\!] \\ \forall\ c \in \{c_1, c_2\} \end{array} \begin{cases} h \sim GP(0, k_h(t, \cdot|\lambda_1)) & \textit{protein} \\ g_{c_0} \sim GP(h, k_{g_{c_0}}(t, \cdot|\lambda_1)) & \textit{conditions} \\ f_{cr} \sim GP(g_{c_0}, k_{f_{cr}}(t, \cdot|\lambda_2)) & \textit{replicates} \\ y_{cri} = f_{cr}(t_i) + \epsilon_{cri} & \textit{observations} \\ \epsilon_{cri} \overset{iid}{\sim} \mathcal{N}(0, \beta^2) \end{cases} \tag{54}$$

If this model should be estimated, the parameters $\sigma_h^2$ and $\sigma_{g_{c_0}}^2$ wouldn't be identifiable. Indeed, the marginal prior on $g_{c_0}$ (after integrating out $h$) is given by

$$\begin{aligned} g_{c_0} &\sim GP(0, k_h(t, \cdot|\lambda_1) + k_{g_{c_0}}(t, \cdot|\lambda_1)) \\ &\sim GP(0, k_{h+g_{c_0}}(t, \cdot|\lambda_1)) \\ \text{with} \quad \sigma_{h+g_{c_0}}^2 &= \sigma_h^2 + \sigma_{g_{c_0}}^2 \end{aligned} \tag{55}$$

Hence, there exists an infinite number of valid values for $\sigma_h^2$ and $\sigma_{g_{c_0}}^2$. Instead of fitting this joint model, we suggest to fit the full model given by Eq (8), in which $\sigma_h^2$, $\sigma_g^2$ and $\sigma_{f_{cr}}^2$ are identifiable for all $r, c$. Parameters values estimated for the full model and given by $\theta_p^{MLE-II}$ are then plugged into the joint model to evaluate the log marginal likelihood of the observations under the null hypothesis.

##### $\Lambda$ can take negative values.

To emphasize again that our statistic $\Lambda$ is related to but different from a standard Likelihood Ratio test (LRT) statistic (as discussed in S3 Appendix, and showed with equation (23)), we discuss hereafter the fact that $\Lambda$ can take negative values, while a standard LRT statistic is always positive.
Indeed, a standard LRT statistic compares the goodness of fit of two competing and *nested* models. Nested means that the model $\mathcal{M}_0$ is simpler than model $\mathcal{M}_1$, because it is defined as a *constrained* version of it: some parameters are typically fixed to some values. Consequently $\mathcal{M}_0$ has less free parameters, i.e parameters to be estimated, meaning that $\mathcal{M}_0$ is a simpler model than $\mathcal{M}_1$. $\mathcal{M}_0$ having less free parameters, $p(Y|T, \theta_{\mathcal{M}_1}^{MLE}) \geq p(Y|T, \theta_{\mathcal{M}_0}^{MLE})\ \forall Y, T$ because model $\mathcal{M}_1$, being more complex, can typically fit any data more closely.
The situation is different with the statistic $\Lambda$. Models $\mathcal{M}_0$ and $\mathcal{M}_1$ are not nested, and $\mathcal{M}_0$ has the same number of free parameters as $\mathcal{M}_1$. This means that the complexity of

both models is equivalent. However, $\mathcal{M}_1$ is more *flexible*, because the structure of the covariance matrix, and especially the structure of the index kernel corresponding to the condition level, is less constrained than $\mathcal{M}_0$. Hence, for proteins showing distinct melting behaviours between conditions, we expect a large value of $\Lambda$, because $\mathcal{M}_1$ will better accommodate these observations thanks to its flexibility. On the contrary, if the melting behaviours of the two conditions are similar, both models $\mathcal{M}_0$ and $\mathcal{M}_1$ will be able to fit the data closely. However, the marginal likelihood of the observations under model $\mathcal{M}_0$ will be larger than under $\mathcal{M}_1$. To give the reader some intuition, this can be seen from the expression of the marginal likelihood:

$$p(Y|T,\theta,\mathcal{M}) \quad = \int p(Y|f(T))p(f(T)|T)df \ , \tag{56}$$

where the probability of the observations are integrated over the function space. If the model is more flexible, the function space will be larger, and the individual probability of any function $p(f(T)|T)$ will be lower (the density is flatter).
Hence, for conditions showing very similar melting behaviours, the value of $\Lambda$ is expected to be small, and potentially negative.

##### Approximation of the null distribution

The proposed statistic, which is non-standard, has an unknown distribution under the null hypothesis. Therefore, this unknown distribution, also called null distribution of the statistic, has to be approximated to further determine the statistical significance of large values of $\Lambda$.
To this aim, we suggest to proceed similarly as done in Phillips et al. [28], and to sample dynamics according to the multivariate normal describing the observations distribution under the joint model (Eq (34)). This multivariate normal is specified in Eq (35), with blocks $\widetilde{\Sigma}_p^{\mathcal{M}_0}[cr, c'r']$ given by :

$$\begin{cases} \left(\sigma_h^2+\sigma_g^2\right)\cdot K^{t,\lambda_1}(T_{pcr},T_{pcr})+\sigma_{f_{cr}}^2\cdot & K^{t,\lambda_2}(T_{pcr},T_{pcr}) \\ & if \;\; r=r' \;\; and \;\; c=c' \\ \left(\sigma_h^2+\sigma_g^2\right)\cdot K^{t,\lambda_1}(T_{pcr},T_{pc'r'}) & otherwise \end{cases} \tag{57}$$

We refer the reader to S2 Appendix and S3 Appendix for a more detailed derivation of this block structure.
This sampling method presents two main advantages. First the generated dynamics share the same statistical characteristics than the real data (similar covariance and noise structures), and can notably reproduce the presence of punctual replicates' outliers (see S2 Fig). This permits to estimate a reliable distribution of $\Lambda$ under the null hypothesis, by integrating *imperfect* and *noisy* dynamics, similar to the ones expected in reality. Secondly, generated samples for each protein have the same numbers of observations and replicates than the protein itself. Considering that a lower number of observations would decrease the fitting accuracy and thus impact the value of $\Lambda$, it is important to have null data presenting a similar amount of information than the real data.
Once a sample, i.e. observations simulated under the null hypothesis, has been drawn for a protein $p$, one can fit model (8) to these data to obtain the estimated parameters $\theta_p^{0,MLE-II}$, and compute the corresponding $\Lambda_p^0$ according to Eq (22).
Subsequently, multiple ways of computing the null distribution for $\Lambda$ can be considered and are summarised in S4 Table. The simplest method consists in sampling one to few samples $S_p$ per protein, and combining these samples to form a null dataset of size $S$, from which $S$ values of $\Lambda^0$ are obtained, and used to approximate the null distribution

of $\Lambda$. Nevertheless, some constraints on $S_p$ should be respected:

$$\begin{aligned} &S_p \geq 1 \ \forall \ p \\ &S_p = S'_p \ \forall \ p, p' \\ &S = \sum_p S_p \geq 1000 \ . \end{aligned} \tag{58}$$

By requiring at least one sample of each protein, we ensure that statistical characteristics of all proteins are represented in the null dataset. Forcing all proteins to be sampled at the same frequency guarantees that there will be no bias towards values of $\Lambda$ that could results from specific statistical characteristics of some over-represented proteins. Finally, the size of the null dataset should be maximized. Indeed the null distribution approximation improves as $S$ grows, along with the precision of the computed p-value whose accuracy is by definition bounded by $S$. Thus increasing S also increases the power of the subsequent statistical analysis.
A more sensitive, but also more computationally expensive, way of approximating the null distribution of $\Lambda$ consists in approximating the null distribution per protein, instead of combining samples from all proteins. Therefore, $S_p$ samples (with $S_p$ large, typically $S_p \geq 1000$) should be drawn for each protein $p$. We refer to S4 Table for more details.

##### The choice of the null distribution approximation: an example

In order to understand how to choose the null distribution estimation, it is important to realise that the amount of information entering the model, i.e. the number of observations used for the model fitting, impacts the range of values $\Lambda$ can take. Indeed, more observations, for example through a larger number of included conditions, will generally improve the model fit and the model certainty, thus decreasing the value of $\Lambda$. Consequently, we recommend to use a more precise null distribution estimation (typically per protein or per sub-group of proteins) if a large variability in the number of observations entering the fit is observed among the proteins entering the analysis.
As an example, we consider the analysis of the phospho-TPP dataset [11] presented in the results section *Peptide-level TPP-TR with multiple conditions* of the main text. In this case, proteins could present between one and 664 phospho-peptides, each phospho-peptide observed in three to five replicates. Hence, the total number of observations could greatly vary between proteins. The approach proposed in the main text is to compute the null distribution per protein (method (D) of S4 Table). We show that this approach, denoted hereafter *protein-wise* null distribution approximation, is very sensitive (Fig 5 of the main text), and captures many more hits than the published $T_m$ analysis. However, this method is computationally expensive, and we thus would like to propose a trade-off between null distribution correctness and computational complexity. For this reason, we introduce a *group-wise* null distribution approximation, described as method (E) of S4 Table. In this approach, groups correspond to set of proteins presenting the same number of observed phopho-peptides, thus reducing the variation in number of observations entering the fits between the proteins of a group.
We restate hereafter how GPMelt has been applied to this dataset, for both the *protein-wise* and *group-wise* null distribution approximations. As explained in the results section *Peptide-level TPP-TR with multiple conditions* of the main text, we reduced the maximal number of phospho-peptides per fit to 20, leading to a three-level hierarchical model with at most 21 conditions (counting the control condition being the median over the non-phosphorylated peptides for this gene name). Choosing 20 phospho-peptides as upper limit enables us to capture, thanks to a *unique* fit per protein, any significant phospho-peptides for more than 90% of the proteins in the dataset (1828 out of 1949 proteins). For the remaining 121 proteins with more than 20

phospho-peptides, we randomly split the phospho-peptides in batches, while maximizing the size similarity of the generated batches. For each of these 121 proteins, each generated batch of at most 20 phospho-peptide is called a *pseudo-protein*, and share the same control condition. These pseudo-proteins play the same role as proteins in our dataset, and are fitted independently. Thanks to this split in batches of at most 20 phospho-peptides, the fits of the 664 phospho-peptides observed for protein SRRM2 could be obtained via the fitting of 34 pseudo-proteins (thus representing 20 times less fits than fitting a model per phospho-peptide for this protein).
Moreover, the creation of pseudo-proteins allows to increase the groups sizes (denoted hereafter by $N_g$ for group $g$), with the groups being defined by the set of proteins or pseudo-proteins sharing the same number of phospho-peptides entering the fit. Increasing the group size has the effect of reducing the number of groups with very little number of proteins/pseudo-proteins (i.e it reduces the number of groups with $N_g$ small). This is especially advantageous in the case of the group-wise null distribution approximation and we illustrate below why.
Denoting by $S$ the total number of samples used to approximate the null distribution, we consider a protein/pseudo-protein $p$ and define:

- $C_p$ the number of conditions for this protein/pseudo-protein $p$,
- $\widetilde{S_p}$ the number of samples to be drawn for each condition of this protein/pseudo-protein $p$,
- and $S_p$ the total number of samples to be drawn for this protein/pseudo-protein $p$.

These four quantities are linked as follows:

$$S = \sum_p S_p \quad \text{with} \quad \widetilde{S_p} = \left\lceil \frac{S_p}{C_p} \right\rceil .$$

Consequently, we illustrate how $S_p$ varies between the different null distribution approximations proposed in S4 Table (methods C, D and E), for the reader to better appreciate the differences in computational cost. In the case of the protein-wise approximation (S4 Table, method D), $S_p \equiv S$ samples have to be obtained for each protein/pseudo-protein $p$. Considering the group-wise approximation (S4 Table, method E), the total of $S$ samples is obtained by combining the $S_p$ samples of the $N_g$ proteins/pseudo-proteins constituting group $g$, hence:

$$S_p = \left\lceil \frac{S}{N_g} \right\rceil .$$

Finally, if one would proceed to a dataset-wise approximation (S4 Table, method C, only described here for the purpose of the comparison, likely not appropriate for this dataset as explained above), then the total of $S$ samples is obtained by combining the $S_p$ samples of each of the $N$ proteins/pseudo-proteins constituting the dataset, leading to:

$$S_p = \left\lceil \frac{S}{N} \right\rceil .$$

The reduction in number of samples between the protein- and group-wise approximation is depicted in S19Fig, panels B and C. In this example, the size of the null dataset is $S = 1e4$ for the protein-wise and $S = 3e5$ for the group-wise null distribution approximation. The value of $S$ for the group-wise null distribution approximation has been defined by rounding the total number of samples obtained when combining the

$S = 1e4$ samples of the protein-wise null distribution approximation for the smallest group ($C_p = 18$).
In the following, we propose to compare the results obtained with the protein-wise and group-wise null distribution approximations. Given that the two approaches present distinct sensitivities due to the differences in null distribution approximations, we choose to focus in S19Fig panels D-F, on the top 450 mono-phosphorylated peptides selected by each method (BH adjusted p-value below 0.003 for the *protein-wise* approach, and below 0.008 for the *group-wise* approach). Similarly, in S19Fig panels G-H, we extend the analysis to include the multi-phosphorylated peptides, and thus select the top 650 phospho-peptides selected by each null distribution approximation (BH adjusted p-value below 0.003 for the *protein-wise* approach, and below 0.007 for the *group-wise* approach).
The two approaches do not capture exactly the same set of phospho-peptides, as depicted in S19Fig, panels D and G. We hereafter explain the reasons behind these differences, and how the user can choose the null distribution approximation according to his/her interest. The difference in hits selection doesn't come from differences in peptides ranking within a protein. The top peptides for protein $X$ remains the top peptides for this protein according to both approaches. However, the value of the p-values (and consequently of the adjusted p-values) associated with each peptide of a protein will differ between the two approximations. Indeed, the p-value is a measure of "how extreme" a change is. This sense of "extreme" is relative to what the change is compared to. Conceptually: in the protein-wise approach, changes in melting behaviour are compared to changes that are likely to occur by chance (i.e. under the null) for this protein, whereas in the group-wise approach, changes are compared to changes that are likely to occur by chance for any protein of this group. Thus, small but reproducible changes observed for a peptide of a protein can be extreme compared to all other changes observed for this protein, but not extreme compared to changes observed for other proteins. In the later case, the peptide will be called significant in the protein-wise approach, but not in the group-wise approach. On the other hand, large changes observed for the melting behaviour of a peptide of a protein can be less extreme than changes observed for other peptides of this same protein. In this case, the peptide can be called significant in the group-wise approach, but will be less significantly extreme according to the protein-wise approach.

Thus the protein-wise approach is better at capturing, for each protein, the peptides showing the most extreme changes in melting behaviour compared to the control condition of this protein. On the other hand, the group-wise approach is better at selecting, within a group of proteins, the peptides showing the largest changes in melting behaviours among all peptides showing changes in melting behaviours compared to the control condition. These conclusions are illustrated in S19 Fig. In S19Fig panels E and H, we show that hits uniquely captured by the group-wise approach present significantly higher values of $|ABC|_{GPMelt}$ (the absolute area between the curves, Eqs (51)). This means that the amplitude of the change between the peptide melting curve and the control condition melting curve is larger for phospho-peptides captured by the group-wise approach. On the other hand, S19Fig panel F shows that hits uniquely captured by the protein-wise approach are associated to significantly higher functional scores than hits uniquely captured by the group-wise approach. Hence, the protein-wise approach will better help to detect, for each protein of the dataset, which phosphosites are more likely to be functional, even in presence of small effect size.

It should be noticed that similar conclusions are valid for protein-level TPP-TR datasets. When comparing changes in melting behaviour induced by multiple conditions, the user might favour the protein-wise null distribution approximation (method (D) of S4 Table) to capture, for each protein, the condition(s) (if any) inducing the largest changes in melting behaviour compared to the control condition. On the other hand, the general null distribution approximation (method (C) of S4 Table) will instead capture the set of combinations { condition + protein } showing the largest effect sizes across the entire dataset (i.e all conditions and all proteins taken together).

##### P-values computation

The computation of the statistic $\Lambda$ under the null hypothesis, as explained in the previous paragraphs, gives access to an approximation of the null distribution of $\Lambda$. This null distribution will be used to compute the p-values associated to the comparisons performed in the dataset.
More precisely, considering a null distribution approximation composed of $S$ values of $\Lambda^0$, denoted $\Lambda^0_s \; \forall \, s \in [\![1, S]\!]$, we defined the p-value of protein $p$, denoted $p_p$ and associated to the statistic $\Lambda_p$, analogously to permutation p-values:

$$p_p = \frac{1 + \sum_s \mathbb{1}[\Lambda^0_s \geq \Lambda_p]}{1 + S} \, . \tag{59}$$

with $\mathbb{1}[\Lambda^0_s \geq \Lambda_p] = 1$ if and only if $\Lambda^0_s \geq \Lambda_p$. The addition of 1 on the numerator (and thus in denominator) corresponds to virtually adding the real protein $p$ to the null dataset [42]. It avoids computing null p-values, which could occur if $\Lambda_p$ is actually so extreme that $\mathbb{1}[\Lambda^0_s \geq \Lambda_p] = 0 \, \forall \, s$. This scenario could indeed happen because of the finite size of the null dataset. Adding one is thus mandatory, as strictly positive p-values are required for subsequent multiple testing correction procedures.

#### S3 Supporting Information: Simulation study

Since GPMelt statistic $\Lambda$ is non standard and present unknown properties, we proceed to a simulation study to guarantee the validity of this statistic and of the sampling approach proposed to approximate its null distribution. It should be noted that this simulation study *does not aim* to compare GPMelt with the other methods discussed in the manuscript (i.e. NPARC [6] and the Bayesian semi-parametric model [16]), especially as the generated synthetic data are non-sigmoidal.
Synthetic data were generated according to GPMelt full and joint models, considering hierarchical models with three or four levels, and with two or three conditions. Additionally, we investigated the effect of the proportion of true differential melting curves in the synthetic dataset, varying this number from 1% to 30%. Finally, we examined the impact of noisy data on GPMelt results, and increased the amount of correlated noise per replicate from 1 to 1000 times more than observed in clean real data. We do not expect observations of all proteins in a TPP-TR dataset to be 1000 times noisier than the clean baseline data we used hereafter, but this could happen for a few proteins, hence we consider this scenario to test the performance of GPMelt in extreme conditions. We describe below how the data were generated.

##### Synthetic datasets generation

While it is relatively easy to simulate sigmoidal melting curves with known differences in ABC (e.g. by simply specifying the $\Delta T_m$), it is more complicated to predict differences in ABC for non-sigmoidal data, like the ones GPMelt has been developed to deal with. What can be done instead is to define appropriate parameters for the full hierarchical GP model $\mathcal{M}_1$ (i.e. the model in which each condition is modeled as a separate task in a multi-task framework) and sample from this model. If the parameters are correctly defined, obtained samples will typically present systematic differences between conditions, hence providing a way to simulated *true differential* observations. In the following, "differential observations" refers to (synthetic) IDs presenting different melting curves between conditions. Importantly, using similarly defined parameters together with the joint hierarchical GP model $\mathcal{M}_0$ (i.e. the model in which at least two conditions are modeled as a shared task), one can generate *true negative* observations. To define appropriate parameters, we selected 160 proteins from the ATP 2019 dataset [18] for which the GPMelt statistic $\Lambda$ was found to be among the 200 largest ones, and for which the computed absolute ABC from GPMelt fit ($|ABC|_{GPMelt}$) is among the top 80%. This selection ensures that the real observations for these proteins are really clean (i.e. reproducible replicates inside a condition and large enough difference between conditions). Using these proteins, we considered the associated estimated type II MLE parameters $\theta^{MLE-II}$ as basis to generate synthetic data.

We hereafter describe the general algorithm used to generate synthetic data from a hierarchical GP model with $L$ levels, $C$ conditions and $R$ replicates in total (with $R_c = \frac{R}{C}$ a constant number of replicates per condition). Given such a model, the likelihood of the observations for one protein $p$ can be expressed as:

$$Y_p|T_p, \theta_p \sim \mathcal{N}(\mathbf{0}, \Sigma_p(T_p, T_p) + \beta_p^2 I_{N_p}) \ . \tag{60}$$

Getting rid of the index $p$ for simplicity, we propose to rewrite $\Sigma$ in terms of index kernels and correlation matrix as follows:

$$\Sigma = \left( \sum_{l=1}^{L-1} K^l \right) \circ K^{t,\lambda_1} + K^L \circ K^{t,\lambda_2} \ , \tag{61}$$

with $K^l$ the index kernel of level $l$ with output-scale given by $\sigma_l^2$, $\forall l \in [\![1, L]\!]$. For simplicity, we also assumed that the lengthscales of the correlation matrices are kept fixed for all levels between 1 and $L-1$ ( and set to $\lambda_1$), while the lengthscale associated to the replicate level $L$ is allowed to be different (and set to $\lambda_2$ ).
With these notations, the likelihood can be re-interpreted as follows:

$$\begin{aligned} Y|T, \theta &\sim \mathcal{N}(\mathbf{0}, \left( \sum_{l=1}^{L-1} K^l \right) \circ K^{t,\lambda_1} + \left( K^L \circ K^{t,\lambda_2} + \beta^2 I_N \right)) \\ &\sim \mathcal{N}(\mathbf{0}, \Sigma_m(T,T) + \Sigma_n(T,T)) \ . \end{aligned} \tag{62}$$

$\Sigma_m$ (with $m$ standing for *model*) describes the correlation captured by modelling the data (typically using the protein and conditions levels in a three level HGP model). $\Sigma_n$ (with $n$ standing for *noise*) describes the correlation capturing the noise in the data. This noise itself is composed of a correlated part ($K^L \circ K^{t,\lambda_2}$), which captures biological differences between replicates, and an uncorrelated noise ($\beta_p^2 I_N$), modeling measurement errors.
We now consider the problem of defining parameters for a synthetic hierarchical GP model with $L_s$ levels, $C_s$ conditions and $R_s$ replicates in total (with $R_{s,c} = \frac{R_s}{C_s}$ a constant number of replicates per conditions), based on the parameters estimated for the real protein $p$. The idea is to change the index kernels $K^l$, via the definition of new values for the output-scales $\sigma_l^2$, $\forall l \in [\![1, L]\!]$, while keeping reasonable values for these output-scales. Reasonable values are values which would be biologically plausible. To this aim, we take the following approach: considering one real replicate $r$, one can summarise the total amount of explained variance for this replicate by:

$$\sum_{l=1}^{L-1} \sigma_l^2 + \sigma_{L,r}^2 + \beta^2 \ . \tag{63}$$

Keeping $\lambda_1, \lambda_2$ and $\beta$ unchanged for simplicity, we propose to generate synthetic values of $\{\sigma^2_{l,s}\}_{l=1}^{L-1}$ and $\{\sigma^2_{L_s,r}\}_{r=1}^{R_s}$, such that, overall, the amount of explained variance by the model part and the correlated-noise part is maintained:

$$\begin{aligned} \sum_{l=1}^{L-1} \sigma_l^2 &\equiv \sum_{l=1}^{L_s-1} \sigma^2_{l,s} \\ \sum_{r=1}^{R} \sigma^2_{L,r} &\equiv \sum_{r=1}^{R_s} \sigma^2_{L_s,r} \end{aligned} \tag{64}$$

Considering the type II MLE parameters $\theta_p^{MLE-II}$ estimated for this real protein $p$, we can define two quantities: the average amount of explained variance by one level of the model part $\overline{\sigma_m^2}$, and the average amount of explained variance by one replicate of the correlated-noise part of the model $\overline{\sigma_n^2}$:

$$\begin{aligned} \overline{\sigma_m^2} &= \frac{1}{L-1} \sum_{l=1}^{L-1} \sigma_l^2 \\ \overline{\sigma_n^2} &= \frac{1}{R} \sum_{r=1}^{R} \sigma^2_{L,r} \end{aligned} \tag{65}$$

To be noted, the definition of $\overline{\sigma_n^2}$ allows the output-scale of the lowest level (replicate level) to have a different value for each replicate. This is especially useful to better model replicate to replicate variability and has been typically used in our manuscript.

Using these quantities, we consequently define for the synthetic model the total amount of explained variances by the model part $\sigma^2_{m_s,total}$ and the total amount of explained variance by the correlated-noise part of the model $\sigma^2_{n_s,total}$:

$$\begin{aligned} \sigma^2_{m_s,total} &= \overline{\sigma_m^2} \cdot (L_s - 1) \\ \sigma^2_{n_s,total} &= \overline{\sigma_n^2} \cdot R_s \end{aligned} \tag{66}$$

Finally, we randomly split these total amount of explained variance between the different levels (for $\sigma^2_{m_s,total}$) and replicates ( for $\sigma^2_{n_s,total}$):

$$\begin{aligned} \sigma^2_{m_s,total} &= \sum_{l=1}^{L_s-1} \sigma^2_{l,s} \\ \sigma^2_{n_s,total} &= \sum_{r=1}^{R_s} \sigma^2_{L_s,r} \end{aligned} \tag{67}$$

The last step consist in defining that $\sigma^2_{2,s} = \max_{l \in [\![1, L_s-1]\!]} \sigma^2_{l,s}$. Indeed, we observed that reproducible differences between conditions are obtained if the output-scale corresponding to the conditions (i.e. level two) is larger than the other output-scales for the model part. Considering the interpretation of the output-scales, this choice is natural, as we wish to generate observations for which the largest amount of variations is explained by the condition under the full model. To be noted, this choice of output-scales also ensures cleaner observations generated under the joint model.

#### Variability of the synthetic datasets

To broadly test GPMelt framework, we considered the typical three-level HGP models, but also deeper hierarchies, using the four-level HGP models. Additionally, we investigated the effect of the proportion of true differential melting curves in the synthetic dataset, varying this number from 1% to 30%. Finally, we propose to examine the impact of noise on GPMelt results. Especially, we propose to increase the amount of correlated noise in our dataset using a factor of 1 to 1000. More precisely, this consists in multiplying the total amount of explained variance by the correlated-noise part of the model $\sigma^2_{n_s,total}$ (eq (66)) by this factor before splitting the result between replicates. This models the existence of proteins presenting some strong outlier replicates. Finally, we also explored the effect of fixing (or not) the value of the output-scale of the replicate level, i.e having the same value $\sigma^2_{L_s}$ for all replicates, or allowing for a different value $\sigma^2_{L_s,r}$ for each replicate $r$.

In summary, we tested the following configurations (120 configurations in total, with $N = 1000$ IDs for each configuration and $S = 10$ samples per ID for each scenario).

1. Scenarios : HGP models used to generate and subsequently to fit the synthetic data. Short names in bold are the ones used in figures presenting the results (S3 to S12 Figs).
   - **3Lev - Fixed lev 3 - 2 Cds - 2 reps** : three-level HGP model with two conditions, two replicates for each condition and the replicates' output-scale kept fixed.
   - **3Lev - Free lev 3 - 2 Cds - 2 reps** : three-level HGP model with two conditions, two replicates for each condition and the replicates' output-scale allowed to vary between replicates.
   - **3Lev - Fixed lev 3 - 3 Cds - 2 reps** : three-level HGP model with three conditions, two replicates for each condition and the replicates' output-scale kept fixed.
   - **3Lev - Free lev 3 - 3 Cds - 2 reps** : three-level HGP model with three conditions, two replicates for each condition and the replicates' output-scale allowed to vary between replicates.
   - **4Lev - Fixed lev 4 - 2 Cds - 2 reps** : four-level HGP model with two conditions, two sub-groups, two replicates for each sub-group and the replicates' output-scale kept fixed.
   - **4Lev - Free lev 4 - 2 Cds - 2 reps** : four-level HGP model with two conditions, two sub-groups, two replicates for each sub-group and the replicates' output-scale allowed to vary between replicates.
2. Noise factor: factor used to multiply $\sigma^2_{n_s,total}$ (eq (66)) before being split between replicates:
   - $\{1, 10, 100, 1000\}$
3. Fraction of true differential IDs:
   - $\{0.01, 0.05, 0.1, 0.2, 0.3\}$

#### Results of the simulation study

- We start by depicting examples of 10 samples (five true differential and five true negative IDs) for each scenario with different noise levels in S3 to S6 Figs.
- We then discuss the resulting ROC curves (S7 Fig). The columns depict the different scenarios, the rows the different noise factors. The colors of the curves indicate the fraction of true differential IDs. From this figure, it can be seen that,

up to a factor of 100, GPMelt allows to recover almost perfectly all true differential IDs. The performance of GPMelt only starts to drop for a noise factor of 1000, while remaining pretty good in most scenarios. It can be noted that in this case and for the scenarios with three conditions, a very low amount of differential IDs (e.g. a fraction of 0.01, i.e only 10 true differential IDs out of 1000) can strongly impact the performance of GPMelt. It is also important to account for the fact that using a noise factor of 1000 leads to simulate a real dataset with an extremely high level of noise, as shown in S6 Fig, which depicts some samples obtained at this noise level for the different scenarios.

- Subsequently, we investigate the FDR control by plotting the expected vs observed FDR in S8 Fig. Here also, the columns correspond to the different HGP models' specifications, the rows to the different noise factors and the colors to the fraction of true differential IDs. The black line indicates the $y = x$ line, which represents the ideal case. We see from this figure that GPMelt controls the FDR accurately in most of the cases and up to a noise factor of 100. For a noise factor of 1000, the FDR control can not be guaranteed anymore. It can also be noted that for smaller fractions of true differential IDs (typically 0.1), the observed FDR is generally larger than the expected FDR.
- Finally, we present the Quantile-Quantile plots (QQplots) showing the theoretical quantiles of a uniform $(0, 1)$ distribution vs the empirical quantiles computed from the p-values obtained using GPMelt. S9 to S12 Figs show the QQplots obtained for the different scenarios (columns) and different fractions of true differential IDs (rows). Each plot corresponds to a different noise factor. Points following the black line $y = x$ correspond to p-values uniformly distributed, while points diverging from this line are p-values diverging from the uniform assumption. Well-calibrated p-values are obtained when the negative IDs are associated to p-values uniformly distributed, and hence lying along the $y = x$ line, and the differentials IDs have p-values diverging from the $y = x$ line. As the fraction of differential IDs increases, we expect more points to diverge from the $y = x$ line. To be noted, the empirical computation of the p-values has a precision limited by $S$, the amount of samples in the null distribution approximation, and can't be smaller than

$$\min p = \frac{1}{1+S} \ . \tag{68}$$

  Hence, as the number of differential IDs increases in our synthetic datasets, we expect more p-values to tend towards or equal this minimum. Globally, these QQplots show that the p-values are well calibrated for a noise factor of 1 and 10, and remain mostly acceptable for larger noise factors.

### S4 Supporting Information: Sigmoidal normalisation

Briefly, it consists in selecting a subset of proteins whose melting curves fulfill several criteria defining a good quality thermal denaturation curve. A sigmoidal curve fitted on the median of the melting curves of these proteins is then used as reference to normalize all melting curves towards this behaviour We refer the reader to Savitski et al. [1] for a detailed description of the sigmoidal normalisation procedure.

### S5 Supporting Information: Scaling factor

In the main text, we introduced a new scaling of the observations to replace the broadly used Fold Change scaling. Hereafter, we further detail the advantages and drawbacks of the Fold Change scaling, and illustrate how the mean scaling addresses the main limitations of Fold Change scaling. We then show that the mean scaling is more appropriate than the median or geometric mean scaling in the case of melting curves.

##### The mean scaling as a solution to the limitations of the FC

The Fold Change scaling (FC) of the observations consists in scaling intensities of a replicate by the intensity at the lowest measured temperature:

$$\forall r, c, p \quad \rho_{pcr}^{FC} = \gamma_1^{pcr} \ . \tag{69}$$

FC scaling presents several advantages, and are especially useful for the ease of the melting curves interpretation. Typically, the abundance of a given protein at the lowest temperature was assumed to represent the initial size of the protein's pool in the cell. Thus, as the temperature increases, the fraction of this pool which remains in the folded state decreases, and any point on the melting curve can be interpreted as the fold change in folded protein abundance. Especially, the temperature called the melting point $T_m$ corresponds to a fold change of 0.5, i.e. exactly half of the available proteins have denatured and have been aggregated at this temperature. Moreover, scaling all intensities to force the melting curves to start at one is a simple while powerful way to make observations across replicates more comparable. Especially, when measured in independent Mass Spectrometry (MS) runs, raw abundances of proteins are difficult to compare, as they depend on the experimental conditions in a way which is currently unknown. Finally, fixing the initial scaled intensity to one simplifies the fitting of the sigmoid function $S$, with $S_\infty = \lim_{T \to -\infty} S(T) \equiv 1$ being a fixed parameter.

However, this approach presents some significant drawbacks. First of all, by relying on only one measurement, namely the intensity at the lowest measured temperature, this scaling is highly sensitive to measurement errors, and large measurement errors at the lowest temperature in a replicate propagates to all observations of this replicate. Secondly, fixing all observations to one at the lowest temperature violates the assumption of uniformly normally distributed measurement error $\epsilon_i$ at any temperature $T_i$. Indeed, at $T_{min}$, the variance of the measurement error is actually forced to 0. Finally, and more fundamentally, the biological assumption that the full pool of available proteins in a cell can be measured at the lowest temperature is nowadays challenged by the expanding field of phase transitioning. Phase transitioning is the event during which a soluble protein becomes part of an insoluble assembly, or conversely, an insoluble protein is released from an insoluble assembly and becomes soluble [19, 20]. An example of phase transitioning is phase separation, a biological process by which certain proteins (or a sub-population of these proteins) form membrane-less proteinaceous bodies, in which specific cellular functions can be performed. Especially, combining TPP with solubility proteome profiling (SPP), Sridharan *et al.* [18] hypothesised that some proteins could phase transition upon heating, suggesting that these proteins, initially kept insoluble as part of biomolecular condensates, could be released in the cell at higher temperatures. This further implies that the initial pool of soluble proteins at the lowest measured temperature might not represent the totality of proteins available in the cells, suggesting that the proportion of proteins in the soluble form at the lowest temperature does not have to be assumed to one anymore.

Taking all these considerations into account, we propose to introduce a new scaling of the observations, that we denote by *mean scaling*. It consists in scaling all observations in a replicate by the mean intensity computed across temperatures of this replicate:

$$\forall r, c, p \quad \rho_{pcr}^{mean} = \frac{1}{N} \sum_i \gamma_i^{pcr} \ . \tag{70}$$

Such scaling is more robust to possible measurement errors at any temperature, and allows each replicate curve to start at a different value, thus restoring the possibility of a normally distributed error $\epsilon_{t_1}$ at the lowest temperature. Furthermore, the proposed HGP modelling is more flexible than the previously used sigmoid fitting, and doesn't require the melting curves to exactly start at one, hence relaxing the constraint of using fold changes. Furthermore, HGP models (Eqs (8) and (52)) are mathematically defined such that tasks (lowest level of the hierarchy) modeled as being part of a same group (second last level of the hierarchy) are considered identical up to a scaling factor [39]. Theoretically, the model could thus directly work on raw intensities, concurrently fitting melting curves (i.e tasks) and estimating tasks specific scaling factors, with the scaling factor indicating by how much each replicate melting curve (lowest level of the hierarchy) is *vertically* shifted from the common trend (second last level of the hierarchy). However, in practice, raw intensities measured from different MS runs can have several order of magnitudes differences. For this reason, scaling raw intensities beforehand helps to bring replicates observations to a similar range, which is typically recommended when using optimization algorithms based on gradient descent (the current GPMelt implementation with GPyTorch [39] uses the Adam algorithm [43]). In this way, the optimization algorithm is converging faster towards the minima. Finally, the mean scaling of the observations improves the reproducibility between replicates, as explained in the main text, paragraph "Scaling factor" and shown in Fig6 panels E and F. In the next paragraph, we further justify the choice of the mean scaling compared to the median or geometric mean scaling.

**The mean scaling is more appropriate than the median or geometric mean scaling.**

In this work, we propose to scale the observations in a replicate by the mean intensity of this replicate. However, the mean is not the only summary statistic that could have been considered. Typically, the median intensity could be more robust to very extreme observations. Also, the geometric mean is often used as replacement for the arithmetic mean in time series analysis. In the following, we investigate the use of the median and geometric mean as scaling factors, and illustrate why the mean scaling is better suited for our data type than the median and geometric mean scaling. To this aim, we use the ATP 2019 dataset [18]. The pre-processed data provided by this paper are already scaled using the Fold Change. We show hereafter (see following subsection) that the mean, median and geometric mean scaling of FC data are equivalent to the mean, median and geometric mean scaling of the raw data. For this reason, we can use FC data equivalently to raw data to demonstrate the advantages of the mean scaling over the median and geometric mean scaling.
The median is usually used as a more robust estimator of centrality than the mean, which is more sensitive to extreme values. This is typically true for skewed data distribution, which is not the case of TPP datasets. In S21 Fig panel A, we show that the distribution of the FC values are U-shaped. Hence, because the median is approximately the *middle* value of a replicate, this value will be extremely sensitive to the shape of the melting curve. As illustrated in S21 Fig panel B, if one replicate decreases faster than the other, the median value will be much lower than for the other replicate. Hence, the median of each replicate of a condition might end up being very different, leading to scaled replicates looking quite distinct (see S21 Fig panel C, third facet). More generally, we show in S22 Fig panel A that the spread of computed median values (over all replicates and conditions) is much larger than for the mean values. The mean value is less sensitive to differences in global shape of the melting curves, as it takes into account all values along the curves.
Concerning the geometric mean, the impact of the lowest values of the curves, which

might be very small, leads the geometric mean to be potentially very small compared to the other summary statistic (see S22 Fig panel A). Diving the observations by a small number make the scaled replicates look like *exploding* (see S21 Fig panel C, last facet). In S22 Fig panel B, we show that the absolute difference in scaling factors computed for replicates of a same condition tends to be significantly smaller when using the mean, compared to the median or the geometric mean, as scaling factors (one-sided Wilcoxon rank sum test, p-values $< 2.2e-16$).
Taken together, we have shown that the median scaling is not appropriate in the case of melting curves, as the median of a replicate is too sensitive to the shape of the curve, and particularly to the speed of the melting (corresponding to the slope of the curve when the curve is sigmoidal). Moreover, we have shown that the geometric mean, usually more appropriate than the arithmetic mean for times series data, is too sensitive to very low values, that are typically observed in the melting curves at low temperatures. Comparatively, the mean scaling is more robust to the curve shape and less affected by very small intensity values, hence making this scaling a preferred alternative to the FC scaling.

**The mean, median and geometric mean scaling of FC data are equivalent to the mean, median and geometric mean scaling of the raw data.**

In this subsection, we show that the mean, median and geometric mean scaling of FC data are equivalent to the mean, median and geometric mean scaling of the raw data, justifying the use of the ATP2019 dataset in S21 and S22 Figs.
According to our notations, the scaled intensities are given by:

$$\begin{aligned}
\forall \quad & scaling \in \{FC, mean, median, geomMean\}, \\
& (y_i^{cr})^{scaling} = \frac{\gamma_i^{cr}}{\rho_{cr}^{scaling}} \\
& \text{with} \quad \rho_{cr}^{FC} = \gamma_1^{cr} \\
& \rho_{cr}^{mean} = \frac{1}{N_{cr}} \sum_{i=1}^{N_{cr}} \gamma_i^{cr} \\
& \rho_{cr}^{median} = \begin{cases} \gamma_{\frac{N_{cr}+1}{2}}^{cr} & \text{if } N_{cr} \text{ is odd} \\ \dfrac{\gamma_{\frac{N_{cr}}{2}}^{cr} + \gamma_{\frac{N_{cr}}{2}+1}^{cr}}{2} & \text{if } N_{cr} \text{ is even} \end{cases} \\
& \rho_{cr}^{geomMean} = \exp\left( \frac{1}{N_{cr}} \sum_{i=1}^{N_{cr}} \log \gamma_i^{cr} \right)
\end{aligned} \tag{71}$$

We then denote by $(\rho_{cr}^{scaling})^{FC}$ the scaling computed from the FC instead of the raw intensities, with $scaling \in \{mean, median, geomMean\}$. We have:

$$\begin{aligned}
(\rho_{cr}^{mean})^{FC} &= \frac{1}{N_{cr}} \sum_{i=1}^{N_{cr}} (y_i^{cr})^{FC} \\
&= \frac{1}{N_{cr}} \sum_{i=1}^{N_{cr}} \frac{\gamma_i^{cr}}{\rho_{cr}^{FC}} \\
&= \frac{1}{N_{cr} \cdot \gamma_1^{cr}} \sum_{i=1}^{N_{cr}} \gamma_i^{cr} \\
&= \frac{\rho_{cr}^{mean}}{\gamma_1^{cr}} \ .
\end{aligned} \tag{72}$$

Similarly, we have:

$$(\rho_{cr}^{median})^{FC} = \begin{cases} (y^{cr}_{\frac{N_{cr}+1}{2}})^{FC} & \text{if } N_{cr} \text{ is odd} \\ \dfrac{(y^{cr}_{\frac{N_{cr}}{2}})^{FC} + (y^{cr}_{\frac{N_{cr}}{2}+1})^{FC}}{2} & \text{if } N_{cr} \text{ is even} \end{cases}$$
$$= \frac{\rho_{cr}^{median}}{\gamma_1^{cr}} \ , \tag{73}$$

and

$$\begin{aligned}(\rho_{cr}^{geomMean})^{FC} &= \exp\left(\frac{1}{N_{cr}}\sum_{i=1}^{N_{cr}}\log\left((y_i^{cr})^{FC}\right)\right) \\ &= \exp\left(\frac{1}{N_{cr}}\sum_{i=1}^{N_{cr}}\log\frac{\gamma_i^{cr}}{\rho_{cr}^{FC}}\right) \\ &= \exp\left(\frac{1}{N_{cr}}\sum_{i=1}^{N_{cr}}\left(\log\gamma_i^{cr} - \log\rho_{cr}^{FC}\right)\right) \\ &= \exp\left(\frac{1}{N_{cr}}\sum_{i=1}^{N_{cr}}\log\gamma_i^{cr}\right)\cdot\exp\left(\frac{-1}{N_{cr}}\sum_{i=1}^{N_{cr}}\log\rho_{cr}^{FC}\right) \\ &= \rho_{cr}^{geomMean}\cdot\exp(\log(\frac{1}{\gamma_1^{cr}})) \\ &= \frac{\rho_{cr}^{geomMean}}{\gamma_1^{cr}} \ .\end{aligned} \tag{74}$$

Thus, the scaling of FC data is equivalent to the scaling of the raw data:

$$\begin{aligned}((y_i^{cr})^{FC})^{mean} &= \frac{(y_i^{cr})^{FC}}{(\rho_{cr}^{mean})^{FC}} = \frac{\gamma_i^{cr}}{\gamma_1^{cr}}\cdot\frac{\gamma_1^{cr}}{\rho_{cr}^{mean}} = (y_i^{cr})^{mean} \\ ((y_i^{cr})^{FC})^{median} &= \frac{(y_i^{cr})^{FC}}{(\rho_{cr}^{median})^{FC}} = \frac{\gamma_i^{cr}}{\gamma_1^{cr}}\cdot\frac{\gamma_1^{cr}}{\rho_{cr}^{median}} = (y_i^{cr})^{median} \\ ((y_i^{cr})^{FC})^{geomMean} &= \frac{(y_i^{cr})^{FC}}{(\rho_{cr}^{geomMean})^{FC}} = \frac{\gamma_i^{cr}}{\gamma_1^{cr}}\cdot\frac{\gamma_1^{cr}}{\rho_{cr}^{geomMean}} = (y_i^{cr})^{geomMean}\end{aligned} \tag{75}$$

### Supplementary Figures

#### S1 Fig.

**Summary of GPMelt's algorithmic steps.**

#### S2 Fig.

**Sampling from the hierarchical model allows to generate data with similar statistical properties than the real dataset, including a similar noise level and the presence of outliers.**

#### S3 Fig.

**Example of samples from the synthetic datasets with a noise factor of 1.**

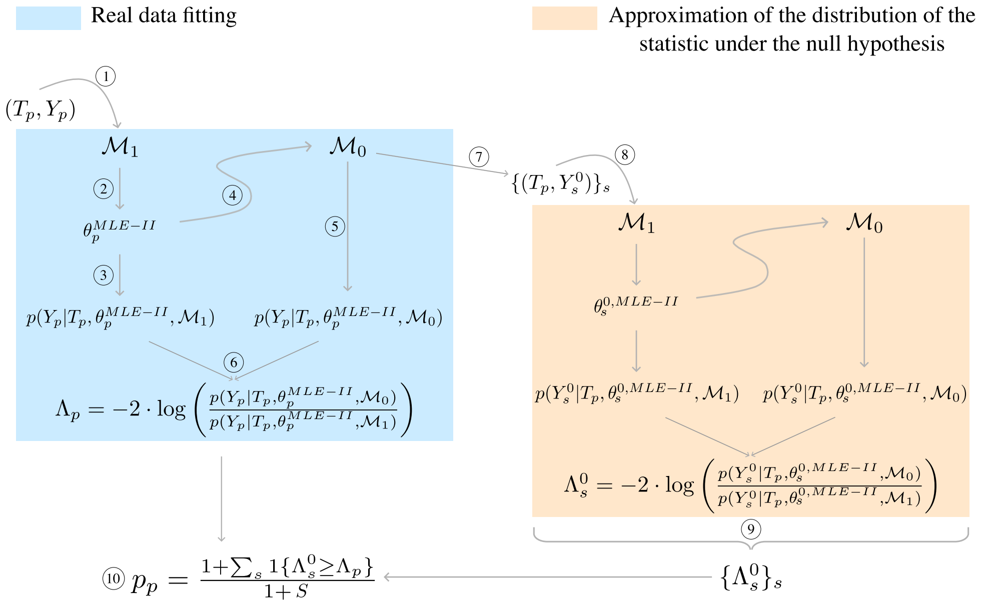


**Fig S1. Summary of GPMelt's algorithmic steps.** (1) The procedure starts by feeding the observed data $(T_p, Y_p)$ of protein $p$ to the full model $\mathcal{M}_1$ for fitting via type II MLE. (2) The obtained parameters estimates are given by $\theta_p^{MLE-II}$. (3) The parameters estimates are used to compute the log marginal likelihood of the observations for this model $\log p(Y_p|T_p, \theta_p^{MLE-II}, \mathcal{M}_1)$. (4) The parameters estimates are then plugged into the joint model $\mathcal{M}_0$ which only differs from the full model $\mathcal{M}_1$ by a change in the covariance matrix structure. (5) The log marginal likelihood of the observations for this joint model $\log p(Y_p|T_p, \theta_p^{MLE-II}, \mathcal{M}_0)$ are computed. (6) The statistic $\Lambda_p$ is defined as the ratio of the previously computed log marginal likelihood. (7) The joint model $\mathcal{M}_0$ with plugged in parameters estimates $\theta_p^{MLE-II}$ is used to draw $S$ samples $\{(T_p, Y_s^0)\}_s$ from the null hypothesis. It can be noticed that the temperatures $T_p$ are the same as for the real data. (8) Each sample $(T_p, Y_s^0)$ then goes to the same steps as described before. Fitting the model $\mathcal{M}_1$ to this sample provides the parameters' type II MLE $\theta_s^{0,MLE-II}$. $\theta_s^{0,MLE-II}$ is used to evaluate the log marginal likelihood of the sample under $\mathcal{M}_1$ and $\mathcal{M}_0$, given that the parameters of $\mathcal{M}_0$ have been fixed to $\theta_s^{0,MLE-II}$. Finally, the statistic $\Lambda_s^0$ is computed for this sample. $\Lambda_s^0$ represents a value of the statistic under the null hypothesis. (9) The statistics of all samples $\{\Lambda_s^0\}_s$ are combined to form the approximation of the distribution of the statistic $\Lambda$ under the null. (10) The p-value associated with $\Lambda_p$ is computed from $\{\Lambda_s^0\}_s$ similarly as a permutation p-value. **Note:** depicted here is the protein-wise null distribution approximation (methods B and D of S4 Table). If the null distribution approximation is obtained by combining samples from all proteins of the dataset (i.e. dataset-wise methods A and C of S4 Table), then the set $\{\Lambda_{ps}^0\}_s$ of values of the statistic under the null obtained for protein $p$ is combined with the sets obtained for all other proteins. The p-value (step 10) is computed using this combined set $\{\Lambda_{ps}^0\}_{p,s}$ instead of $\{\Lambda_s^0\}_s$. Similarly, for the group-wise method E of S4 Table, the p-value of a protein in a group is computed using the combined set $\{\Lambda_{ps}^0\}_{p,s}$ over all proteins $p$ belonging to this group.

### S4 Fig.

**Example of samples from the synthetic datasets with a noise factor of** 10**.**

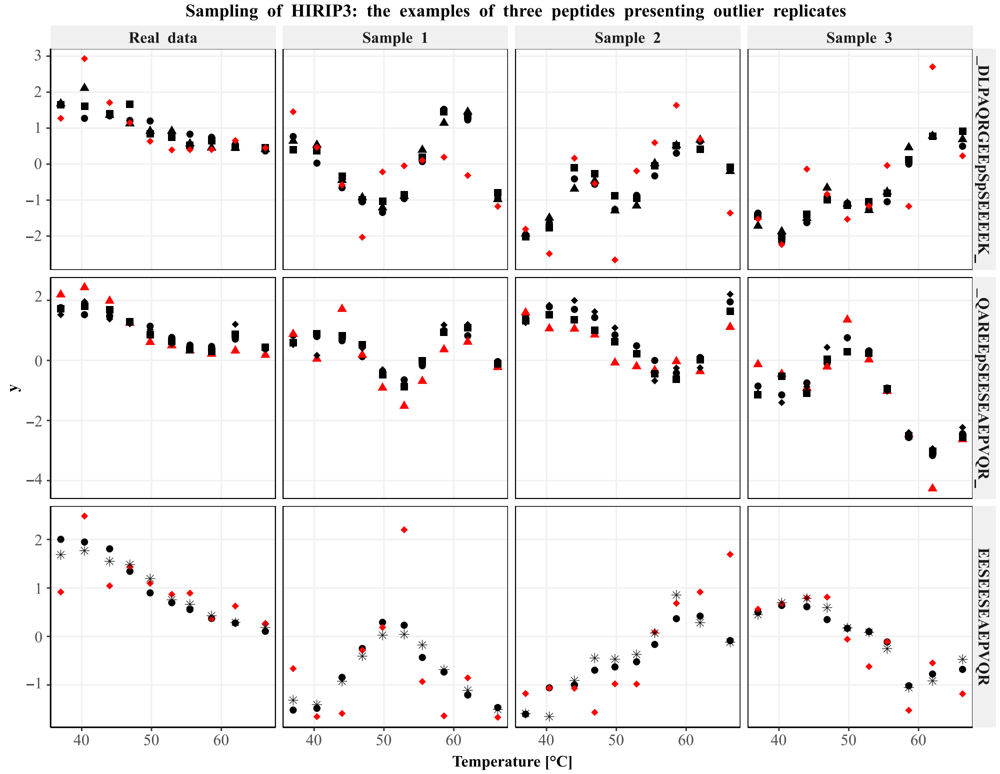


**Fig S2. Sampling from the hierarchical model allows to generate data with similar statistical properties than the real dataset, including a similar noise level and the presence of outliers.** Three peptides (one per row) of protein HIRIP3 from the phosphoTPP dataset [11] have been selected to demonstrate the capacity of the model to generate samples (columns 2 to 4) presenting a similar noise level than the original data (column 1). For each peptide $\pi_j$, a replicate $r$ of the real dataset is highlighted in red. Indeed, a large value for $\sigma^2_{\eta_{\pi_j r}}$ has been estimated from the real data for these replicates, indicating that they were likely to be outlier replicates, or to contain at least one outlier value. This is certainly the case, as can be appreciated from the data visualization (first column). When sampling from a joint HGP model $\mathcal{M}_0$ (here four-levels), the generated data present similar statistical properties than the real data. Especially, replicates presenting a strong covariance in the real data also behave quite similarly in the sampled data (black replicates). On the opposite, replicates which are deviating from the general trend in the real dataset, or for which at least one value is an outlier (red replicates) also present deviating behaviours or outlier values in the sampled data. The amount of deviation for replicate $r$ of peptide $\pi_j$ is controlled by the value of $\sigma^2_{\eta_{\pi_j r}}$ estimated on the real data. Similarly, the amount of similarity between replicates of a peptide $\pi_j$ from condition $c$ is measured by the value of $\sigma^2_{c\pi_j r}$ estimated on the real data.

### S5 Fig.

**Example of samples from the synthetic datasets with a noise factor of** 100**.**

### S6 Fig.

**Example of samples from the synthetic datasets with a noise factor of** 1000**.**

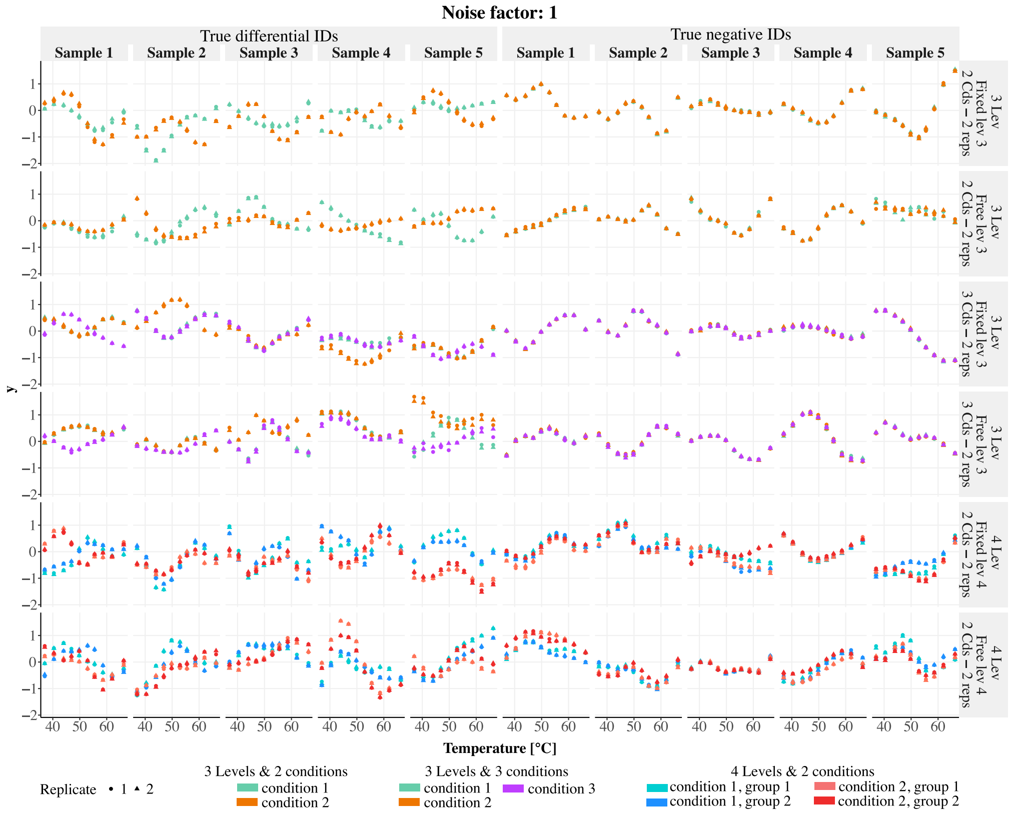


**Fig S3. Random samples coming from the synthetic datasets with a noise factor of 1.** The columns correspond to the different samples, and the rows to different scenarios (i.e HGP model specifications). Five randomly selected true differential samples (obtained under the full model) and five randomly selected true negative samples (obtained under the joint model) are presented for each of the six scenarios. For the three-level HGP models, colors represent the conditions. For the four-level HGP models, colors represent subgroups, with red and orange corresponding to one condition, and blue and light blue corresponding to another condition. For true differential IDs generated using four-level HGP models, replicates coming from the orange and red sub-groups, resp. blue or light blue sub-groups, typically vary similarly but differently from replicates from the other set of color.

#### S7 Fig.

**ROC curves presenting GPMelt performance on the simulated datasets.**

#### S8 Fig.

**Expected vs observed FDR for the simulated datasets.**

#### S9 Fig.

**Quantile-quantile plots of the p-values obtained using GPMelt on the synthetic datasets generated with a noise factor of 1.**

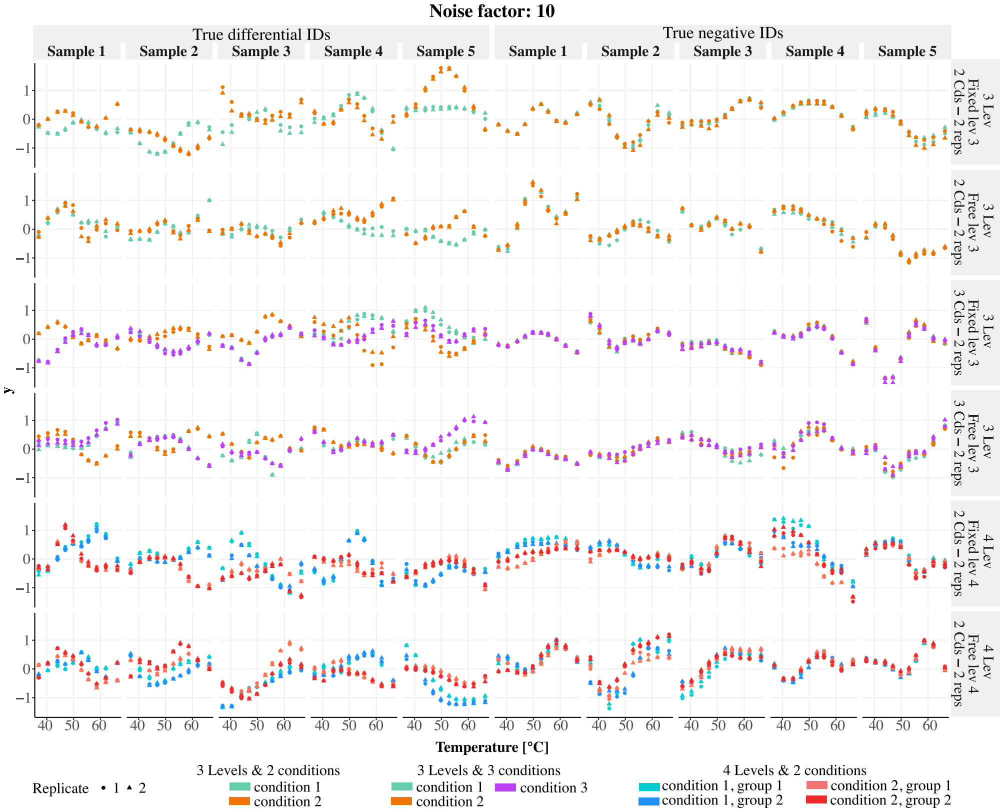


**Fig S4. Random samples coming from the synthetic datasets with a noise factor of 10.** The columns correspond to the different samples, and the rows to different scenarios (i.e HGP model specifications). Five randomly selected true differential samples (obtained under the full model) and five randomly selected true negative samples (obtained under the joint model) are presented for each of the six scenarios. For the three-level HGP models, colors represent the conditions. For the four-level HGP models, colors represent subgroups, with red and orange corresponding to one condition, and blue and light blue corresponding to another condition. For true differential IDs generated using four-level HGP models, replicates coming from the orange and red sub-groups, resp. blue or light blue sub-groups, typically vary similarly but differently from replicates from the other set of color.

### S10 Fig.

**Quantile-quantile plots of the p-values obtained using GPMelt on the synthetic datasets generated with a noise factor of 10.**

### S11 Fig.

**Quantile-quantile plots of the p-values obtained using GPMelt on the synthetic datasets generated with a noise factor of 100.**

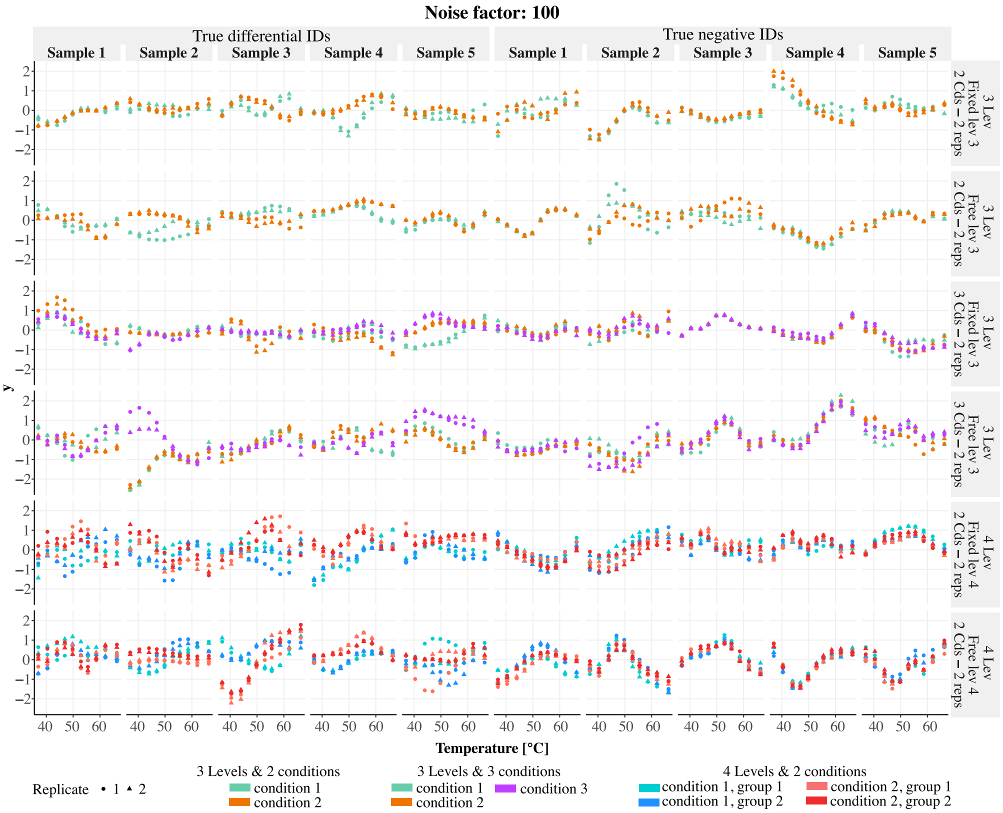


**Fig S5. Random samples coming from the synthetic datasets with a noise factor of** 100**.** The columns correspond to the different samples, and the rows to different scenarios (i.e HGP model specifications). Five randomly selected true differential samples (obtained under the full model) and five randomly selected true negative samples (obtained under the joint model) are presented for each of the six scenarios. For the three-level HGP models, colors represent the conditions. For the four-level HGP models, colors represent subgroups, with red and orange corresponding to one condition, and blue and light blue corresponding to another condition. For true differential IDs generated using four-level HGP models, replicates coming from the orange and red sub-groups, resp. blue or light blue sub-groups, typically vary similarly but differently from replicates from the other set of color.

#### S12 Fig.

**Quantile-quantile plots of the p-values obtained using GPMelt on the synthetic datasets generated with a noise factor of** 1000**.**

#### S13 Fig.

**Extension to deeper hierarchies (schematic description).**

#### S14 Fig.

**Extension to deeper hierarchies and PTMs cross-talk.**

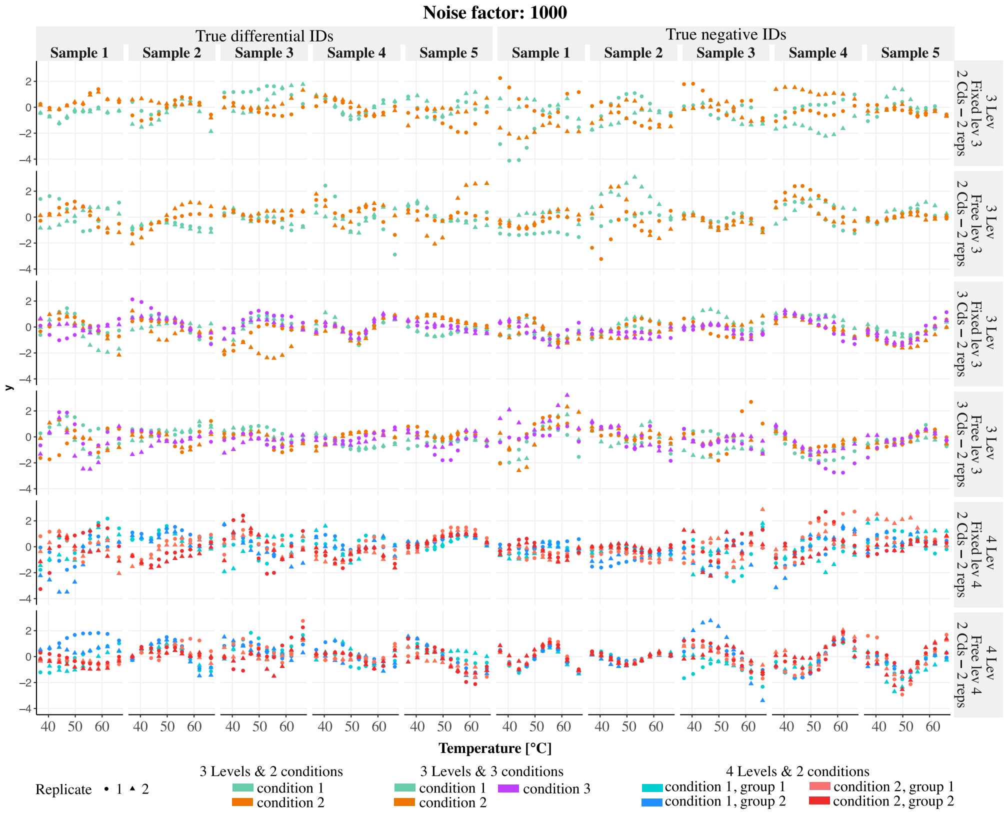


**Fig S6. Random samples coming from the synthetic datasets with a noise factor of** 1000. The columns correspond to the different samples, and the rows to different scenarios (i.e HGP model specifications). Five randomly selected true differential samples (obtained under the full model) and five randomly selected true negative samples (obtained under the joint model) are presented for each of the six scenarios. For the three-level HGP models, colors represent the conditions. For the four-level HGP models, colors represent subgroups, with red and orange corresponding to one condition, and blue and light blue corresponding to another condition. For true differential IDs generated using four-level HGP models, replicates coming from the orange and red sub-groups, resp. blue or light blue sub-groups, typically vary similarly but differently from replicates from the other set of color. With such a high level of noise, some true negative IDs could be seen as potential differential IDs, and reciprocally.

### S15 Fig.

**Additional results for protein-level TPP-TR datasets**

### S16 Fig.

**Additional examples of non sigmoidal curves (ATP 2019 dataset [18])**

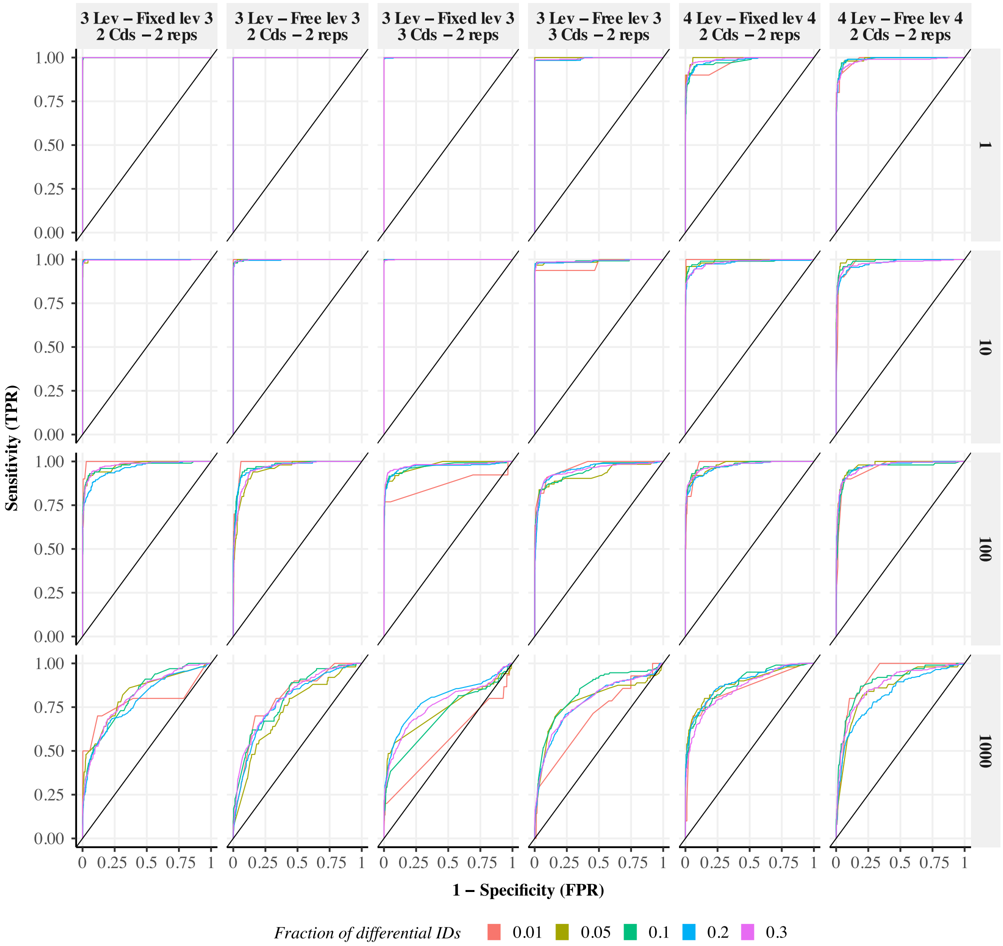


**Fig S7. ROC curves describing GPMelt performance on the** 120 **tested synthetic dataset configurations.** The columns correspond to the different scenarios (i.e HGP model specifications) and the rows to different noise factors. The colors of the curves indicate the fraction of true alternative IDs among the $N = 1000$ simulated IDs. The black lines indicate the curves $y = x$, representing the performance of a random classifier.

#### S17 Fig.

**Comparison of p-values histograms for the benchmarking datasets (NPARC vs GPMelt).**

#### S18 Fig.

**Schematic visualisation of the phospho-TPP dataset [11] analysis.**

#### S19 Fig.

**Additional results on the phospho-TPP dataset [11] analysis.**

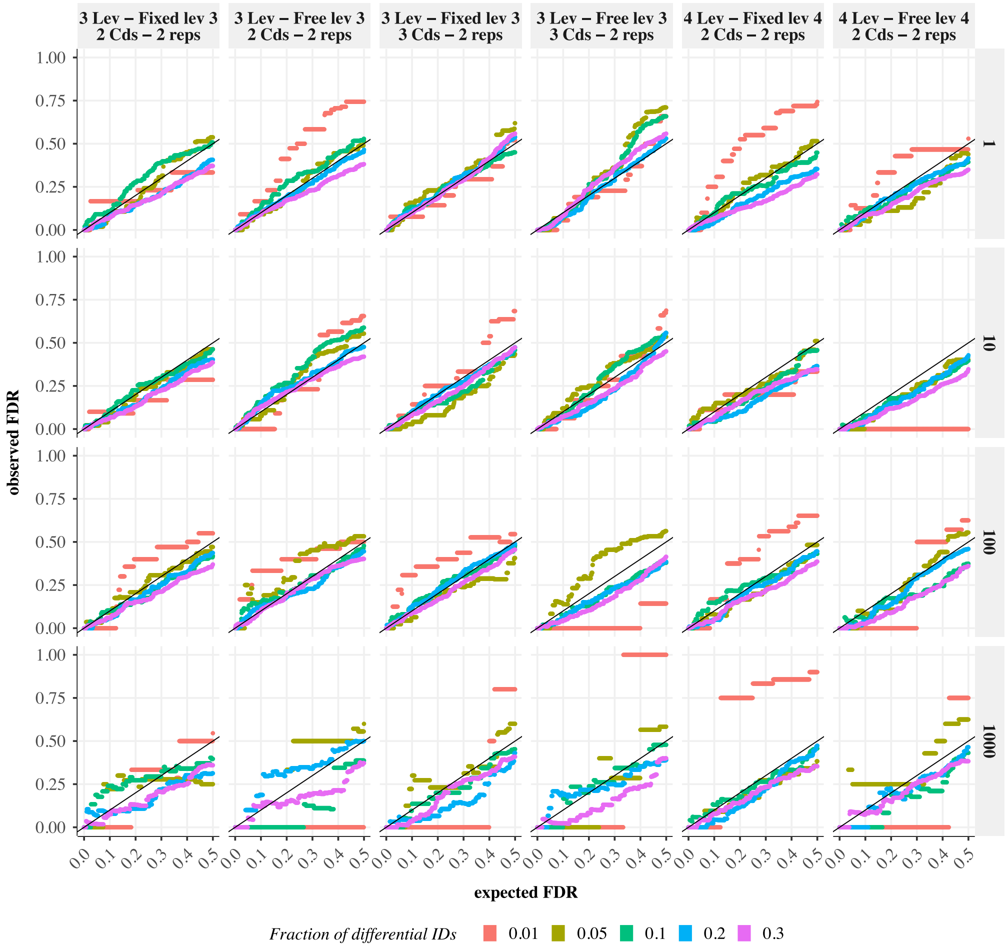


**Fig S8. Expected vs observed FDR for the 120 tested synthetic datasets configurations.** The columns correspond to the different scenarios (i.e HGP model specifications) and the rows to different noise factors. The colors of the curves indicate the fraction of true differential IDs among the $N = 1000$ simulated IDs. The black lines indicate the curves $y = x$, representing the ideal case.

#### S20 Fig.

**ABC vs absolute ABC.**

#### S21 Fig.

**The mean scaling is more appropriate than the median or geometric mean scaling when considering melting curves.**

#### S22 Fig.

**The mean is more stable than the median or geometric mean when considering melting curve observations.**

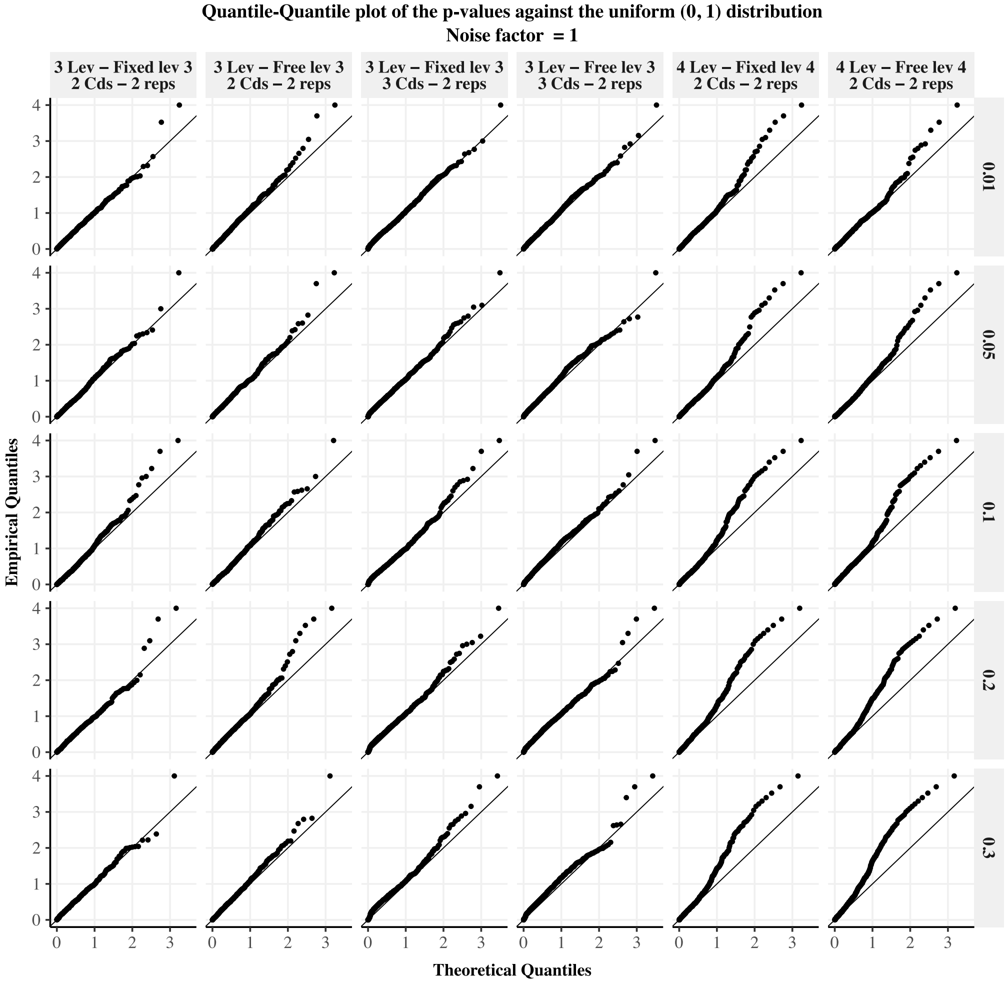


**Fig S9. QQPlots obtained for a noise factor of 1, considering the different synthetic datasets scenarios and proportions of differential IDs.** The columns correspond to the different scenarios (i.e HGP model specifications) and the rows to different fractions of true differential IDs among the $N = 1000$ simulated IDs. The black lines indicate the curves $y = x$, on which uniformly $[0, 1]$ distributed p-values should lie.

### S23 Fig.

**Computational cost of the HGP model fitting and null distribution approximation for the ATP2019 and phosphoTPP datasets.**

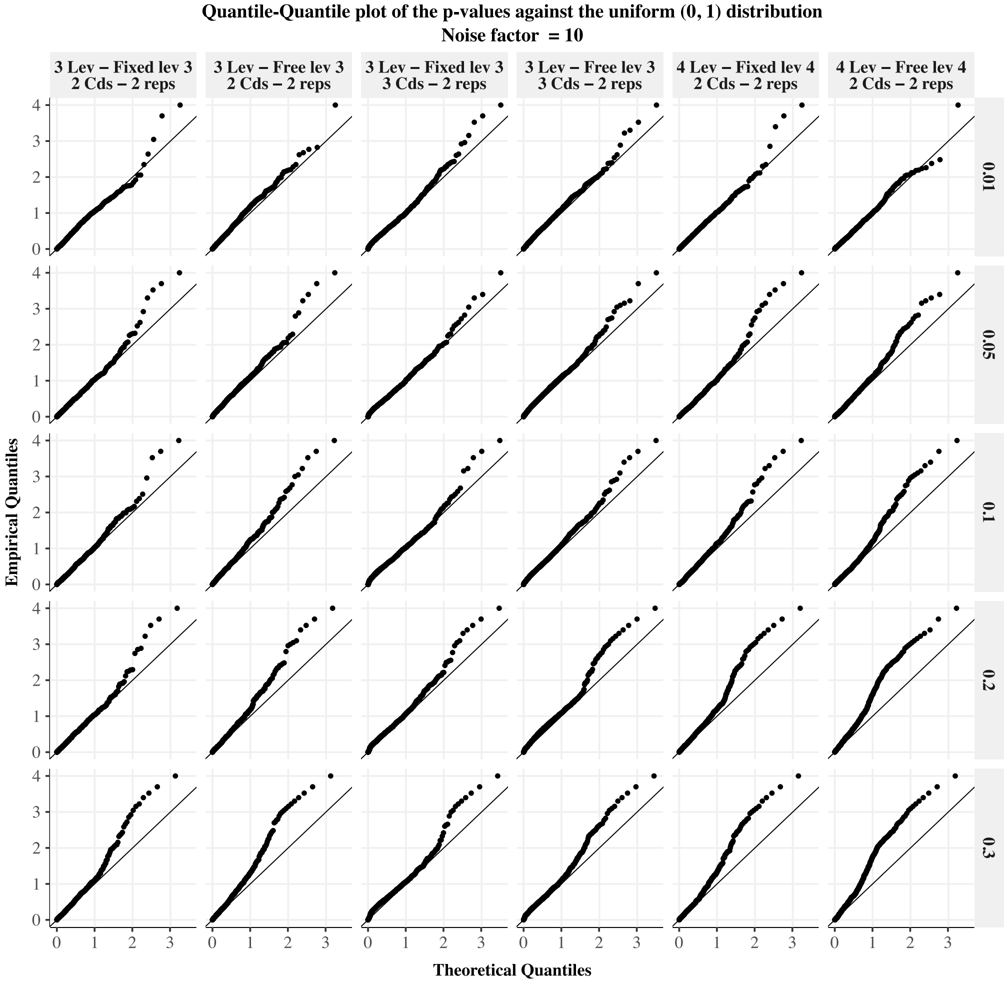


**Fig S10. QQPlots obtained for a noise factor of 10, considering the different synthetic datasets scenarios and proportions of differential IDs.** The columns correspond to the different scenarios (i.e HGP model specifications) and the rows to different fractions of true differential IDs among the $N = 1000$ simulated IDs. The black lines indicate the curves $y = x$, on which uniformly $[0, 1]$ distributed p-values should lie.

#### S1 Table: Models comparison

#### S2 Table: Description of the benchmarking datasets.

#### S3 Table: Model specifications used for the benchmarking datasets.

#### S4 Table: Null distribution approximation methods.

#### References

1. Savitski MM, Reinhard FB, Franken H, Werner T, Savitski MF, Eberhard D, et al. Tracking cancer drugs in living cells by thermal profiling of the proteome.

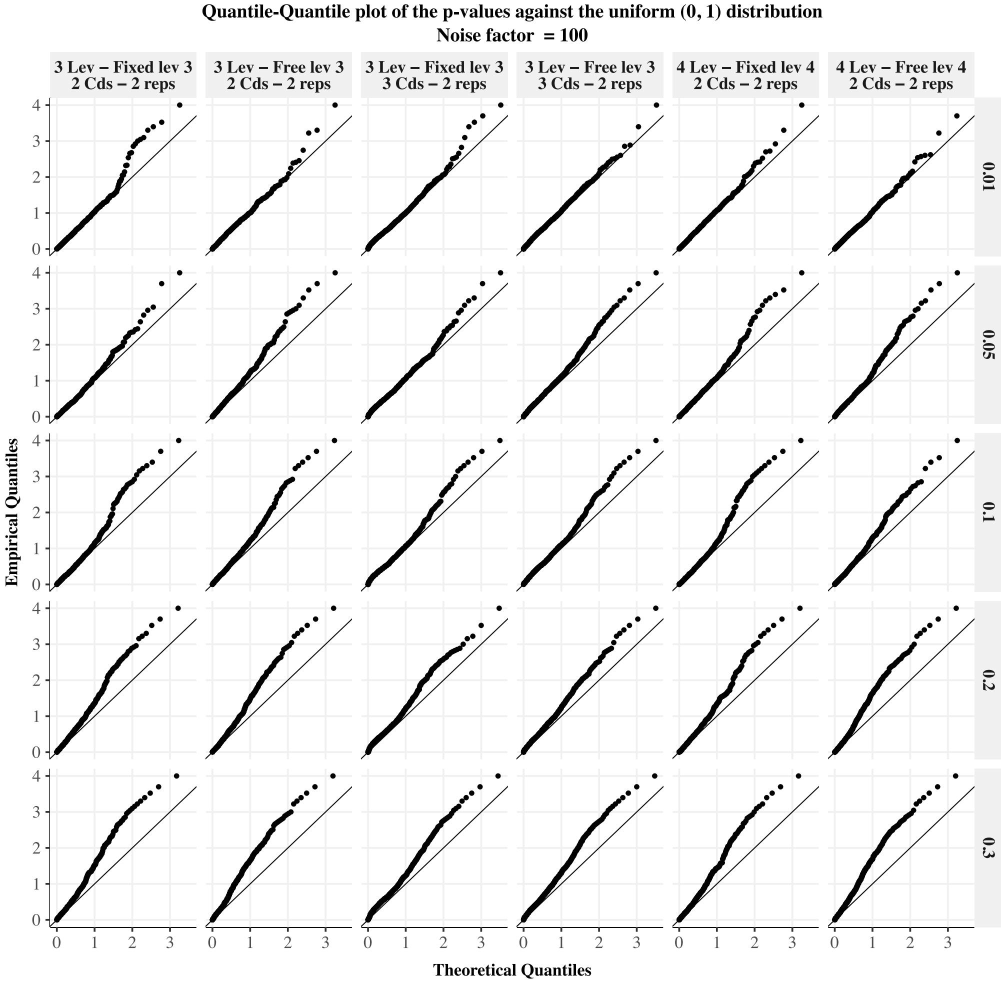


**Fig S11. QQPlots obtained for a noise factor of** 100**, considering the different synthetic datasets scenarios and proportions of differential IDs.** The columns correspond to the different scenarios (i.e HGP model specifications) and the rows to different fractions of true differential IDs among the $N = 1000$ simulated IDs. The black lines indicate the curves $y = x$, on which uniformly $[0, 1]$ distributed p-values should lie.

Science. 2014;346(6205):1255784.

2. Molina DM, Jafari R, Ignatushchenko M, Seki T, Larsson EA, Dan C, et al. Monitoring drug target engagement in cells and tissues using the cellular thermal shift assay. Science. 2013;341(6141):84–87.

3. Bantscheff M, Lemeer S, Savitski MM, Kuster B. Quantitative mass spectrometry in proteomics: critical review update from 2007 to the present. Analytical and bioanalytical chemistry. 2012;404:939–965.

4. Werner T, Sweetman G, Savitski MF, Mathieson T, Bantscheff M, Savitski MM. Ion coalescence of neutron encoded TMT 10-plex reporter ions. Analytical chemistry. 2014;86(7):3594–3601.

5. Becher I, Werner T, Doce C, Zaal EA, Tögel I, Khan CA, et al. Thermal profiling reveals phenylalanine hydroxylase as an off-target of panobinostat. Nature chemical biology. 2016;12(11):908–910.

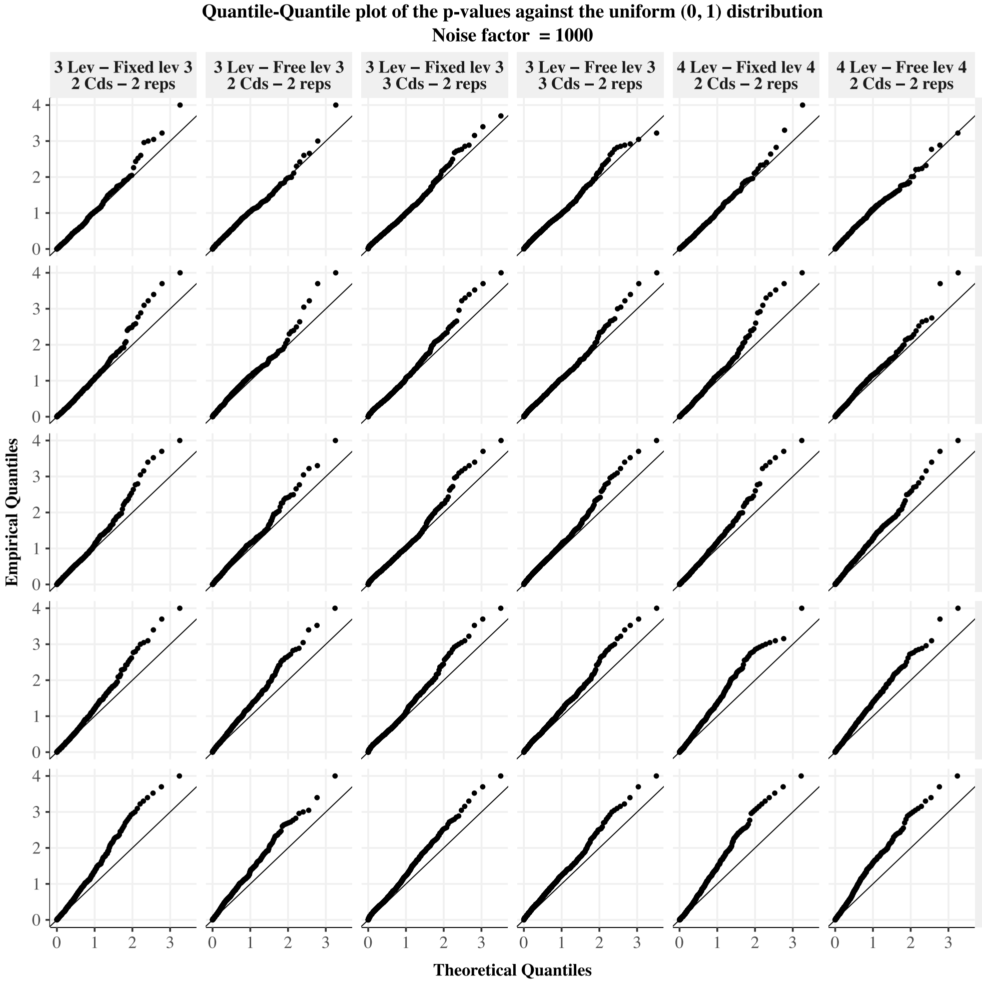


**Fig S12. QQPlots obtained for a noise factor of 1000, considering the different synthetic datasets scenarios and proportions of differential IDs.** The columns correspond to the different scenarios (i.e HGP model specifications) and the rows to different fractions of true differential IDs among the $N = 1000$ simulated IDs. The black lines indicate the curves $y = x$, on which uniformly $[0, 1]$ distributed p-values should lie.

6. Childs D, Bach K, Franken H, Anders S, Kurzawa N, Bantscheff M, et al. Nonparametric analysis of thermal proteome profiles reveals novel drug-binding proteins. Molecular & Cellular Proteomics. 2019;18(12):2506–2515.

7. Kurzawa N, Becher I, Sridharan S, Franken H, Mateus A, Anders S, et al. A computational method for detection of ligand-binding proteins from dose range thermal proteome profiles. Nature communications. 2020;11(1):5783.

8. Franken H, Mathieson T, Childs D, Sweetman GM, Werner T, Tögel I, et al. Thermal proteome profiling for unbiased identification of direct and indirect drug targets using multiplexed quantitative mass spectrometry. Nature protocols. 2015;10(10):1567–1593.

9. Becher I, Andrés-Pons A, Romanov N, Stein F, Schramm M, Baudin F, et al. Pervasive protein thermal stability variation during the cell cycle. Cell.

**Table S1. Models comparison** (*) with $X$ the number of levels in the HGP model. Constraints on the lengthscales can be added to reduce the number of parameters. (**) $c$ the condition, $r$ the replicate. Constraints on the output-scales can be added to reduce the number of parameters. (***) For the four-level HGP model: $c$ the condition, $j$ the peptide, $r$ the replicate. Constraints on the output-scales can be added to reduce the number of parameters.

| Model | NPARC | Bayesian sigmoid | Bayesian semi-parametric | GPMelt |
|---|---|---|---|---|
| Reference | Childs et al. [6] | Fang et al. [16] | | this paper |
| Mean $m(\cdot)$ | $S_{a,b,p}(t) = \frac{1-p}{1+\exp(b-\frac{a}{t})} + p$ | | | 0 |
| Underlying assumption concerning the mean | Sigmoidal mean | | | No a-priori assumption |
| Kernel $k(\cdot,\cdot)$ | RBF kernel with null output-scale | | RBF kernel | Combination of RBF kernels |
| Error term $\epsilon_i$ | $\epsilon_i \overset{iid}{\sim} \mathcal{N}(0, \beta^2)$ | | | |
| Parameters and hyperparameters | $a, b, p, \beta$ | $a, b, p, \beta$ | $a, b, p, \beta$ $\lambda, \sigma$ | $\beta, \sigma_h, \sigma_g$<br>$\{\lambda_i\}_{i=1}^{X}$ (*)<br>$\{\sigma_{f_{cr}}\}_{c,r}$ (**)<br>$\{\sigma_{\eta_{c\pi_j r}}\}_{c,j,r}$ (***) |
| Priors on | None | $a, b, p, \beta$<br><br>null and alternative models probabilities | $a, b, p, \beta$<br>$\lambda, \sigma$<br>null and alternative models probabilities | None |
| Estimation procedure | Non-linear least square estimation | Hamiltonian Monte-Carlo sampling | | Type II Maximum likelihood Estimation |
| Programming language | R | RStan | | Python and R (combined in a Nextflow pipeline) |

2018;173(6):1495–1507.

10. Mateus A, Hevler J, Bobonis J, Kurzawa N, Shah M, Mitosch K, et al. The functional proteome landscape of Escherichia coli. Nature. 2020;588(7838):473–478.

11. Potel CM, Kurzawa N, Becher I, Typas A, Mateus A, Savitski MM. Impact of phosphorylation on thermal stability of proteins. Nature methods. 2021;18(7):757–759.

12. Mateus A, Kurzawa N, Becher I, Sridharan S, Helm D, Stein F, et al. Thermal proteome profiling for interrogating protein interactions. Molecular systems biology. 2020;16(3):e9232.

13. Le Sueur C, Hammarén HM, Sridharan S, Savitski MM. Thermal proteome profiling: Insights into protein modifications, associations, and functions. Current Opinion in Chemical Biology. 2022;71:102225.

**Table S2. Description of the benchmarking datasets.**

| | Dataset | Comparison | Approximation of ground truth | Number of IDs | Other applied methods and references to results (if applicable) |
|---|---|---|---|---|---|
| protein-level TPP-TR two conditions | ATP 2015 [32] | Vehicle vs Mg-ATP [2$\mu$M] | GO term: ATP-binding proteins | 4177 proteins | $T_m$ and NPARC [6] Bayesian sigmoid [16] Bayesian semi-parametric models [16] |
| | Panobinostat [8] | Vehicle vs Panobinostat [1$\mu$M] | | 3649 proteins | |
| | Staurosporine 2014 [1] | Vehicle vs Staurosporine [20$\mu$M] | GO term: protein kinase activity | 4505 proteins | |
| | Staurosporine 2021 [31] | Vehicle vs Staurosporine [20$\mu$M] | GO term: protein kinase activity | 4403 proteins | $T_m$ [31]; NPARC |
| | ATP 2019 [18] | Vehicle vs Mg-ATP [10 mM] | GO term: ATP-binding proteins | 4772 proteins | NPARC |
| protein-level TPP-TR multiple conditions | Dasatinib [1] | Condition C1: Vehicle vs Dasatinib [0.5$\mu$M] Condition C2: Vehicle vs Dasatinib [5$\mu$M] | | 4768 proteins | $T_m$ and NPARC [6] Bayesian sigmoid [16] Bayesian semi-parametric models [16] |
| peptide-level TPP-TR multiple conditions | phospho-TPP [11] | Phospho-peptides vs median of non-phosphorylated peptides | Functional scores [33] | 13990 phospho-peptides from 1949 proteins | $T_m$ [11] |
| | phospho-TPP [11] | Comparison of phosphorylation patterns | | 4073 phospho-peptides from 310 proteins | |


14. Kurzawa N, Stahl M, Leo I, Kunold E, Becher I, Audrey A, et al. Deep thermal proteome profiling for detection of proteoforms and drug sensitivity biomarkers. bioRxiv. 2022; p. 2022–06.

15. Schellman JA. The thermodynamics of solvent exchange. Biopolymers: Original Research on Biomolecules. 1994;34(8):1015–1026.

16. Fang S, Kirk PD, Bantscheff M, Lilley KS, Crook OM. A Bayesian semi-parametric model for thermal proteome profiling. Communications biology. 2021;4(1):1–15.

17. Ruan C, Ning W, Liu Z, Zhang X, Fang Z, Li Y, et al. Precipitate-Supported Thermal Proteome Profiling Coupled with Deep Learning for Comprehensive Screening of Drug Target Proteins. ACS Chemical Biology. 2022;17(1):252–262.

18. Sridharan S, Kurzawa N, Werner T, Günthner I, Helm D, Huber W, et al. Proteome-wide solubility and thermal stability profiling reveals distinct regulatory roles for ATP. Nature communications. 2019;10(1):1–13.

**Table S3. Model specifications used for the benchmarking datasets.** (1) *The minimum distance between successive temperatures is used as a lower limit for the lengthscale.*(2) *For very complex models, multiple optima could exist. Hence, the fitting of the observations for each protein has been estimated three independent times, and the parameters leading to the best fit (lowest final loss value from the type II MLE estimation) are selected for the null distribution sampling and the rest of the analysis.* (3) *The maximum distance between successive temperatures is used as a lower limit for the lengthscale, to favour smoother fit estimations.*

| | Dataset | HGP model | Scaling | Constraints on the length-scales | Constraints on the output-scales | Null distribution approximation (see S4 Table) |
|---|---|---|---|---|---|---|
| protein-level TPP-TR two conditions | ATP 2015 [32] | Three-level HGP model | Fold change | $\lambda_1 = \lambda_2 = \lambda_3 \equiv \lambda$<br>$\lambda > min_i\lvert T_{i+1} - T_i\rvert(1)$ | None | Method A with $S_p = 10$ |
| | Panobinostat [8] | | | | | |
| | Staurosporine 2014 [1] | | | | | |
| | Staurosporine 2021 [31] | | | | | |
| | ATP 2019 [18] | | | | $\sigma_{f_{cr}} \equiv \sigma_f \quad \forall c, r$ | |
| protein-level TPP-TR multiple conditions | Dasatinib [1] | Three-level HGP model | Fold change | $\lambda_1 = \lambda_2 = \lambda_3 \equiv \lambda$<br>$\lambda > min_i\lvert T_{i+1} - T_i\rvert(1)$ | None | Method A with $S_p = 10$ |
| peptide-level TPP-TR multiple conditions | phospho-TPP [11]; *Phospho-peptides vs median of non-phosphorylated peptides* | Three-level HGP model (2) | Mean scaling | $\lambda_1 = \lambda_2 \equiv \lambda \neq \lambda_3$<br>$\lambda > max_i\lvert T_{i+1} - T_i\rvert(3)$ | None | Protein-wise: method D with $S_p \equiv S = 1e4$<br>Group-wise: method E with $S = 3e5$ |
| | phospho-TPP [11]; *Comparison of phosphorylation patterns* | Four-level HGP model | Mean scaling | $\lambda_1 = \lambda_2 = \lambda_3 \equiv \lambda \neq \lambda_4$<br>$\lambda > max_i\lvert T_{i+1} - T_i\rvert(3)$ | $\sigma_{f_{c\pi_j}} \equiv \sigma_f \;\forall j, \forall c$ | Protein-wise: method D with $S_p \equiv S = 1e4$ |


19. Brangwynne CP, Eckmann CR, Courson DS, Rybarska A, Hoege C, Gharakhani J, et al. Germline P granules are liquid droplets that localize by controlled dissolution/condensation. Science. 2009;324(5935):1729–1732.

20. Hyman AA, Weber CA, Jülicher F. Liquid-liquid phase separation in biology. Annual review of cell and developmental biology. 2014;30:39–58.

21. Dignon GL, Zheng W, Kim YC, Mittal J. Temperature-controlled liquid–liquid phase separation of disordered proteins. ACS central science. 2019;5(5):821–830.

22. Williams CK, Rasmussen CE. Gaussian processes for machine learning. vol. 2. MIT press Cambridge, MA; 2006.

23. Bates DM, Watts DG. Nonlinear regression analysis and its applications. vol. 2. Wiley New York; 1988.

24. Hensman J, Lawrence ND, Rattray M. Hierarchical Bayesian modelling of gene expression time series across irregularly sampled replicates and clusters. BMC bioinformatics. 2013;14(1):1–12.

25. Bonilla EV, Chai K, Williams C. Multi-task Gaussian process prediction. Advances in neural information processing systems. 2007;20.

26. Kass RE, Raftery AE. Bayes factors. Journal of the american statistical association. 1995;90(430):773–795.

27. Liu Z, Barahona M. Similarity measure for sparse time course data based on Gaussian processes. In: Uncertainty in Artificial Intelligence. PMLR; 2021. p. 1332–1341.

28. Phillips NE, Manning C, Papalopulu N, Rattray M. Identifying stochastic oscillations in single-cell live imaging time series using Gaussian processes. PLoS computational biology. 2017;13(5):e1005479.

29. Aebersold R, Agar JN, Amster IJ, Baker MS, Bertozzi CR, Boja ES, et al. How many human proteoforms are there? Nature chemical biology. 2018;14(3):206–214.

30. Smith LM, Kelleher NL. Proteoform: a single term describing protein complexity. Nature methods. 2013;10(3):186–187.

31. Zinn N, Werner T, Doce C, Mathieson T, Boecker C, Sweetman G, et al. Improved proteomics-based drug mechanism-of-action studies using 16-Plex isobaric mass tags. Journal of Proteome Research. 2021;20(3):1792–1801.

32. Reinhard FB, Eberhard D, Werner T, Franken H, Childs D, Doce C, et al. Thermal proteome profiling monitors ligand interactions with cellular membrane proteins. Nature methods. 2015;12(12):1129–1131.

33. Ochoa D, Jarnuczak AF, Viéitez C, Gehre M, Soucheray M, Mateus A, et al. The functional landscape of the human phosphoproteome. Nature biotechnology. 2020;38(3):365–373.

34. UniProt: the Universal Protein knowledgebase in 2023. Nucleic Acids Research. 2023;51(D1):D523–D531.

35. Benjamini Y, Hochberg Y. Controlling the false discovery rate: a practical and powerful approach to multiple testing. Journal of the Royal statistical society: series B (Methodological). 1995;57(1):289–300.

36. Beltrao P, Bork P, Krogan NJ, van Noort V. Evolution and functional cross-talk of protein post-translational modifications. Molecular Systems Biology. 2013;9(1):714. doi:https://doi.org/10.1002/msb.201304521.

37. Vantini M, Mannerström H, Rautio S, Ahlfors H, Stockinger B, Lähdesmäki H. PairGP: Gaussian process modeling of longitudinal data from paired multi-condition studies. Computers in Biology and Medicine. 2022;143:105268. doi:https://doi.org/10.1016/j.compbiomed.2022.105268.

38. Jung F, Frey K, Zimmer D, Mühlhaus T. DeepSTABp: A Deep Learning Approach for the Prediction of Thermal Protein Stability. International Journal of Molecular Sciences. 2023;24(8):7444.

39. Gardner, Jacob R and Pleiss, Geoff and Bindel, David and Weinberger, Kilian Q and Wilson, Andrew Gordon GPyTorch: Blackbox Matrix-Matrix Gaussian Process Inference with GPU Acceleration Advances in Neural Information Processing Systems, 2018.

40. Wikipedia contributors Hadamardd product (matrices) — Wikipedia, The Free Encyclopedia `https://en.wikipedia.org/w/index.php?title=Hadamardd_product_(matrices)&oldid=1122748935` accessed 5-December-2022

41. Wikipedia contributors Kronecker product — Wikipedia, The Free Encyclopedia `https://en.wikipedia.org/w/index.php?title=Kronecker_product&oldid=1097138322` accessed 12-August-2022

42. Phipson, Belinda and Smyth, Gordon K Permutation P-values should never be zero: calculating exact P-values when permutations are randomly drawn Statistical applications in genetics and molecular biology. 2010

43. Kingma, Diederik P and Ba, Jimmy Adam: A method for stochastic optimization arXiv preprint arXiv:1412.6980. 2014

44. Wu T, Hu E, Xu S, Chen M, Guo P, Dai Z, et al. clusterProfiler 4.0: A universal enrichment tool for interpreting omics data. The innovation. 2021;2(3).

45. Di Tommaso P, Chatzou M, Floden EW, Barja PP, Palumbo E, Notredame C. Nextflow enables reproducible computational workflows. Nature biotechnology. 2017;35(4):316–319.

46. R Core Team. R: A Language and Environment for Statistical Computing; 2023. Available from: `https://www.R-project.org/`.

47. Kassambara A. ggpubr: 'ggplot2' Based Publication Ready Plots; 2023. Available from: `https://CRAN.R-project.org/package=ggpubr`.

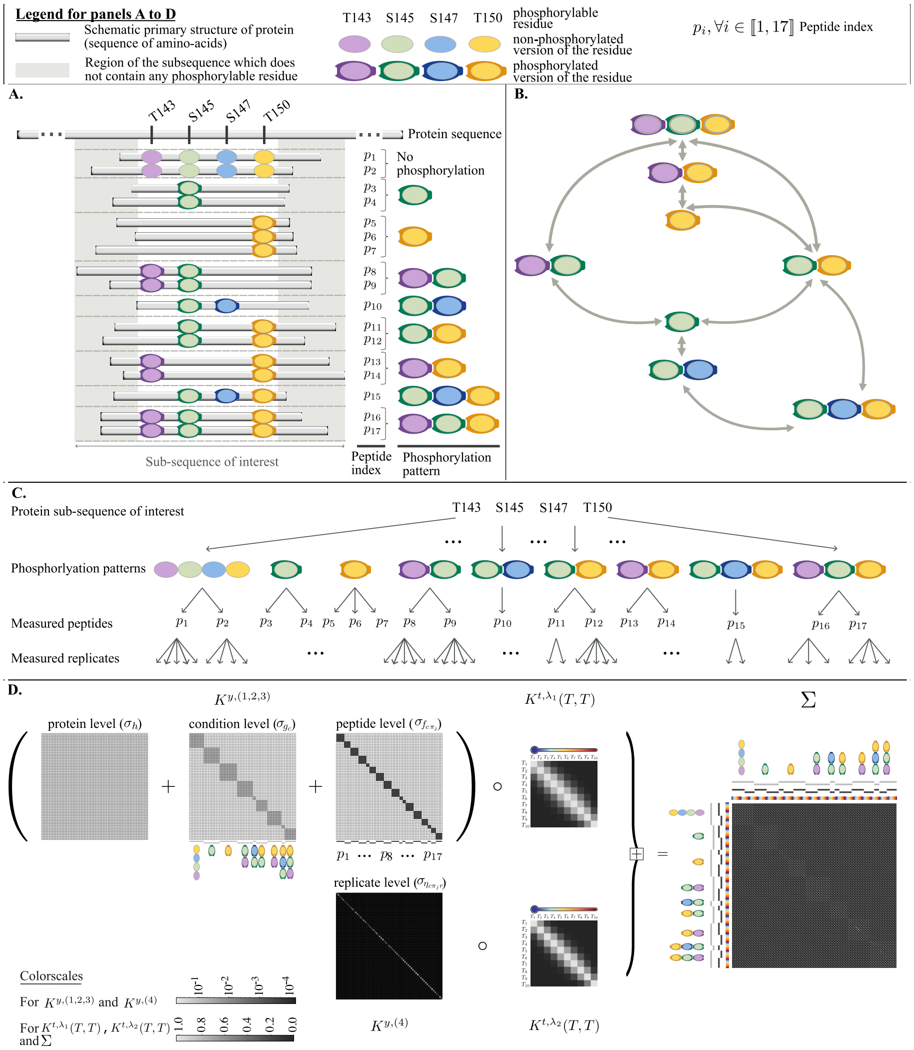


**Fig S13. Extension to deeper hierarchies (schematic description)** To complement the approach presented in Fig 5 (main text) aiming to detect functionally relevant phosphosites, we propose an analysis focused on the phosphorylation patterns instead of the phospho-peptides, leading to more interpretable biological results. We schematically illustrate here how to apply this method to the phosphoTPP peptide-level TPP-TR dataset [11]. Especially, we focus on a real example considering a subsequence of the MARCKS protein (amino acids 136 to 158). (A) A schematic representation of the available data for this sub-sequence. Exactly four phosphorylable residues, namely *T143, S145, S147, T150*, are located on this sub-sequence. 17 peptides spanning this region are observed, presenting nine different phosphorylation patterns, including no phosphorylation at all. (B) A visualisation of the links between these phosphorylation patterns. Depending on the user and his/her biological focus, the comparison of some melting behaviours might be of special interest. However, no definition of the *control* condition is required before model fitting. The statistic $\Lambda$ associated to any comparison is directly available from the fit of the four-level hierarchical model presented in panel (C). In this model, corresponding to Eq (52), the additional level in the hierarchy groups peptides by phosphorylation patterns (considered as *conditions*). (D) Illustration of the covariance matrix $\Sigma$ corresponding to a constrained version of Eq (52). To reduce the model complexity (and as described in more details in S1 Supporting Information), the following constraints have been introduced: $\lambda_1 = \lambda_2$ *and* $\sigma_{f_{c\pi_j}} \equiv \sigma_f \;\; \forall j, \forall c$. As derived in S2 Appendix, the full covariance matrix $\Sigma$ is given by $K^{y,(1,2,3)} \circ K^{t,\lambda_1}(T,T) + K^{y,(4)} \circ K^{t,\lambda_2}(T,T)$. $K^{y,(1,2,3)}$ is obtained by summing the index kernels corresponding to protein, condition and peptide levels (levels one to three).

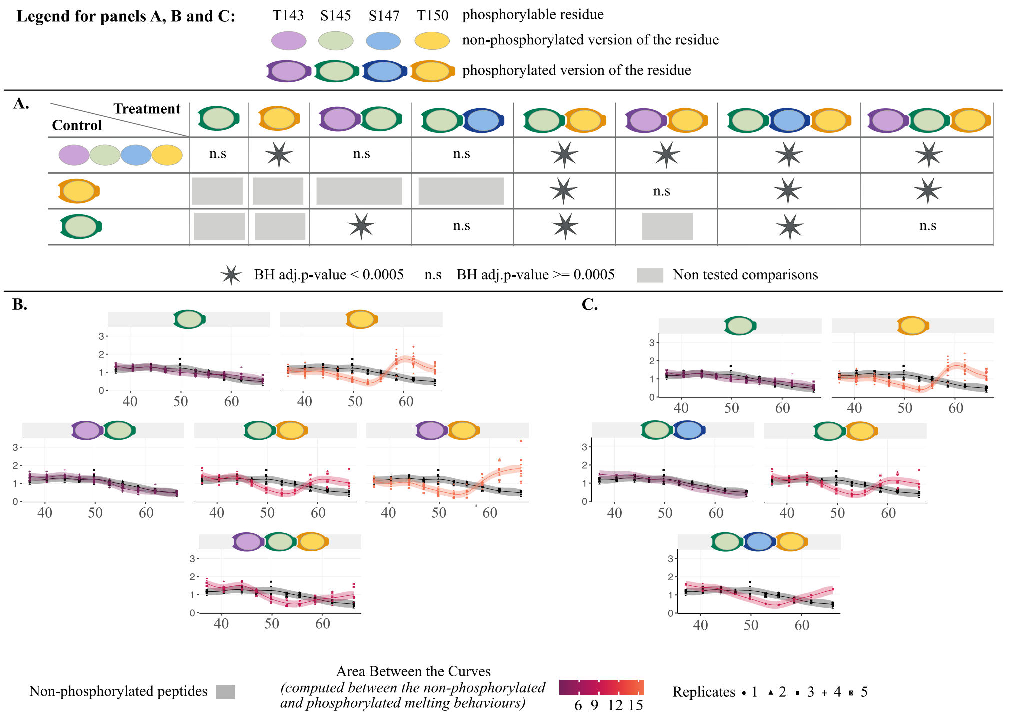


**Fig S14. Extension to deeper hierarchies (real example).** Considering the real example schematically presented in S13 Fig, we explore the principle of cross-talk between phosphorylation events, i.e. whether evidences of PTMs cooperativity can be observed [36]. We are considering a sub-sequence of MARCKS protein containing exactly four phosphorylable residues, namely *T143, S145, S147, T150*. Nine different phosphorylation patterns, including no phosphorylation at all, and a total of 17 peptides, are observed. As mentioned previously, multiple *control* conditions could be chosen to explore different biological questions. In panel (A) we propose three sets of comparisons, one per row. In the first row, we consider investigating the effect on the melting behaviour of any phosphorylation events, and thus choose the non phosphorylated peptides as *control* condition. This comparison strikingly shows that only phosphorylation patterns containing $pT150$ are found to induce a significant change in melting behaviour. Following this observation, we wondered if $pT150$ alone is responsible for this change in melting behaviour, and thus if the melting behaviours of all phosphorylation patterns including $pT150$ are similar. In the second row, using *pS150* as control condition, we show that {*pS145, pT150*}, {*pT143, pS145, pT150*} and {*pS145, pS147, pT150*} present a melting behaviours significantly different from the melting behaviour of $pT150$ alone. This suggests a cross-talk between these phosphorylated sites. Indeed, the addition of other phosphorylation events next to $pT150$ seem to have a different effect on the protein than the single phosphorylation of *T150*. Similarly, one could be interested in the possible cross-talk between *T143, S145* and *S147* phosphosites. Using *pS145* as control condition in the last row, we show that {*pT143, pS145*} but not {*pS145, pS147*} present a significantly different melting behaviour than *pS145*. An independent null distributions of size $S = 1e4$ for each comparison has been derived, and results are presented with an $\alpha$-level of 0.0005 on the BH adjusted p-values. Panels (B) and (C) propose a visualisation of the different melting behaviours, with the black curves corresponding to the non-phosphorylated peptides, and the colored curves to the phosphorylated peptides. The color reflects the absolute Area Between the Curve ($|ABC|_{GPMelt}$) between the non-phosphorylated peptides and the phosphorylated peptides. The $|ABC|_{GPMelt}$ has been computed using the predictive distribution of the four-level HGP model, as described in S4 Appendix, Eq (51).

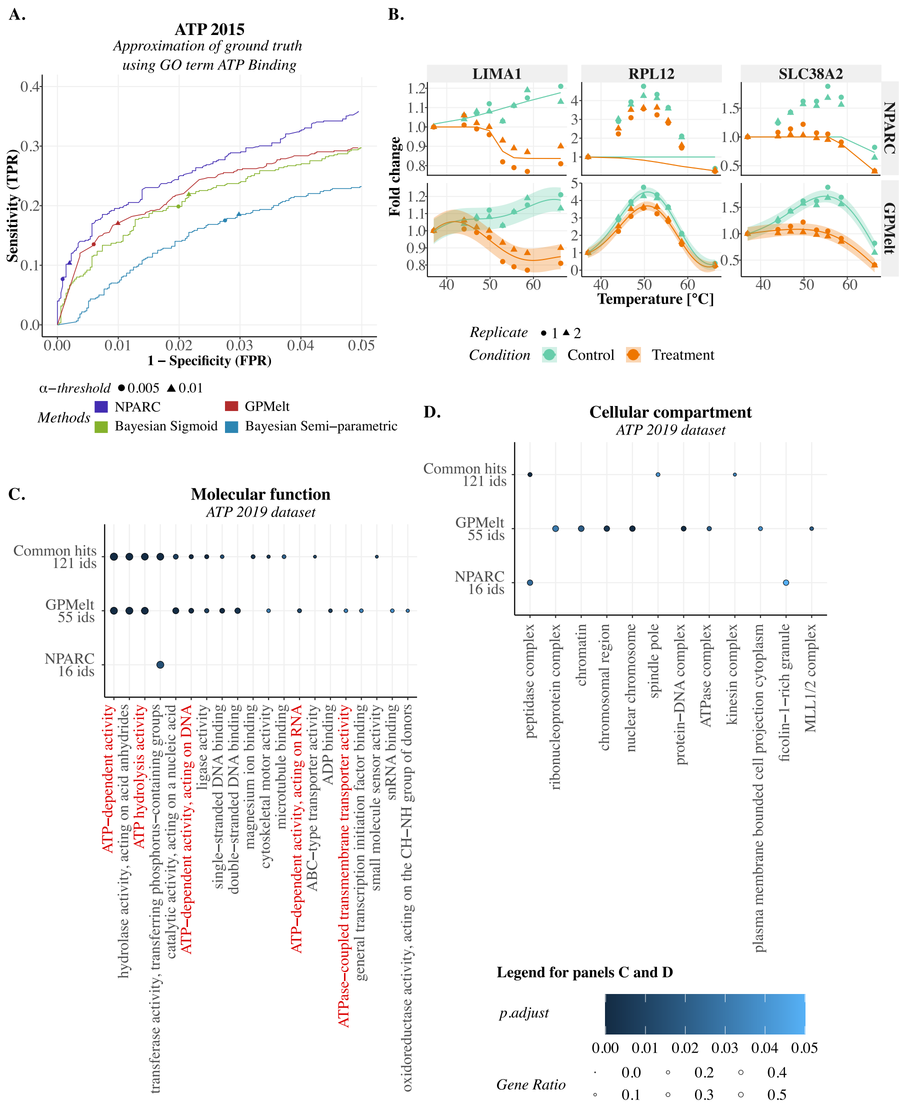


**Fig S15. Additional results for protein-level TPP-TR datasets.** (A) [ATP 2015 dataset [32]] Approximate receiver operator characteristic (ROC) curves comparing the results of NPARC [6], the Bayesian sigmoid model [16], the Bayesian semi-parametric model [16] and GPMelt on the ATP 2015 [32] dataset. The set of proteins expected to be targeted by the treatment are defined using the Gene Ontology (GO) Consortium annotations curated in Uniprot [34] (annotations downloaded in march 2023). 599 out of 4177 proteins are annotated as ATP binding proteins. The points on the curves correspond to the sensitivity and specificity of NPARC and GPMelt at an $\alpha$-threshold of $\alpha \in \{0.005, 0.01\}$ on the BH adjusted p-values, resp. a threshold of $1-\alpha$ on the posterior probabilities of the alternative model for the Bayesian sigmoid and Bayesian semi-parametric models. (B) [Staurosporine 2021 dataset [31]] Example of three proteins found to have a low rank according to NPARC analysis, while being selected as top proteins by GPMelt. NPARC ranking is due to NPARC fits either failing (LIMA, joint model fit failed) or miss-fitting the actual melting curves (SLC38A2 and RPL12). (C-D) [ATP 2019 dataset [18]] Dotplot of gene ontology molecular function (panel C) and cellular compartment (panel D) and enrichment results. The enrichment analysis is performed independently for the 16 unique hits of NPARC (captured with an $\alpha$-threshold of 0.05 on the BH adjusted p-values), the 55 unique hits of GPMelt (captured with an $\alpha$-threshold of 0.001 on the BH adjusted p-values) and the 121 common hits. The enrichment analysis is performed with the R package clusterProfiler [44] (v4.8.3), with background defined by the set of proteins identified in the experiment.

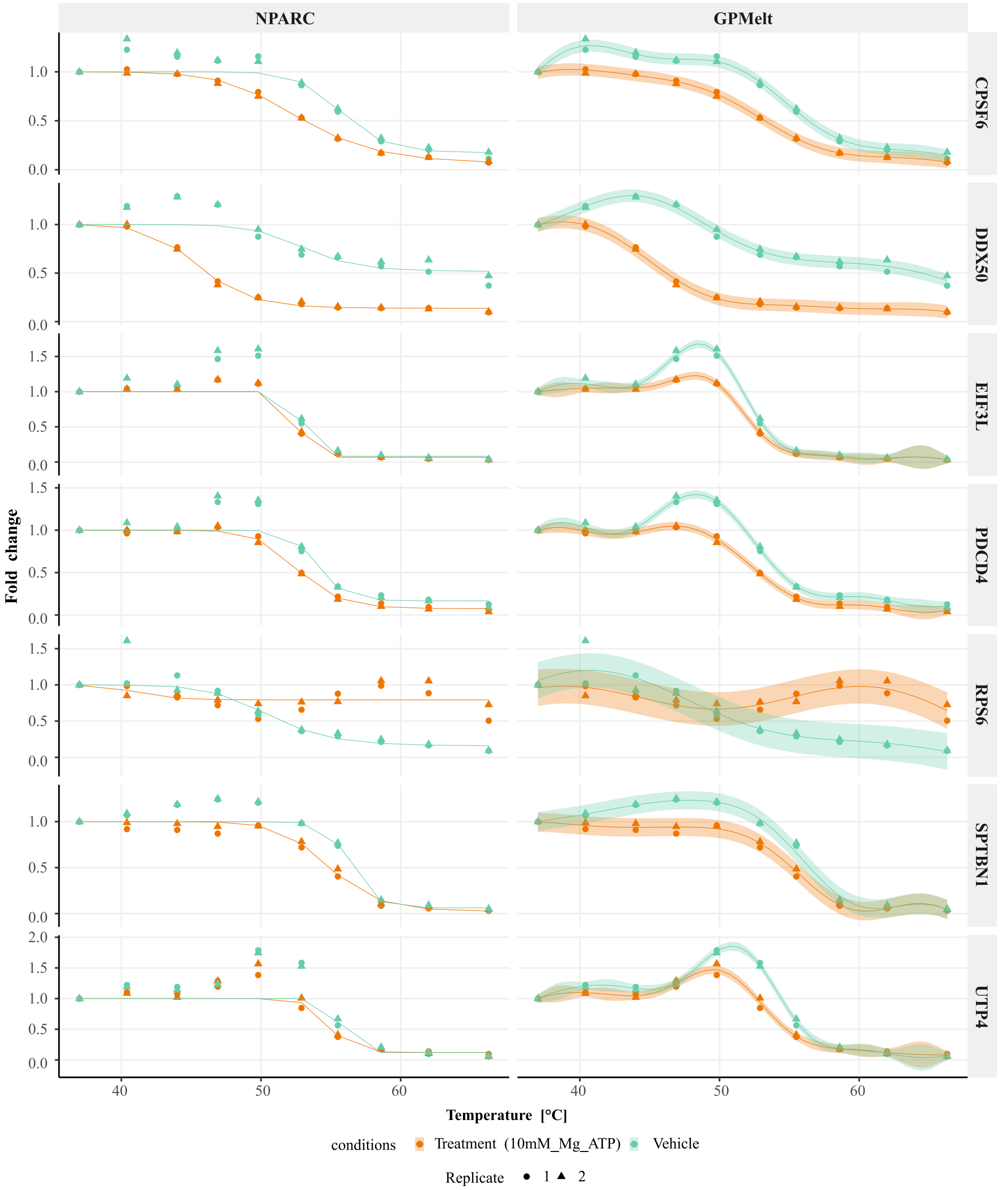


**Fig S16. Additional examples of non sigmoidal curves (ATP 2019 dataset [18])** Examples of the fits obtained for NPARC (left column) vs a three-level HGP model (right column). These proteins are depicted by crosses in Fig 4C of the main text.

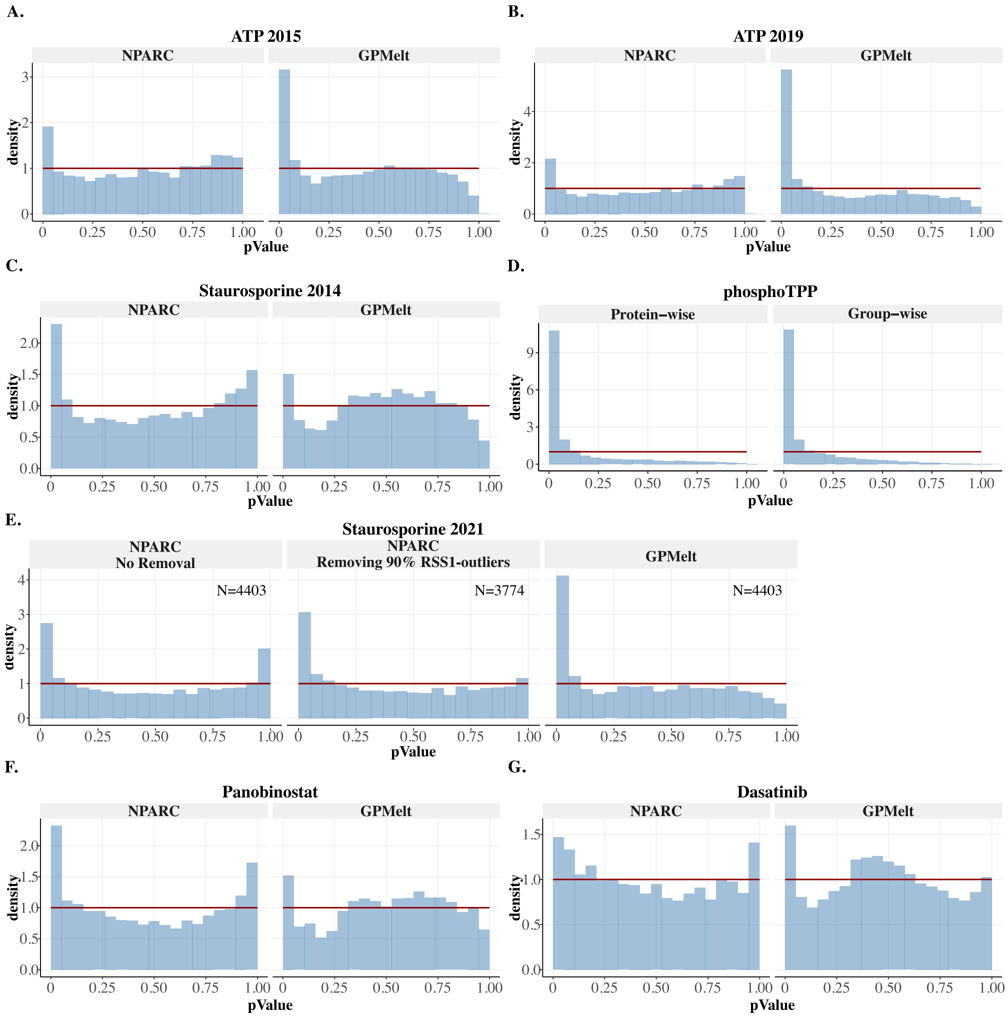


**Fig S17. Comparison of p-values histograms for the benchmarking datasets.** Blue: p-values histograms obtained for each benchmarking dataset, in function of the method used. Red horizontal line: uniform distribution betwen $[0, 1]$, expected if all proteins would follow the null hypothesis. It can be noticed that the p-value histograms of NPARC for datasets Saturosporine 2014 (C), Panobinostat (F) and Dasatinib (G) suffer from the presence of a peak on the right, suggesting that the assumption of uniform distribution of p-values under the null is not valid. Similarly, the application of NPARC to the Staurosporine 2021 dataset [31] (E) show a clear peak on the right (left-most panel). We suggest that most proteins with associated extreme p-values would present non-sigmoidal melting curves for at least one of the two conditions (control and/or treatment). These curves are fitted with NPARC's model for the alternative hypothesis $H_1$. Non-sigmoidal melting curves lead to poor sigmoidal fits, i.e presenting large Residual Sum of Square (denoted by $RSS_1$ from NPARC notation). After computing the $90th$-percentile of the $RSS_1$ values for over all proteins, we removed the proteins having an $RSS_1$ value above the $90th$-percentile. Plotting the p-values histogram after this removal (middle panel) allows to recover an expected shape. (Right panel) The p-values histogram obtained after applying GPMelt on the dataset without any filtering of the proteins. (D) P-values histograms for GPMelt with a three-level HGP model applied on the phospho-TPP dataset (case phospho-peptides vs median of non-phosphorylated peptides) for either the protein-wise or group-wise null distribution approximation.

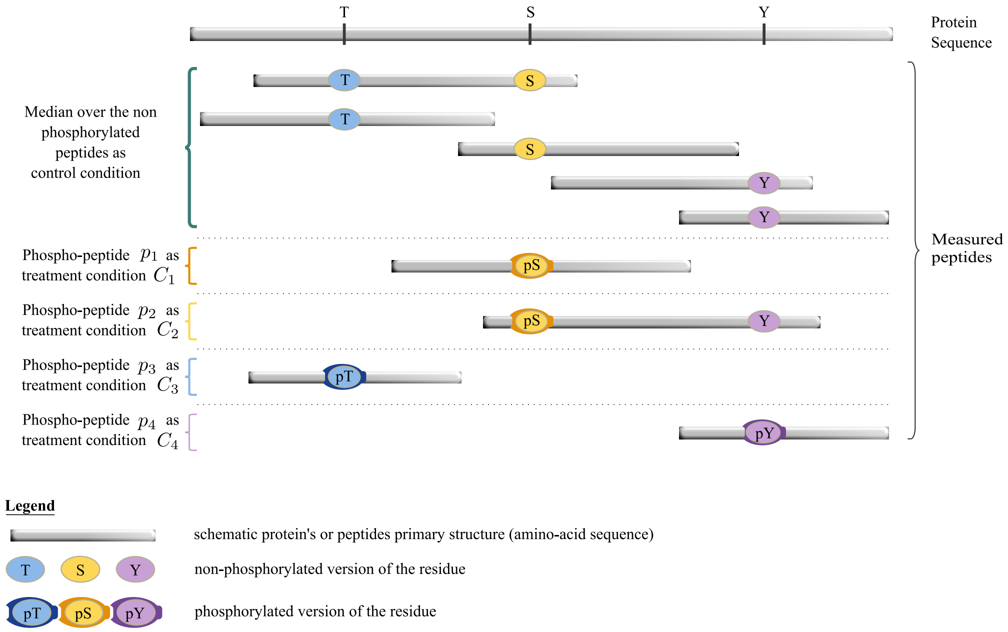


**Fig S18. Schematic visualisation of the phospho-TPP dataset [11] analysis.** We proceed to a reanalysis of the phospho-TPP dataset consisting in comparing the melting behaviours of phosphorylated peptides (denoted $p_1$ to $p_4$ in the figure) to the melting behaviour of the non-phosphorylated peptides associated to the same entry in the protein database (and forming the control condition). We illustrate the analysis using a schematic protein, presenting three phosphorylable residues (S,T and Y) along its sequence. Beside non-phophorylated peptides, four phospho-peptides are also observed. Replicates are not depicted in this figure. For each peptide, we require at least 2 replicates, and for each protein at least 3 non-phosphorylated peptides. Each phospho-peptide can be interpreted as a *treatement* condition, and the median observations computed over the non-phophorylated peptides are used as *control* condition. The five conditions are fitted simultaneously. In this analysis, we propose to use the median over the non-phophorylated peptides, as this is computationally efficient, while allowing to detect phospho-peptides with melting behaviours significantly different from the average melting behaviour of the *non-phosphorylated version* of the protein. We suggest that the use of the average melting behaviour deals with the case where different non-phophorylated peptides present various melting behaviours, for example due to the presence of isoforms with different melting behaviours. A more complete analysis, but out of the scope of this benchmarking, would be to compare the melting behaviour of any peptide (i.e. phosphoralyted or not) to the average melting behaviour of the protein.

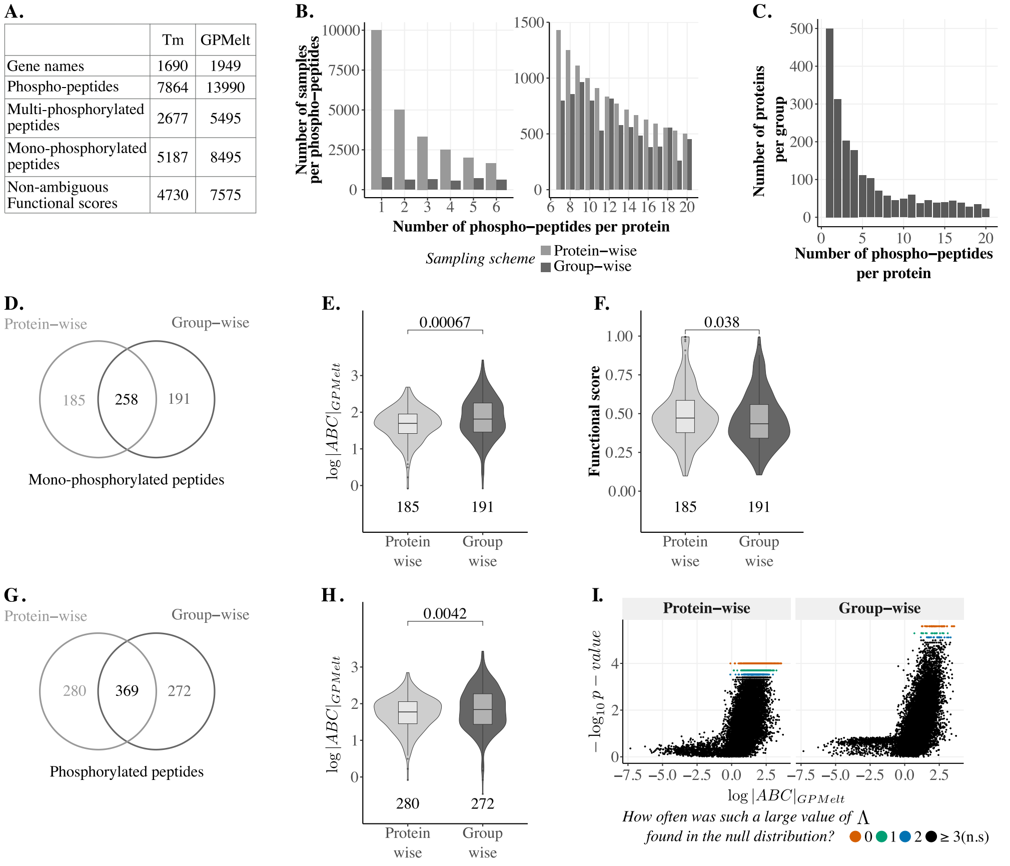


| | Tm | GPMelt |
|---|---|---|
| Gene names | 1690 | 1949 |
| Phospho-peptides | 7864 | 13990 |
| Multi-phosphorylated peptides | 2677 | 5495 |
| Mono-phosphorylated peptides | 5187 | 8495 |
| Non-ambiguous Functional scores | 4730 | 7575 |

**Fig S19. Additional results on the phospho-TPP dataset [11] analysis.** (A) An overview of the data entering the published $T_m$ analysis [11] versus the GPMelt analysis using a three-level HGP model. (B) Given a final size $S = 1e4$ of the null distribution approximation, the plot compares the required number of samples per phospho-peptide $\widetilde{S_p}$ (y-axis) to the number of phospho-peptide per protein $C_p$ (x-axis) for the *protein-wise* and *group-wise* approximation. The following relation holds: $\widetilde{S_p} = \left\lceil \frac{S_p}{C_p} \right\rceil$. In the case of the *protein-wise* approximation, $S_p \equiv S$ samples are obtained per protein. In the case of the *group-wise* null distribution approximation, $S_p = \left\lceil \frac{S}{N_g} \right\rceil$, with $N_g$ the size of the group the protein belongs to (depicted in panel (C)).(C) Size $N_g$ of the group $g$ for the *group-wise* approximation. The groups are defined by the set of proteins sharing the same number of phospho-peptides $C_p$. (D-F) **Results comparison for the mono-phosphorylated peptides.** (D) Venn diagram comparing the top 450 mono-phosphorylated peptides selected by each of the two null distribution approximations. (E) The absolute Area between the Curves $|ABC|_{GPMelt}$ for the mono-phosphorylated peptides uniquely selected by the protein-wise approach ($n = 185$) or the group-wise approach ($n = 191$) are compared (one-sided Wilcoxon signed-rank test). (F) Similarly, the functional score [33] of the mono-phosphorylated peptides uniquely selected by the protein-wise approach ($n = 185$) or the group-wise approach ($n = 191$) are compared (one-sided Wilcoxon signed-rank test). (G-I) **Results comparison considering all phosphorylated peptides.** (G) Venn diagram comparing the top 650 phospho-peptides selected by each of the two null distribution approximations. (H) Comparison of $|ABC|_{GPMelt}$ for the phospho-peptides uniquely selected by the protein-wise approach ($n = 280$) or the group-wise approach ($n = 272$) (one-sided Wilcoxon signed-rank test). (I) Volcano plot representing $\log|ABC|_{GPMelt}$ in function of $-\log_{10}(p - values)$. The colors indicates how extreme the values of $\Lambda$ are, compared to the values found in the null distribution approximation.

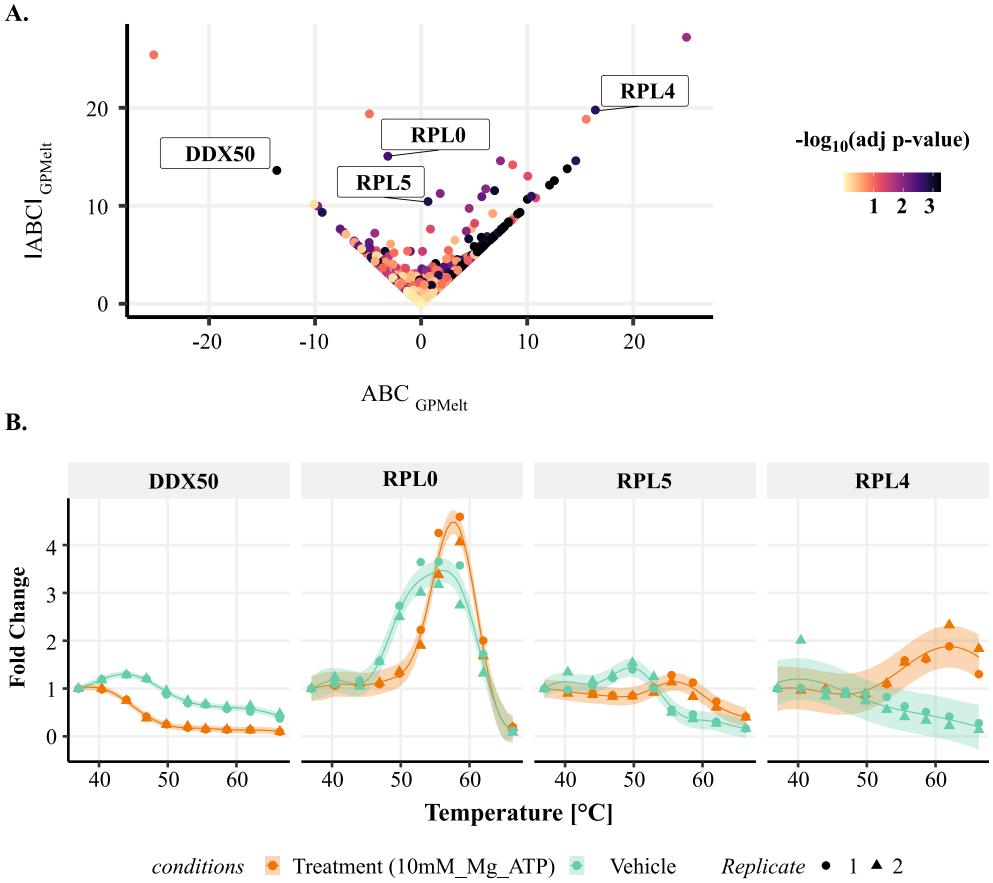


**Fig S20. ABC vs absolute ABC.** Results obtained from the ATP 2019 dataset [18](A) Comparing the Area Between the Curve $ABC_{GPMelt}$ to the absolute Area Between the Curve $|ABC|_{GPMelt}$. Proteins presenting small $ABC_{GPMelt}$ but large $|ABC|_{GPMelt}$ are proteins for which one curve will successively pass above and below the other one, with approximately similar amplitudes. For these proteins (e.g. RPL0, RPL5), there is not a unique/simple "stabilisation" or "destabilisation" effect (unlike the ones observed for DDX50 or RPL4).

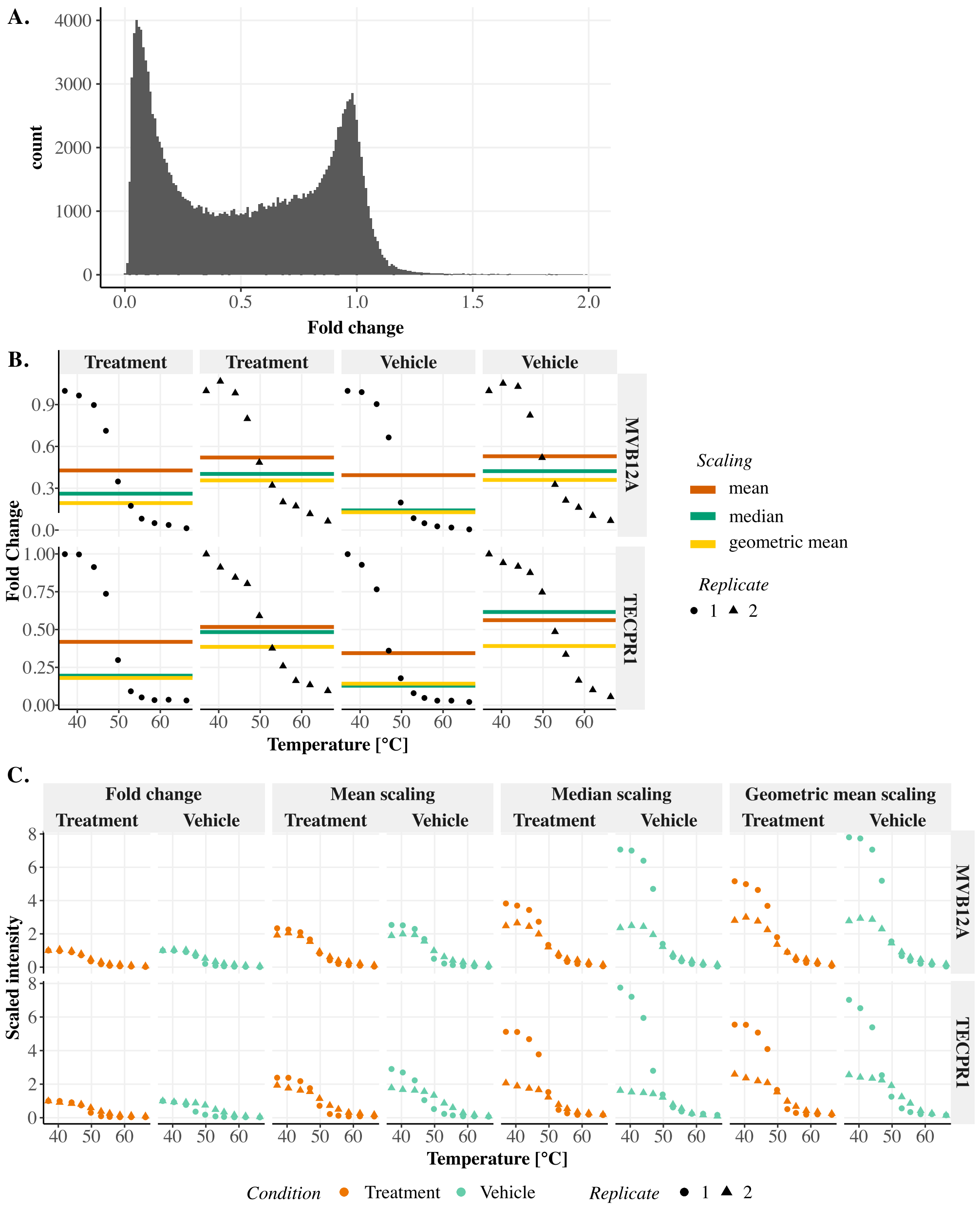


**Fig S21. The mean scaling is more appropriate than the median or geometric mean scaling when considering melting curves.** Plots are obtained using the ATP 2019 dataset [18] (A) Histogram of observed Fold Changes (FC). FC at $T = 37^\circ$ are not shown (because $\text{FC}(37^\circ) \equiv 1$). FC above two are also excluded for the sake of the visualisation (removing 143 out of 171792 values). (B) Example of two proteins illustrating the sensitivity of the median and geometric mean to the shape of the melting curve (especially to the steepness of the sigmoidal curve). For both proteins, the effect is particularly strong for the vehicle condition. The mean, median and geometric mean are computed for each replicate and represented by a colored line. (C) Effect of the FC, mean, median and geometric mean scalings on the shape of the replicates for the two proteins of panel (B). Due to the sensitivity of the median and geometric mean to the replicate shape, the resulting scaled replicates are very different within a condition.

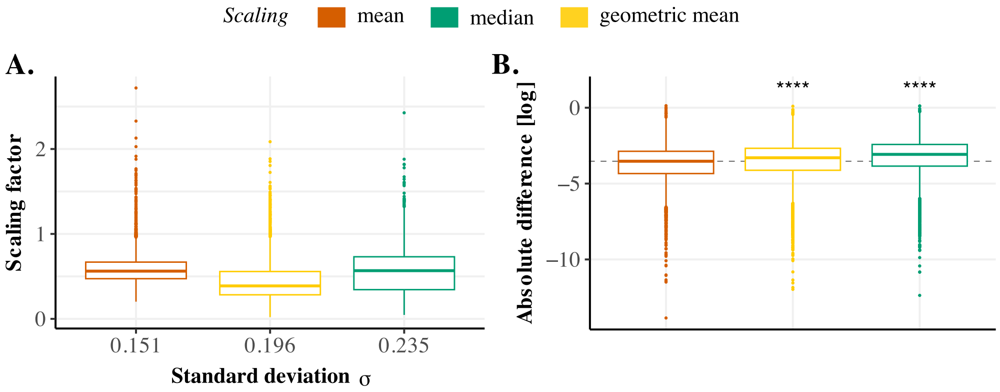


**Fig S22. The mean is more stable than the median or geometric mean when considering melting curve observations.** Plots are obtained using the ATP 2019 dataset [18]. (A) Boxplots of the computed mean, median and geometric mean for the different replicates of the different conditions of the ATP 2019 dataset [18]. The computed mean, median and geometric mean could be used as *scaling factors* for the replicates. Boxplots are ordered by their standard deviation $\sigma$. (B) Boxplot of the absolute difference in scaling factors within replicates of a condition (log scale). The one-sided Wilcoxon rank sum test rejects the null hypothesis in favor of the alternative hypothesis $\mathcal{H}_1$, with $\mathcal{H}_1$: using the mean scaling produces systematically smaller differences in scaling factors between replicates of a condition compared to the use of the median or geometric mean scaling (p-values $< 2.2e-16$).

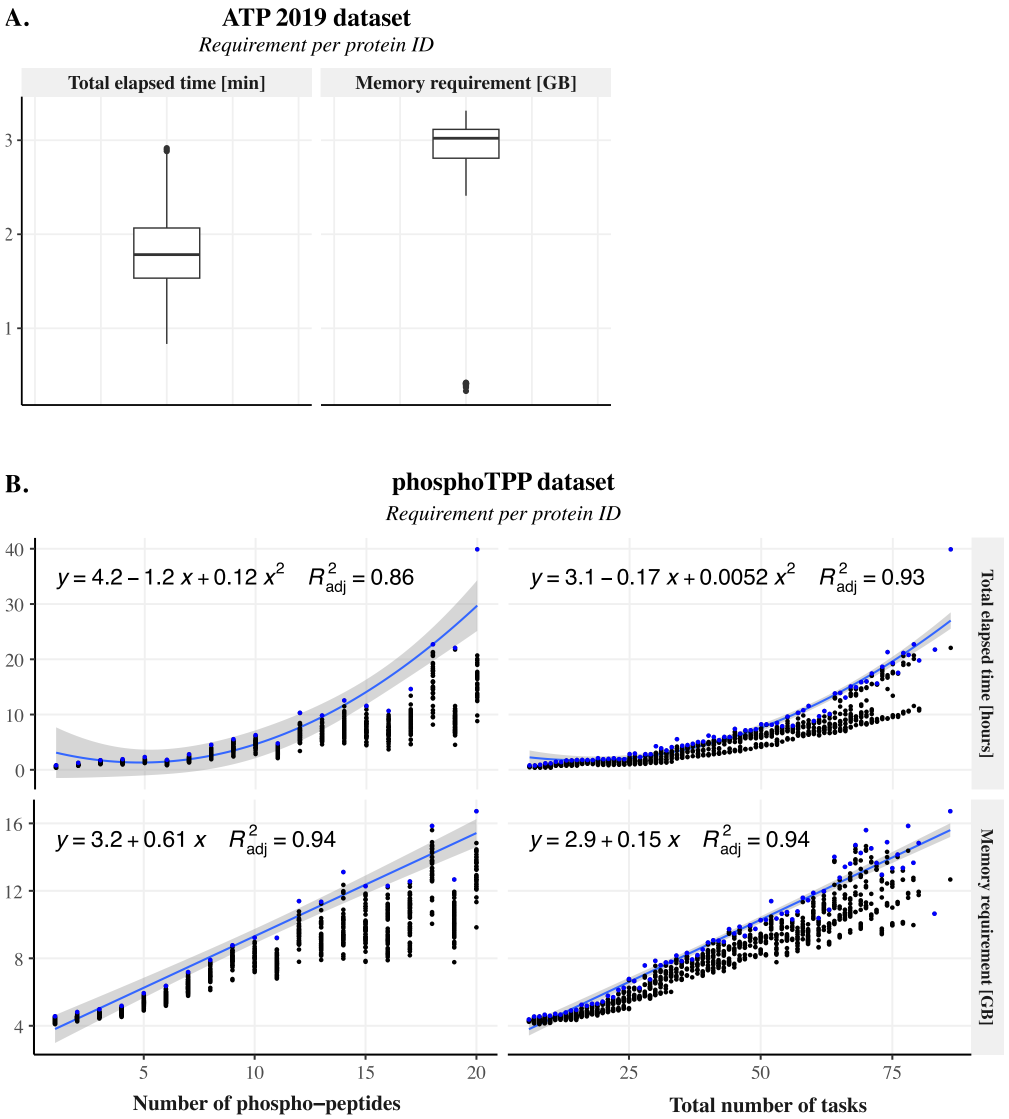


**Fig S23. Computational cost of the HGP model fitting and null distribution approximation for the ATP2019 and phosphoTPP datasets.** We consider here the running time of the algorithmic steps consisting in fitting the real data, sampling from the joint model $\mathcal{M}_0$ and fitting the model $\mathcal{M}_1$ via type II MLE to these samples. The p-value computation time is not included here and is negligible.**(A) A protein-level example.** Boxplot of the total running time (in minutes) and memory requirement (in GBs) for each protein ID of the ATP 2019 dataset [18]. For this dataset, 10 samples per ID are drawn from the joint model and fitted. Values for $N = 4772$ proteins ID are represented. **(B) A peptide-level example.** For this dataset, 10000 samples per ID are drawn from the joint model and consequently fitted. Values for $N = 2178$ proteins ID are represented. The first row represents the total running time (in hours) and the second row the memory requirement (in GBs) for each protein ID. The left column presents these values in function of the number of phospho-peptides of the protein, while the right column presents these values in function of the total number of tasks, i.e. the total number of replicates across all peptides. In blue are the largest values observed for each number of phospho-peptides, resp. for each total number of tasks. A regression line is fitted to these blue points, using the $R$ [46] (version 4.3.1) function *stat_regline_equation* from the ggpubr package [47] (version 0.6.0). The equation and $R^2$ values are indicated on the top left corner of each panel. This provides an upper bound for the time and memory requirement for a protein ID of this dataset, in function of the dataset size of this protein (total number of replicates or number of phospho-peptides). The code has been run on EMBL cluster.

| Number of treatment conditions | Method | Predicted distribution of the observations under the null hypothesis (used to draw the samples) | Number of samples per protein $S_p$ ($\widetilde{S_p} = \left\lceil \frac{S_p}{C_p} \right\rceil$) | Set of estimated parameters of the full HGP model $\mathcal{M}_1$ fitted to the sampled dataset of protein $p$ | Set of $\Lambda$ values obtained for the sampled dataset of protein $p$ | Null distribution approximation (final size $\geq S$) |
|---|---|---|---|---|---|---|
| one | A (*) | $\mathcal{N}(0, \widetilde{\Sigma}^{\mathcal{M}_0} + \beta_p^2 I_{N_p})$ | $S_p = \left\lceil \frac{S}{P} \right\rceil$ | $\{\theta^{MLE-II}_{p,s,\mathcal{M}_0}\}_{s \in [\![1,S_p]\!]}$ | $\{\Lambda^0_{p,s}\}_{s \in [\![1,S_p]\!]}$ | $\bigcup_p \{\Lambda^0_{p,s}\}_{s \in [\![1,S_p]\!]}$ |
| | B (**) | | $S_p = S$ | | | $\{\Lambda^0_{p,s}\}_{s \in [\![1,S_p]\!]}$ |
| $\geq 2$ $\{1, ..., C_p\}$ | C (*) | $\mathcal{N}(0, \widetilde{\Sigma}_p^{\mathcal{M}_{0,c_1}} + \beta_p^2 I_{N_p})$ ⋮ $\mathcal{N}(0, \widetilde{\Sigma}_p^{\mathcal{M}_{0,c_{C_p}}} + \beta_p^2 I_{N_p})$ | $\left\lceil C_p \cdot \frac{S}{\sum_{p'} C_{p'}} \right\rceil$ | $\{\theta^{MLE-II}_{p,s,\mathcal{M}_{0,c_1}}\}_{s \in [\![1,\widetilde{S_p}]\!]}$ ⋮ $\{\theta^{MLE-II}_{p,s,\mathcal{M}_{0,c_{C_p}}}\}_{s \in [\![1,\widetilde{S_p}]\!]}$ | $\{\Lambda^{0,c_1}_{p,s}\}_{s \in [\![1,\widetilde{S_p}]\!]}$ ⋮ $\{\Lambda^{0,c_{C_p}}_{p,s}\}_{s \in [\![1,\widetilde{S_p}]\!]}$ | $\bigcup_p \bigcup_c \{\Lambda^{0,c}_{p,s}\}_{s \in [\![1,\widetilde{S_p}]\!]}$ |
| | D (**) | | $S_p = S$ | | | $\bigcup_c \{\Lambda^{0,c}_{p,s}\}_{s \in [\![1,\widetilde{S_p}]\!]}$ |
| | E (***) | | $S_p = \left\lceil \frac{S}{N_g} \right\rceil$ | | | $\bigcup_{p \in g} \bigcup_c \{\Lambda^{0,c}_{p,s}\}_{s \in [\![1,\widetilde{S_p}]\!]}$ |

**Table S4. A summary of the different procedures proposed to approximate the null distribution of the statistic $\Lambda$.**
Notation: This table aims to explain the possible algorithms to approximate the distribution of the statistic $\Lambda$ (Eq (22)) under the null hypothesis. All the proposed methods (A to E) approximate this distribution by generating a set of at least $S$ values evaluated on $S$ samples generated under the null hypothesis. To describe these algorithms, we consider a protein $p$, with $p \in [\![1, P]\!]$, presenting $c$ treatment conditions, with $c \in [\![1, C_p]\!]$. For methods (A) and (B), $C_p \equiv 1$. For each protein, the set of all model parameters, denoted by $\theta_p$, is estimated via the fitting of the appropriate full HGP model $\mathcal{M}_1$. The model architecture (number of levels in the hierarchy, along with the number of independent lengthscales and output-scales) defines the exact content of $\theta_p$. Using the fitted model parameters $\theta_p^{MLE-II}$, the covariance matrix $\widetilde{\Sigma}_p^{\mathcal{M}_{0,c}}$ (Eq (57)) can be computed. This covariance matrix is used to define the distribution of the observations under the null hypothesis (third column). This distribution is further used to sample $S_p$ samples for protein $p$, with $S_p$ given by the fourth column. The value of $\theta^{MLE-II}_{p,s,\mathcal{M}_{0,c}}$ (fifth column) is obtained by fitting sample $s$, with $s \in [\![1, S_p]\!]$ or $s \in [\![1, \widetilde{S_p}]\!]$, for condition $c$ (i.e $s$ has been sampled under the null hypothesis that condition $c$ and the control condition are jointly modeled). Using this estimate, the statistic $\Lambda^{0,c}_{p,s}$ can be computed (sixth column). The final null distribution approximation of the statistic $\Lambda$ is defined by combining the values $\Lambda^{0,c}_{p,s}$, as described for each method in the last column: (*) dataset-wise, (**) protein-wise, (***) group-wise. In the last row of the table, we introduce the index $g$ to design a group of protein of size $N_g$.
**Method (A)**: A unique null distribution approximation is estimated per TPP-TR dataset. This is the simplest method to estimate the null distribution, and we typically use it for protein-level TPP-TR data with only two conditions (e.g. control and treatment). In this procedure, a small number $S_p$ of samples (typically 1 to few) is drawn for each protein. The null distribution of the statistic $\Lambda$ is then approximated by combining the computed null values $\Lambda^0$ for all samples of all proteins. **Method (B)**: Protein-specific null distribution approximation for TPP-TR data with only two conditions (e.g. control and treatment). The null distribution of the statistic is estimated per protein. This method is more rigorous but it is also more computationally expensive. In this case, $S$ samples are drawn and fitted for each protein. The null distribution of the statistic $\Lambda$ is approximated by combining the values of $\Lambda^0$ computed for each of the $S$ samples. **Methods (C-E)**: In the presence of multiple conditions, there exists multiple joint models. Namely, considering $C_p$ treatments and one control, $C_p$ joint models can be defined. We denote by $\mathcal{M}_{0,c}$ the model jointly modeling the control condition with treatment $c$, for $c \in [\![1, C_p]\!]$. **Method (C)**: Dataset null distribution estimation in the presence of multiple conditions. It consists in computing the same number of $\Lambda^0$ values for each comparison, and combining all values across proteins to form a unique null distribution. We only recommend this method if few conditions are available (typically 2 to 3), or if there is little amount of variation in the number of observed conditions across proteins. **Method (D)**: Protein-specific null distribution approximation in the presence of multiple conditions. This method is recommended for large number of conditions, especially if there is a large variation in the number of observed conditions and/or observations across proteins. For example, in the proposed analysis of the phospho-TPP dataset (see results section of the main text), proteins could be observed in one to twenty conditions, with each condition presenting three to five replicates (a condition corresponds to a phospho-peptide in this analysis). The protein-specific null distribution is obtained per protein by combining the values of $\Lambda^0$ obtained for all samples of all conditions of this protein. **Method (E) as a simplification of method (D)**: Group-wise null distribution approximation in the presence of multiple conditions. Similarly to method (D), the method (E) is recommended if there is a large variation in the number of observed conditions and/or observations across proteins, but when estimating a protein-specific null distribution is too expensive. In this case, we propose to group proteins (for example per actual number of observed conditions), and combine the protein-specific null distributions of all proteins belonging to a group, in order to reduce the number of samples per protein. Typically, if $S$ is the targeted size of the null dataset, and $N_g$ is the number of proteins in group $g$, then only $\left\lceil \frac{S}{N_g} \right\rceil$ instead of $S$ samples per protein in group $g$ are necessary. This method is exampled in S2 Supporting Information.
